# Time-resolved single-cell transcriptomics uncovers dynamics of cardiac neutrophil diversity in murine myocardial infarction

**DOI:** 10.1101/738005

**Authors:** Ehsan Vafadarnejad, Giuseppe Rizzo, Laura Krampert, Panagiota Arampatzi, Vallery Audy Nugroho, Dirk Schulz, Melanie Roesch, Paul Alayrac, Jose Vilar, Jean-Sébastien Silvestre, Alma Zernecke, Antoine-Emmanuel Saliba, Clément Cochain

**Affiliations:** Helmholtz Institute for RNA-based Infection Research (HIRI), Helmholtz-Center for Infection Research (HZI), 97080 Würzburg, Germany; Comprehensive Heart Failure Center Würzburg, University Hospital Würzburg, 97078 Wüzburg, Germany; Core Unit Systems Medicine, University of Würzburg, 97080 Würzburg, Germany; Institute of Experimental Biomedicine, University Hospital Würzburg, 97080 Würzburg, Germany; Institut National de la Santé et de la Recherche Médicale (INSERM), UMRS-970, Paris Centre de Recherche Cardiovasculaire, Université Paris Descartes, Sorbonne Paris Cité, 75015 Paris, France

## Abstract

**Objective:** After myocardial infarction, neutrophils rapidly and massively infiltrate the heart, where they can promote both tissue healing and damage. Here, we investigated the dynamics of cardiac neutrophil heterogeneity after infarction.

**Methods and results:** We employed single-cell transcriptomics (scRNA-seq) to investigate temporal neutrophil heterogeneity in the heart after murine myocardial infarction. At day 1, 3, and 5 after infarction, neutrophils could be delineated into six distinct clusters with specific time-dependent patterning and proportions. While the majority of neutrophils at day 1 were characterized by high expression of chemokines (e.g. *Cxcl3*, *Ccl6*), and putative activity of transcriptional regulators involved in hypoxic response (*Hif1a*) and emergency granulopoiesis (*Cebpb*), two major subsets of *Siglecf^hi^* (enriched for e.g. *Icam1* and *Tnf*) and *Siglecf^low^* (*Slpi, Ifitm1*) neutrophils were found at 3 and 5 days. Flow cytometry analysis confirmed the presence of LY6G^+^SIGLECF^hi^ and LY6G^+^SIGLECF^low^ neutrophils in the heart from 3 days after infarction onwards. LY6G^+^SIGLECF^hi^ neutrophils were absent from the bone marrow, blood and spleen, suggesting local acquisition of surface SIGLECF. Acquisition of the SIGLECF^hi^ state was paralleled by features of neutrophil ageing and activation (ICAM1^hi^CXCR4^hi^CD49d^hi^CD62L^low^). scRNA-seq of atherosclerotic aortas revealed two neutrophil subsets with gene expression patterns reminiscent of the major *Siglecf*^hi^ and *Siglecf*^low^ cardiac neutrophil subpopulations, revealing that these populations may be present across distinct contexts of cardiovascular inflammation.

**Conclusion:** Altogether, our data provide a time-resolved census of neutrophil diversity and gene expression dynamics in the mouse ischemic heart at the single-cell level, and suggests that temporal neutrophil heterogeneity is in part driven by local transition to a SIGLECF^hi^ state.

## Introduction

Neutrophils infiltrate the heart massively and rapidly after myocardial infarction (MI) (1),(2). Although long considered pathogenic, given their functional features (production of pro-inflammatory cytokines, neutrophil extracellular traps, proteases and reactive oxygen species) (3),(4), a recent report proposed a protective role of neutrophils via modulation of macrophage function (5), and several lines of evidence indicate that neutrophils may also have essential functions in tissue healing e.g. by promoting angiogenesis (6). Initially defined as a homogeneous population of sentinels constituting a first line of defense against invading microorganisms, recent research has highlighted the functional heterogeneity of neutrophils in mice and humans (7),(8),(9). How neutrophil functional heterogeneity is shaped by specific tissue niches, and in distinct pathological settings is still poorly understood (7),(8),(9). In MI, a process of temporal neutrophil polarization has been suggested, with N1 polarized pro-inflammatory neutrophils infiltrating the heart early after MI (day1), while at day 5 and 7 the proportion of N2 polarized anti-inflammatory neutrophils increases (10). However, the gene expression analysis performed in this report was limited to measurements of few selected markers commonly associated with *in vitro* M1/M2-polarized macrophages, an arbitrary dichotomy that may be of limited relevance to neutrophils in cardiac inflammation *in vivo* (11),(12). The recent development of single-cell RNA-sequencing (scRNA-seq) in high-throughput allows investigating immune cell transcriptional states in disease models with high resolution, and has recently been employed to uncover novel immune cell transcriptional states in e.g. myocardial infarction (13),(14) or atherosclerosis (15),(16). Furthermore, novel methods of oligonucleotide-barcoded antibody labeling of cells prior to their processing for scRNA-seq allows a multimodal measurement of the expression of cell surface epitopes in addition to transcripts expression (17),(18) and the multiplexing of several biological samples in a single scRNA-seq library (19). These technological advances improve the precision of immune cell subset identification, and allow comparing cells from different biological samples with minimal inter-sample technical bias and cellular doublets.

Here, we employed multiplexed time-series scRNA-seq analysis of cardiac myeloid cells to investigate the time-dependent heterogeneity of neutrophils in a murine model of MI. Combining transcriptomic analysis with flow cytometry validation experiments of cell surface protein expression levels, we describe the temporal diversity of neutrophil states in the infarcted heart, and show a local transition of neutrophils towards a SIGLECF^hi^ state within the ischemic cardiac tissue in mice.

## Results

### Time dependent transcriptional heterogeneity of neutrophils

Neutrophil infiltration in the infarcted mouse heart peaks around day 1 to 3 post-MI in mice, and then quickly resolves with only minimal neutrophils remaining at day 5 or 7 post-MI (Figure S1A, and (1), (2)). To analyze neutrophil transcriptional heterogeneity during the acute post-MI phase, we performed a multiplexed time series of scRNA-seq analyses combining cell surface epitope labeling and transcriptomics (cell hashing (17) and CITE-Seq (19), **methods**). MI was induced by permanent ligation of the left anterior descendant coronary artery in male C57BL6/J, and the timing of surgery was adjusted to minimize the impact of circadian oscillations on neutrophil gene expression patterns (20). Cell suspensions from the heart of mice at 1, 3, and 5 days after myocardial infarction were labeled with a viability dye and anti-CD11B antibodies coupled to distinct fluorochromes, and viable CD11B^+^ cells were sorted simultaneously (Figure 1A). Blood-borne neutrophils were excluded from analysis via i.v. injection of anti-CD45.2-APC before sacrifice (21). As expected, we observed two major populations of CD64^+^LY6G^−^ monocytes/macrophages and CD64^−^LY6G^+^ neutrophils that also expressed characteristic transcripts of monocytes/macrophages (*Csf1r*) and granulocytes (*Cxcr2*, *S100a8*), respectively (Figure 1B). The hashing antibody signal allowed backtracking the time point of origin of each single cell. We focused our analysis on cells corresponding to neutrophils (monocyte/macrophage data will be presented in another report), representing a total of 1405 individual neutrophils after excluding low quality cells and doublets (Figure 1B-C, and Figure S1B, **methods**), in which a median of 1352 expressed genes per cell were detected (Figure S1B). As previously observed by Zilionis et al. (22), the number of detected transcripts was much lower in neutrophils compared to other myeloid cells (we detected a median of 4260 genes/cell in monocyte/macrophages in the same dataset).

**Figure 1:**
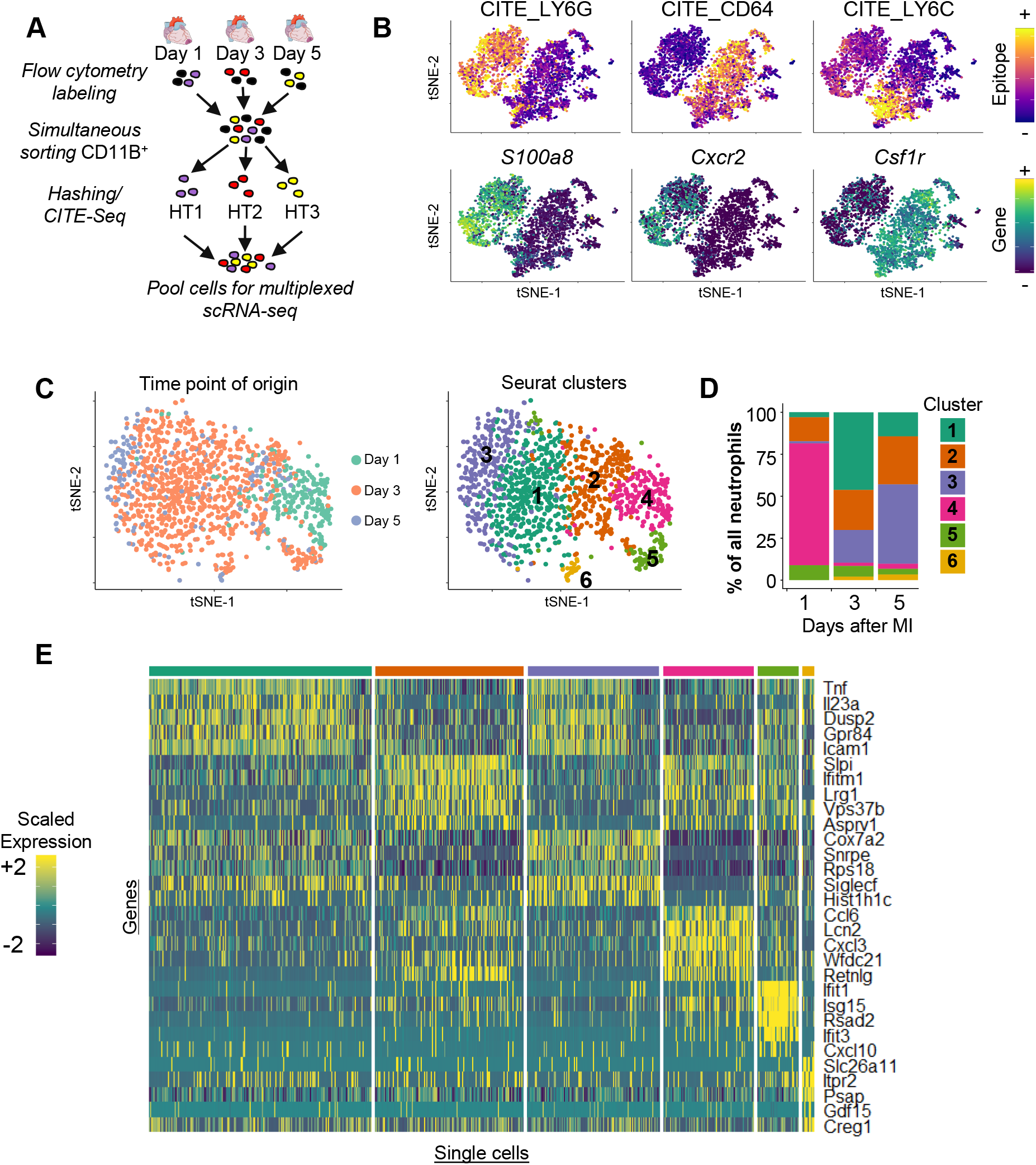
scRNA-seq reveals time-dependent neutrophil transcriptional heterogeneity after MI. **A)** Flowchart of the experimental design (HT: hashing antibody)**; B)** CITE-seq (Cellular Indexing of Transcriptome and Epitopes by sequencing) signal for the indicated surface markers and expression levels (log transformed expression) of the indicated transcripts projected onto the tSNE representation of scRNA-seq gene expression data of the total CD11B+ dataset; **C)** tSNE representation of gene expression data in 1405 neutrophils with time point of origin of individual cells color coded (left) and Seurat cluster assignment (right) projected onto the tSNE plot; **D)** proportion of each cluster among total neutrophils at each time point; **E)** heatmap of the top 5 marker genes (sorted by Log2 fold change) associated with the 6 neutrophil clusters (scaled expression).

In a first clustering analysis, we observed a substantial enrichment in genes encoding ribosomal proteins in some neutrophils. Expression of these transcripts appeared time-dependent, as their proportion was higher in cells originating from the day 3 and day 5 time points (Figure S1C). However, proportions were much lower in neutrophils compared to other cardiac CD11B^+^ cells at all time points (Figure S1D). Given the low likeliness of technical bias in our experimental design, and consistent findings in a separate experiment (see below), this differential expression of transcripts encoding ribosomal proteins most likely represents true biological variability, and could indicate increasing protein synthesis ability of neutrophils over time in the ischemic heart. However, to avoid clustering bias induced by preferential expression of ribosomal protein encoding genes, they were removed from the variable genes used to perform principal component analyses and clustering in Seurat (23),(24) (Figure 1C).

We compared gene expression patterns across all neutrophils at all time points and identified 6 transcriptionally distinct cell clusters (Figure 1C-E, **Supplementary Table**) that showed a time-dependent appearance in the ischemic heart (Figure 1C-D and S1E). As the number of cells sampled for each time point (day 1: 237 cells; day 3: 965 cells; day 5: 203 cells) did not correspond to the real levels of neutrophils in the ischemic heart over time (Figure S1A), we calculated the proportion that each cluster represented at each time point (Figure 1D). The vast majority of neutrophils at day 1 (72.6%) segregated in cluster 4 (*Cxcl3*, *Lcn2, Cd177, Ccl6, Sell, Fpr1*), a profile almost absent at later time points (<3% at day 3 and 5). At day 3, cluster 1 cells (46.2% of total neutrophils; *Tnf*, *Icam1*, *Il23a, Gpr84*) were predominant.

Cluster 2 cells (*Slpi*, *S100a11*, *Ifitm1, Lrg1*) were present at all time points and their levels gradually increased from 14.3% of cells at day 1 to 28.6% at day 5. Cluster 3 cells also increased to reach their highest level at day 5 (47.3% of all neutrophils; *Cox7a2*, *Hist1h1c*, *Siglecf*). Although cluster 1 and cluster 3 showed somewhat similar gene expression patterns and shared high expression levels of several transcripts (e.g. *Icam1*, *Siglecf*, *Dusp2*) compared to other neutrophils (Figure 1E), 78 genes showed significant differential expression between these 2 clusters (**Supplementary Table**), (e.g. *Cxcl2* or *Ptgs2* in cluster 1, *Ndufa11* or *Snrpe* in cluster 3). Cluster 5 represented a minor cluster (<10% of cells at all time points) with a strong type I interferon response signature (*Isg15*, *Rsad2*, *Ifit1*). Finally, cluster 6 cells were characterized by highly specific expression of some transcripts (*Fnip2*, *Cstb*, *Psap*, *Slc26a11, Gdf15, Pdcd1lg2, Mreg, Plgrkt*), and represented 0.0%, 2.1% and 3.4% of all neutrophils at day 1, 3 and 5 post-MI, respectively (Figure 1C-E). We further analyzed expression of some characteristic tissue dissociation-induced immediate early genes (IEGs) (*Egr1*, *Dusp1*, *Jun*, *Jund, Fos*, *Fosb*, *Hsp90aa1, Atf3)* (Figure S1F) (25),(26), but could not detect any clear pattern of expression of these IEGs across clusters, indicating that tissue-digestion induced changes in gene expression are unlikely to affect our clustering analysis. Altogether, these data indicate that 3 major time-dependent neutrophil states can be found in the ischemic heart (cluster 4 at day 1, cluster 1/3 and cluster 2 at day 3 and 5), in addition to two minor populations (the type I interferon response cluster 5, and cluster 6).

To validate our findings, we analyzed cells corresponding to neutrophils in an independent scRNA-seq experiment where total CD45^+^ cells were sorted from the heart before and at 1, 3, 5 and 7 days after infarction (Figure S2A). Consistent with the steady state heart containing only minute amounts of neutrophils, only 6 out of 522 cells in the control condition were annotated as neutrophils (we thus excluded the steady state condition from further analysis). The proportion of ribosomal proteins encoding genes in neutrophils increased in a time-dependent manner (Figure S2B-D). In this dataset, we could recover the major neutrophil states we had observed in the main dataset. A neutrophil cluster enriched for e.g. *Cd177*, *Cxcl3*, *Lcn2* and *Fpr1*, was found at day 1 post-MI (proximal to cluster 4 in the main dataset). At day 3 to 7, we observed a cluster enriched for *Siglecf*, *Icam1*, *Tnf* or *Dusp2* (proximal to cluster 1/3 in the main dataset), and a cluster enriched for e.g. *Ifitm1*, *S100a11* or *Slpi* (proximal to cluster 2 in the main dataset) (Figure S2E-G). We also reanalyzed a previously published scRNA-seq dataset of cardiac CD45^+^ cells collected 4 days after MI (27) (Figure S3A-B). Despite low transcript coverage (median genes detected/cell in total CD45^+^ cells=490; Figure S1B), we could detect two major populations of granulocytes: one was enriched for *Siglecf, Tnf, Icam1* and *Dusp2* expression, and the second for *Slpi, S100a11*, and *Ifitm1* (Figure S3C-D). Hence, these clusters respectively correspond to cluster 1/3 and cluster 2 cells observed at day 3 and 5 in our main dataset (Figure 1). Despite technical limitations (low numbers of cells and genes detected per cell (Figure S1B), possible variation caused by batch effects in pooled libraries), results obtained from these two additional datasets are consistent with conclusions obtained from our main dataset.

### SIGLECF^hi^ neutrophils time-dependently populate the infarcted heart

Neutrophils from day 3 and 5 post-MI (in particular cells from clusters 1 and 3) displayed a clear enrichment in the expression of *Siglecf* compared to expression at 1 day after MI (Figure 1 and 2A). We investigated the presence of SIGLECF^hi^ neutrophils in the heart after MI by flow cytometry. As the main switch in neutrophil *Siglecf* expression was evident between day 1 and 3 post-MI when neutrophils were the most abundant in the heart (Figure 2A and S1A), we focused our analysis on these time points. In enzymatically digested cardiac tissue, LY6G^+^SIGLECF^hi^ cells represented ~60% of LY6G^+^ neutrophils at day 3 post-MI, whereas almost all neutrophils were LY6G^+^SIGLECF^low^ at day 1 (Figure 2B-D and S4A). We also noted a population of LY6G^neg^SIGLECF^hi^ cells that were identified as eosinophils based on SSC/FSC properties (Figure 2B). Similar to LY6G^+^SIGLECF^low^ cells, the LY6G^+^SIGLECF^hi^ population was SSC^int^, supporting the notion that these cells are neutrophils (Figure 2B). No significant infiltration of LY6G^+^SIGLECF^hi^ neutrophils was found in the control heart or in the heart of sham operated mice at 1 and 3 days after surgery, as well as in the remote non-ischemic myocardium of mice at 1 and 3 days after MI (Figure S4B). As the detection of some neutrophil-relevant epitopes (e.g. CD62L, CXCR2) was affected by enzymatic processing, we also investigated the time-dependent presence of neutrophils in mechanically dissociated cardiac tissue, which yielded similar results, with a clear population of SSC^int^LY6G^+^SIGLECF^hi^ neutrophils observed at day 3 but not day 1 after myocardial infarction (Figure S4C). At day 5 after MI, SIGLECF^hi^ neutrophils represented half (51.01%) of total neutrophils (Figure S5A-C). Immunofluorescence analysis of MI tissue cryosections corroborated our flow cytometry analysis, with abundant SIGLECF^+^ cells found in the infarcted area at day 3, but not day 1, after MI (Figure 2E).

**Figure 2:**
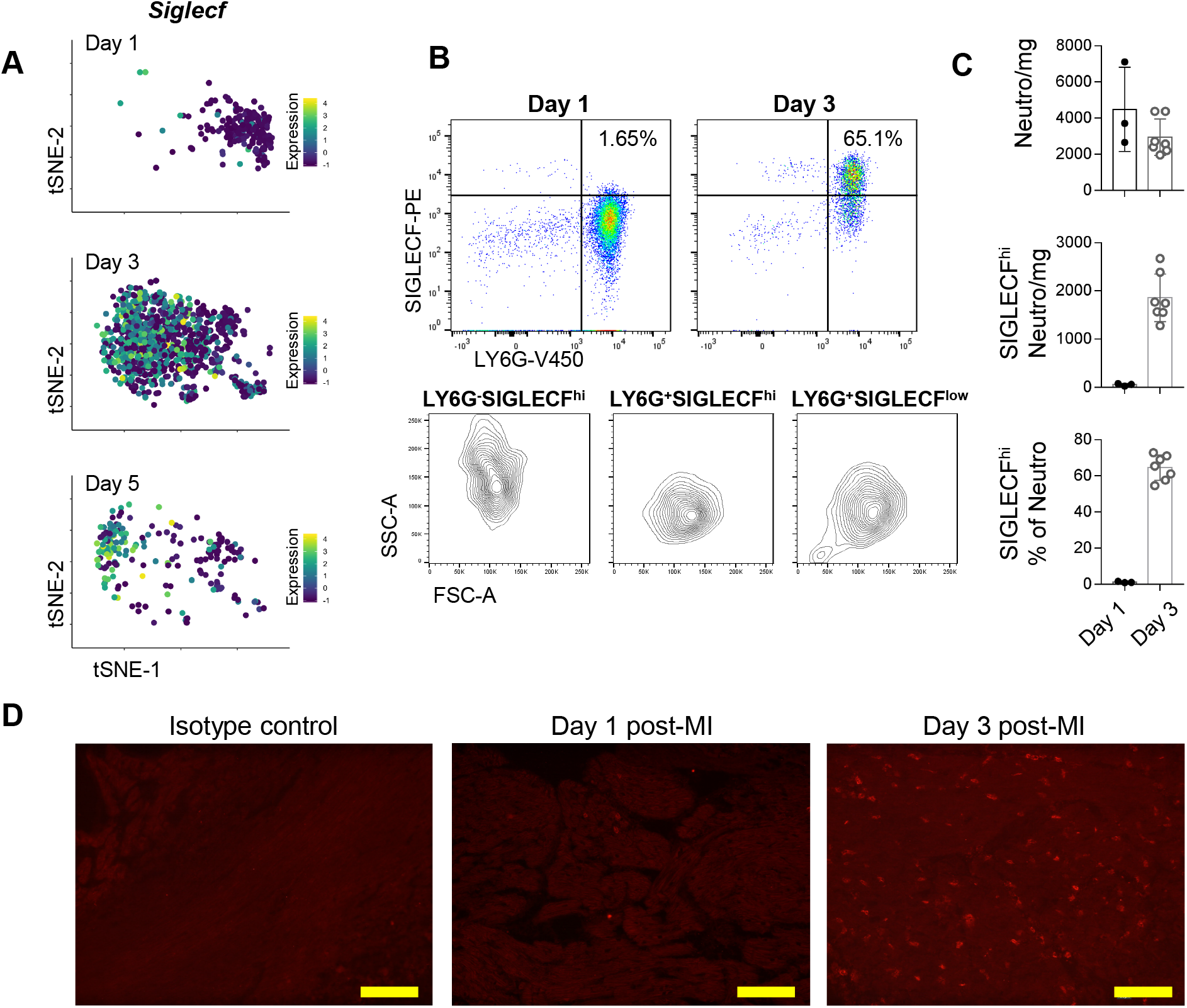
SIGLECF^hi^ neutrophils populate the heart at Day 3 post-MI. **A)** *Siglecf* scaled expression in single cells projected onto the tSNE plot (splitted according to time point of origin)**; B)** top: SIGLECF vs LY6G flow cytometry plots of cardiac cells gated on viable CD45^+^CD11B^+^CD64^low^ at 1 and 3 days after MI; bottom: FSC-A/SSC-A signal in gated LY6G^−^SIGLECF^hi^ eosinophils, LY6G^+^SIGLECF^hi^ and LY6G^+^SIGLECF^low^ neutrophils; **C)** neutrophils/mg, SIGLECF^hi^ neutrophils/mg, and proportion of SIGLECF^hi^ cells among LY6G^+^ neutrophils in the heart at 1 and 3 days after MI; **D)** immunofluorescence staining for SIGLECF in cryosections of hearts in the infarcted area at 1 and 3 days after MI, x200, scale bar 100µm (isotype control staining was performed on day 3 post-MI hearts).

### Acquisition of SIGLECF parallels acquisition of ageing and activation markers

Besides *Siglecf*, several markers previously associated with neutrophil ageing or activation (28), (29) showed differential transcript levels according to time (Figure 3A) or cluster (Figure S6A). The ageing/activation markers *Itga4* (encoding CD49d), *Icam1* (Intercellular Adhesion Molecule-1; ICAM1/CD54), and *Cxcr4* (C-X-C chemokine receptor type 4) levels increased over time, while the marker of “young” neutrophils *Sell* (encoding CD62L) decreased. *Cxcr2* (C-X-C chemokine receptor type 2) transcript levels decreased at the later time points. To investigate whether these changes in transcript levels were observable at the surface protein level, we measured expression of these markers in mechanically dissociated cardiac tissue neutrophils at 1 and 3 days after MI. We observed a clear biphasic expression pattern for ICAM1: at day 1 after MI, most neutrophils appeared negative for ICAM1 expression, in contrast to CD11B^+^LY6G^−^LY6C^hi^ monocytes which were uniformly positive (30) (Figure 3B). However, at day 3 post-MI, ~50% of neutrophils showed expression of ICAM1 (Figure 3B). Consistent with co-enrichment for *Siglecf* and *Icam1* in Cluster 1 and 3 (Figure 2A, 3A and S6A), ICAM1 expression was almost exclusive to SIGLECF^hi^ neutrophils (Figure 3C). Compared to total day 1 neutrophils and day 3 SIGLECF^low^ neutrophils, SIGLECF^hi^ neutrophils had higher CD49d and CXCR4, and lower CD62L surface expression (Figure 3D). We observed no significant difference in CXCR2 surface levels. At day 5, a similar pattern was observed for expression of ICAM1 and CD49d, with higher expression on SIGLECF^hi^ neutrophils (Figure S5D-E, expression of other epitopes was not measured in this experiment). We also evaluated the time dependent expression of transcripts encoding primary, secondary and tertiary granule proteins in the 6 neutrophil clusters (31), (Figure S6B). Consistent with previous transcriptomics analysis of mature neutrophils and their progenitors (32), transcripts encoding primary granule proteins (*Elane*, *Mpo*, *Ctsg*, *Prtn3*) were rarely found, while some genes encoding secondary and tertiary granule proteins were clearly enriched in cluster 4 (*Fpr1*, *Lcn2*, *Itgam*, *Mmp8, Hp*) (Figure S6B). Altogether, these data suggest that neutrophils in cluster 4, preponderant at day 1, may be transcriptionally more proximal to recently produced neutrophils and have features of “young” neutrophils (ICAM1^low^CD49d^low^CXCR4^low^CD62L^hi^), while at day 3 and 5, two major populations of SIGLECF^hi^ (ICAM1^hi^CD49d^hi^CXCR4^hi^CD62L^low^) and SIGLECF^low^ neutrophils (ICAM1^low^CD49d^low^CXCR4^low^CD62L^hi^) are found that likely correspond to cluster 1/3 and cluster 2, respectively.

**Figure 3:**
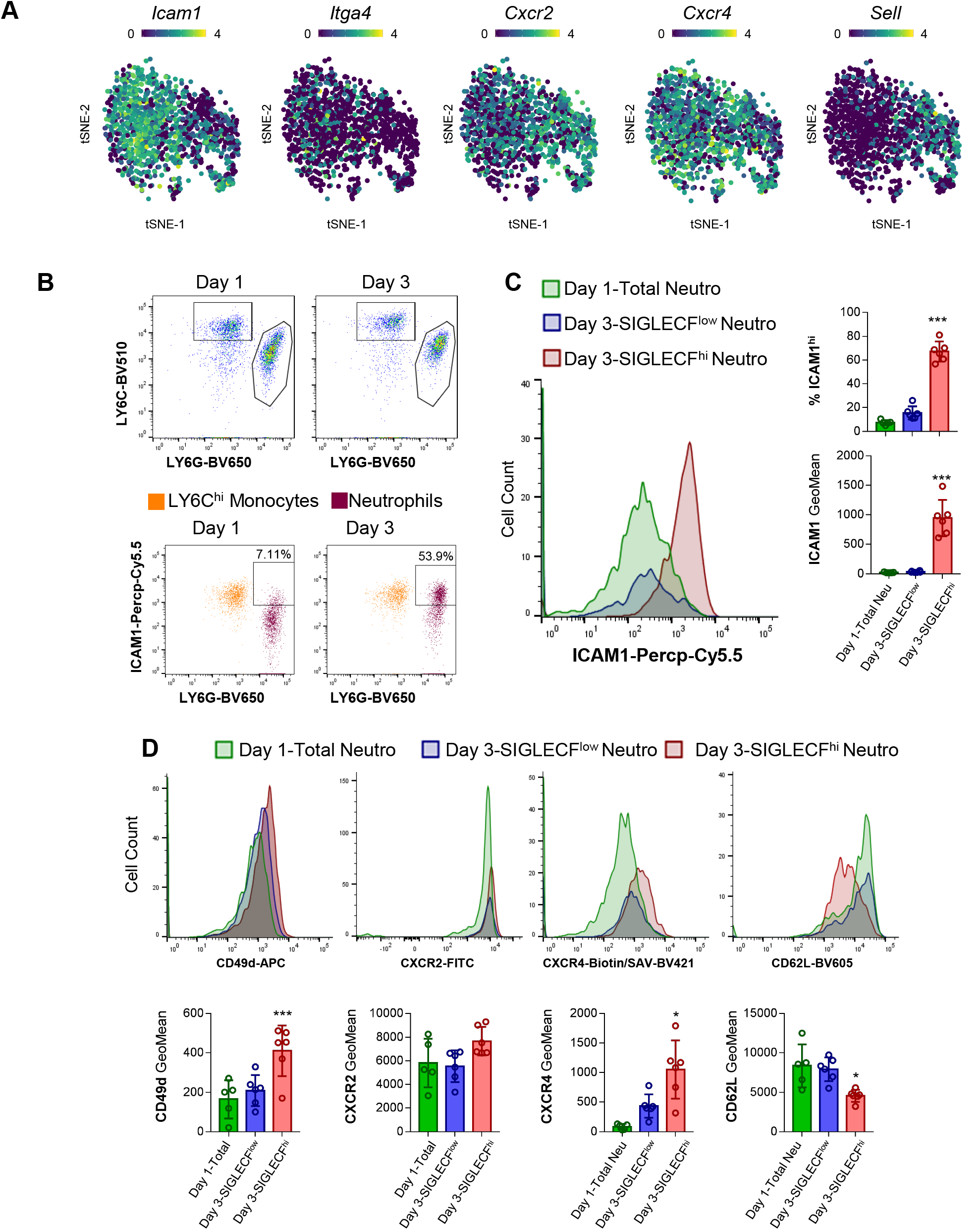
Time-dependent cardiac neutrophil subsets differentially express maturation and activation markers. **A**) Log normalized expression levels of the indicated transcripts projected onto the tSNE plot; **B)** top: gating of LY6G^+^LY6C^int^ neutrophils and LY6G^−^LY6C^hi^ monocytes in viable CD45+CD11B+ cells isolated from mechanically dissociated cardiac tissue; bottom: representative dot plots showing ICAM1 expression in LY6C^hi^ monocytes (orange) and neutrophils (purple) at Day 1 and Day 3 post-MI; **C)** representative histogram of ICAM1 expression and quantification of ICAM1+ cells and fluorescence intensity (Geometric Mean) in the indicated neutrophil subsets; **D)** representative histograms and quantification of fluorescence intensity (Geometric Mean) for the indicated epitopes on cardiac neutrophil subsets at day 1 (total neutrophils) and day 3 (SIGLECF^hi^ and SIGLECF^lo^ neutrophils). *p<0.05; **p<0.01; ***p<0.001

To investigate whether day 1 neutrophils and day 3/5 SIGLECF^hi^ and SIGLECF^low^ neutrophils represent various maturation stages originating from a similar precursor, and whether such potential transitions could be predicted from scRNA-seq data, we employed RNA-Velocity analysis (33). Although this revealed active transcriptional remodeling in many cells, potential transitions within clusters and from one state to another (e.g. from cluster 5 to cluster 4, or from cluster 2 to cluster 1), no clear time-dependent trajectory could be readily observed (Figure S6C).

### Neutrophils do not acquire SIGLECF and ICAM1 surface expression in the bone marrow, spleen or blood

We further evaluated whether SIGLECF^hi^ neutrophils were produced in the BM after MI, and transited via the blood circulation before infiltrating the heart. However, we could not detect any substantial population of SIGLECF^hi^ neutrophils in the blood, BM or spleen of MI and sham-operated mice at 1 and 3 days post-surgery (Figure 4A-B). Also after permeabilization and SIGLECF labeling we could not detect a SIGLECF^hi^ neutrophil population in the BM or blood in control mice and at 1 and 3 days after MI (Figure S7A). As we observed a clear co-expression of surface ICAM1 in cardiac SIGLECF^hi^ neutrophils, we also analyzed ICAM1 expression on BM, splenic and blood neutrophils, but did not detect any significant changes after MI (Figure S7B). Altogether, these data suggest that neutrophils acquire SIGLECF and ICAM1 surface expression in the heart and not at upstream production sites or in the circulation.

**Figure 4:**
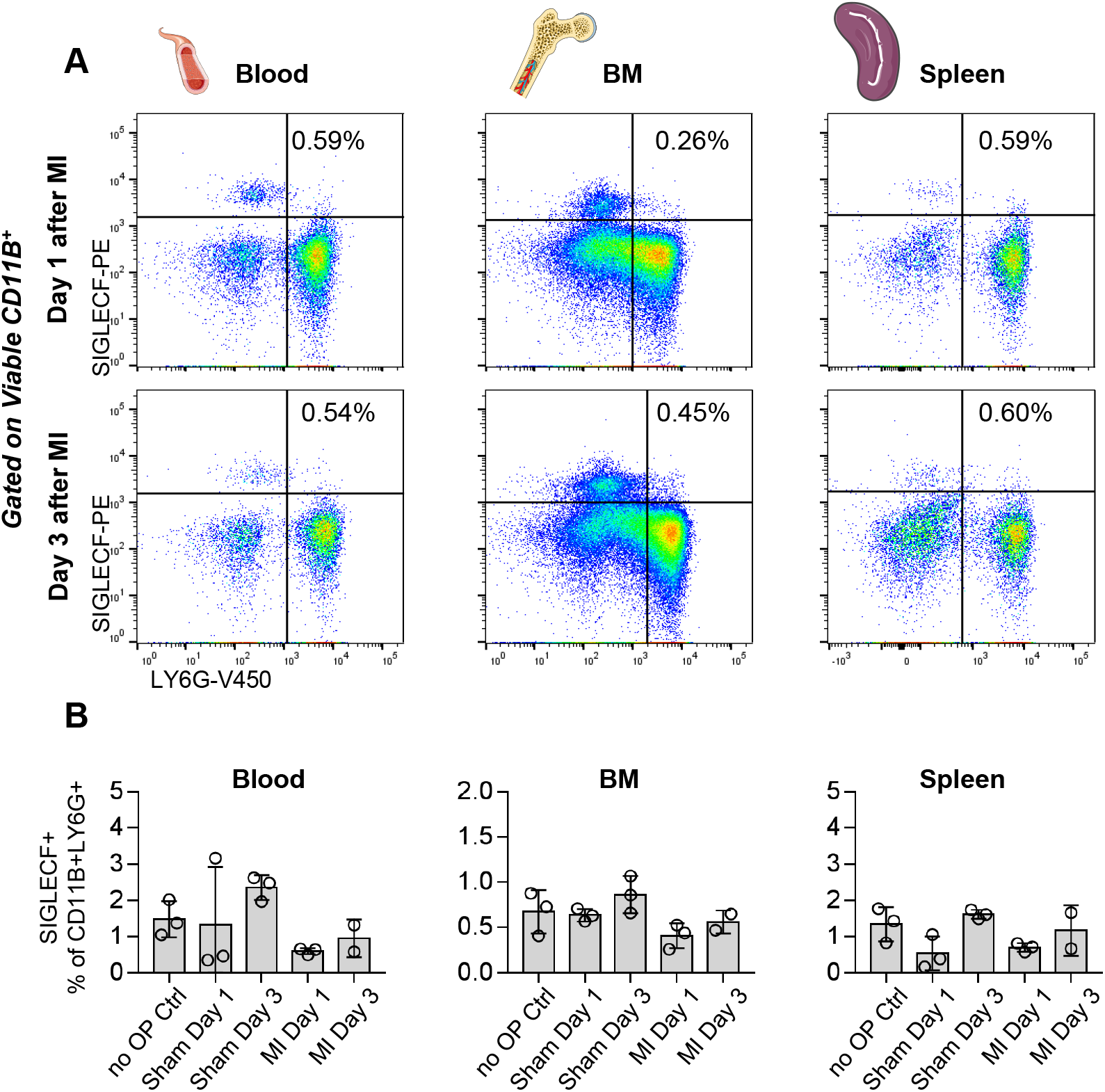
Circulating, bone marrow and splenic neutrophils do not acquire SIGLECF after MI. **A)** SIGLECF vs LY6G flow cytometry plots of cells from the indicated organs gated on viable CD45+CD11B+; **B)** proportion of SIGLECF+ neutrophils in the indicated organs in untouched mice, sham operated mice and MI-operated mice at 1 and 3 days after surgery.

### Single-cell regulatory network inference reveals putative regulators of neutrophil gene expression dynamics

We then sought to identify putative transcriptional regulators that may modulate gene expression in neutrophils in the infarcted heart and underlie their temporal heterogeneity. To this end, we employed single-cell regulatory network inference analysis (SCENIC) (34). SCENIC infers activity of gene regulatory networks (regulons) based on co-expression of transcription factors and their putative target genes (34). Analysis of regulon activity showed preferential activation of specific transcriptional regulators in the 6 neutrophil clusters (Figure 5A). In cluster 4, we observed preferential activity of regulons corresponding to e.g. *Hif1a, Snai1* and *Xbp1*, as well as *Cebpb*, a transcription factor with a critical role in emergency granulopoiesis(35). Regulons related to type I interferon response had enriched activity in Cluster 5 (*Irf7*, *Stat1*) (Figure 5A-B). Some regulons showed preferential activity in neutrophils from Cluster 1 and 3, e.g. *Maff* or *Creb5* (Figure 5A-B). Altogether, these data show putative cluster and time-dependent activation of gene expression regulatory networks in neutrophils in the ischemic heart.

**Figure 5:**
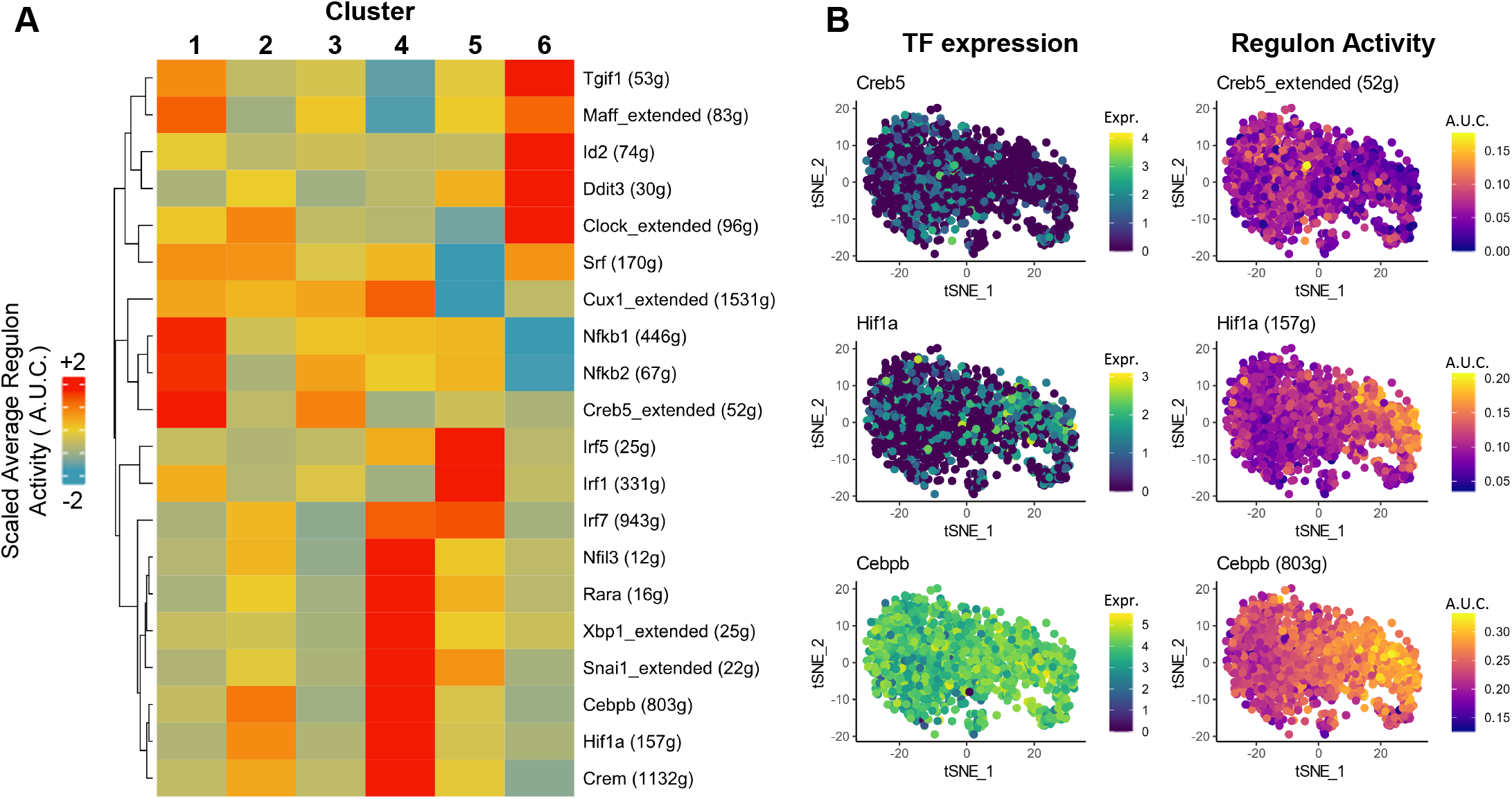
Single-cell regulatory network inference (SCENIC) analysis of neutrophils in the post-MI heart. **A)** heatmap of scaled average regulon activity (average area under the curve (A.U.C.) measured in SCENIC) in the 6 neutrophil clusters; **B) e**xpression of the indicated transcription factors (TF, log normalized expression) and activity of the corresponding regulon (ext: extended) in individual neutrophils as determined by SCENIC analysis and projected onto the tSNE plot representing the dataset (see **Figure 1**).

### scRNA-seq of atherosclerotic aortas reveals two distinct neutrophil subsets

To evaluate whether similar neutrophil diversity could be observed in a distinct cardiovascular disease context, we analyzed scRNA-seq data from 2106 CD45^+^ cells isolated from control (371 cells) and atherosclerotic (1735 cells) aortas of Low densitiy lipoprotein receptor deficient (*Ldlr-/-*) mice (corresponding to a previously published dataset combined with novel data, see ref (16), and **methods**) (Figure S8). Although only 66 cells corresponding to neutrophils were observed (Figure S8A), expression of the characteristic transcripts *Siglecf* and *Icam1* appeared to localize to a subpopulation with segregated tSNE coordinates (Figure S8B-D). This cluster was only observed in atherosclerotic aortas (Figure S8), and besides *Siglecf* and *Icam1*, these cells were enriched for the expression of *Tnf*, *Dusp2* or *Cxcr4*. The other neutrophil population (Figure S8B-D) was enriched for genes associated with cardiac *Siglecf^low^* neutrophils (i.e. cluster 2 and 4 in our main dataset) such as *Wfdc21*, *Cd177*, *Slpi*, *Hp* and had higher proportions of *Sell* exressing cells (Figure S8D). Despite low cell numbers in this dataset, these findings suggest that the major neutrophil dichotomy (i.e. *Siglecf^hi^* versus *Siglecf^low^*) may be also present in atherosclerotic aortas.

## Discussion

Here we demonstrate the time-dependent presence of diverse neutrophil states defined by discrete transcriptional and cell surface protein expression profiles in the infarcted mouse heart, and identify potential transcriptional regulators underlying this heterogeneity. Our observations are in contrast with the notion of a temporal polarization of pro-inflammatory N1 followed by anti-inflammatory N2 neutrophils (10). Although we found clearly different neutrophil states in the heart at day 1 post-MI versus day 3 onwards, our data indicate higher complexity in neutrophil states, with gene expression profiles that do not fall within the N1/N2 categorization, and do not allow assigning the various neutrophil clusters into subsets with distinctive pro or anti-inflammatory functions.

Our data suggest that temporal neutrophil heterogeneity is at least in part driven by a local neutrophil phenotypic transition within the ischemic heart. The clear bimodal expression of *Siglecf* transcripts and SIGLECF surface protein by neutrophils at day 1 versus day 3 onwards allowed us to track neutrophil state switch by flow cytometry. We observed expression of SIGLECF in cardiac neutrophils at day 3 and 5 post-MI, but could not detect SIGLECF^hi^ neutrophils in the blood, bone marrow or spleen, indicating that protein SIGLECF expression in neutrophils is acquired within the infarcted tissue. Similar results were observed for ICAM1 expression. In lung tumor bearing mice, abundant SIGLECF^hi^ neutrophils were found in tumors, while circulating neutrophils were SIGLECF^neg^ but had increased *Siglecf* mRNA levels, indicating that neutrophils may be primed to acquire SIGLECF or other specific gene expression patterns before infiltrating inflamed tissues (36). Thus, to what extent temporal neutrophil heterogeneity in the ischemic heart is driven by local processes versus priming at upstream production sites remains to be addressed. Future single-cell analyses of bone marrow and blood neutrophils before and after MI will likely help resolve this issue.

The functional implication of the local acquisition of surface SIGLECF and ICAM1 expression by murine neutrophils in the ischemic heart remains to be elucidated. As SIGLECF ligation has been proposed to induce apoptosis in eosinophils (37),(38) and its upregulation in neutrophils coincides with resolution of their infiltration in the ischemic heart, acquisition of SIGLECF expression in neutrophils could be associated with increased neutrophil apoptosis and would constitute an active process promoting the resolution of inflammation. Whether SIGLECF expression in murine neutrophils in disease states is of any relevance regarding neutrophil biology in other species is unclear. SIGLECF is considered a murine functional paralog of human Siglec-8 (39). In humans, Siglec-8 is expressed on circulating eosinophils, and to a lesser extent on mast cells and basophils (39),(40). However, based on sequence homology in their extracellular domain, SIGLECF was first proposed as an ortholog of human Siglec-5 (CD170), which is widely expressed by granulocytes, monocytes and other immune cells (40),(41),(42),(43). Regardless of these considerations, SIGLECF expression could constitute a valuable tool to track neutrophil diversity within diseased tissues in murine experimental models. ICAM1 expression can be induced by neutrophil activators (LPS, TNFα, zymosan), and can drive effector functions such as ROS production and phagocytosis, placing it as a functional marker of neutrophil activation (29). Whether local acquisition of surface ICAM1 on neutrophils has a functional role in the ischemic heart remains to be investigated. Neutrophil temporal heterogeneity in the heart was also associated with increased expression of transcripts that may orchestrate the inflammatory response. In particular, we observed increased expression of *Tnf*, which may act on numerous immune and non-immune cells (fibroblasts, cardiomyocytes, endothelial cells) in the ischemic heart, as well as genes that may mediate crosstalk of neutrophils with other immune cell types such as IL23R expressing T cells (*Il23a*), or monocyte/macrophages (*Csf1*) as previously proposed (5),(36).

In murine atherosclerotic aortas, we observed two broad neutrophil clusters with gene expression features reminiscent of the major cardiac clusters (i.e. *Siglecf^hi^* and *Siglecf^low^*), suggesting that these populations may be conserved in other cardiovascular inflammation contexts. The functional role of neutrophil heterogeneity in atherosclerosis, and potential tissue-specific aspects of neutrophil diversity will need to be addressed in future studies.

A recent scRNA-seq study in human and mouse lung cancer has evidenced conserved neutrophil transcriptomic signatures across species in disease (22). Notably, neutrophils with a strong type I interferon response, and neutrophils with enriched expression of a specific gene module (e.g. *Cstb*, *Fnip2*, *Ftl1*) reminiscent of cluster 6 in our study, were found across species. Although the cancerous lung and the ischemic myocardium represent two very distinct environments, the study by Zilionis et al. (22) provides proof of concept that disease-associated neutrophil gene expression signatures can be conserved from mice to humans, and raises the possibility that neutrophils similar to the ones we observed in our study may also populate the human ischemic heart. Elucidating the conservation of neutrophil states between the murine and human ischemic heart will be of critical importance to estimate the relevance of basic and pre-clinical research on acute inflammatory processes after MI in murine experimental models.

Major neutrophil functions (reactive oxygen species production, release of granule content, NET-osis, phagocytosis) may be largely independent of their transcriptional state (7). How these functions vary over time in infarct-infiltrating neutrophils requires further investigation. Also, how neutrophil temporal heterogeneity impacts repair of the infarcted heart remains to be determined. This will require precise, well controlled and validated timed depletion strategies (44). Gene expression manipulation using e.g. *S100a8^cre^* or *Ly6g^cre^* mice (45) may help to uncover the functional implications of time-dependent neutrophil gene expression. Neutrophil infiltration is already at its peak at our earliest analysis time point (24 hours), raising the possibility that additional neutrophil states may be found at earlier time points. Finally, we employed single-cell regulatory network inference analysis and RNA-velocity analyses to attempt uncovering transcriptional regulators and state transitions of neutrophils. However, events of gene expression regulation and transitions occurring at upstream sites may be transferred to the inflamed heart upon neutrophil recruitment. Thus, new methods able to capture the true immediate dynamics of transcription such as scSLAM-seq (46) may be better suited to investigate *in situ* phenotypic modulation of neutrophils in the ischemic heart.

In summary, our work provides a high-resolution census of dynamic neutrophil heterogeneity within the infarcted mouse heart. This dataset may constitute a valuable resource for further investigating the functional implications of neutrophil temporal heterogeneity in the infarcted myocardium, and how it may affect ischemic heart repair.

## Acknowledgements

The work was supported by the Interdisciplinary Center for Clinical Research (IZKF [Interdisziplinäres Zentrum für Klinische Forschung]), University Hospital Würzburg (Project IZKF-E-353 to C.C. and project Z-6 to P.A.), by the German Ministry of Research and Education within the Comprehensive Heart Failure Centre Würzburg (BMBF 01EO1504 to C.C. and A.-E.S.), the Deutsche Forschungsgemeinschaft (DFG, German Research Foundation) (Projektnummer 374031971 – TRR 240 to. A.Z).

## Data Availability

Datasets that were generated for this report have been deposited in Gene Expression Omnibus (GSE135310) and will be made freely available upon publication. The additional data of atherosclerotic aorta scRNA-seq have been deposited in Gene Expression Omnibus together with data from ref (16) (GSE97310).The previously published dataset from King et al. that was reanalyzed in this paper (Figure S3) (27) is available under GSE106473.

## Materials and methods

### Myocardial infarction

To induce myocardial infarction, 8 to 10 week old male C57BL6/J mice underwent permanent ligation of the left anterior descendent coronary artery. Mice were exposed to a 12/12 light cycle with lights on at 7:00 and lights off at 19:00. To minimize potential impacts of circadian rhythm on acute inflammatory processes, all surgeries were performed between 9:00 and 13:00. Briefly, mice received an injection of analgesic (Buprenorphin, 0.1mg/kg i.p.) and anesthesia was induced by isoflurane inhalation (4.0%). Mice under deep anesthesia (as revealed by absence of the paw withdrawal reflex) were intubated with an endotracheal cannula and placed under mechanical ventilation (CWE INC. SAR-830-AP; 130 respirations/min, peak pressure max. 18 cm H_2_O) on a heating pad to maintain body temperature. Anesthesia was maintained with 1.5-2.5% isoflurane during the surgery. A 4th intercostal thoracotomy was performed to expose the heart. The left descendent coronary artery was visualized an ligated with a 7/0 non-resorbable nylon suture. The thorax was closed with 3 separated 6/0 non-resorbable nylon sutures and the skin with a continuous 6/0 non-resorbable nylon suture. For pain relief, mice received i.p. injections of buprenorphine (0.1 mg/kg) 6 hours after the surgery and twice daily in the first two days after surgery. All animal studies and numbers of animals used conform to the Directive 2010/63/EU of the European Parliament and have been approved by the appropriate local authorities (Regierung von Unterfranken, Würzburg, Germany, Akt.-Z. 55.2-DMS-2532-2-743).

### Atherosclerosis

Diet-induced atherosclerosis was performed in *Ldlr^−/−^* mice as previously described in ref (16). After 10 weeks of high fat diet feeding, macroscopically visible plaques were mechanically removed from aortas. The plaque tissue (as well as the rest of the aortas) were processed for scRNA-seq as described in ref (16), generating two new datasets that were pooled with data from ref (16) in Seurat. All animal studies and numbers of animals used conform to the Directive 2010/63/EU of the European Parliament and have been approved by the appropriate local authorities (Regierung von Unterfranken, Würzburg, Germany, Akt.-Z. 55.2-DMS-2532-2-286).

### Cell isolation from hearts, sorting and processing for cell hashing/CITE-Seq

Mice were anesthetized with isoflurane (4.0%), received an intravenous injection of 5µg of anti-CD45.2-APC (clone 104, ThermoFisher Scientific) in 100µl of PBS, and left under isoflurane anesthesia for 5-10 minutes before being killed by cervical dislocation. We (16) and others (21) have previously established that this protocol leads to efficient labeling of all circulating leukocytes. Induction of myocardial infarction was macroscopically confirmed and mice received an intracardiac perfusion of PBS and the hearts were collected. The right ventricle was removed as well as the viable myocardium right above the ligature site, so that the infarcted area, its border zone and the adjacent viable myocardium were processed. A similar part of the left ventricle was processed for non-infarcted hearts. The excised hearts were rinsed in 4°C RPMI and gently massaged to expel excess blood, minced with surgery scissors and digested in RPMI containing 450U/ml collagenase I (Sigma-Aldrich C0130), 125U/ml collagenase XI (Sigma-Aldrich C7657), 60U/ml Hyaluronidase (Sigma-Aldrich H3506) for 1 hour at 37°C in a Thermomixer (Eppendorf) with shaking set at 1000rpm. The resulting cell suspension was filtered through a 70µm cell strainer, and incompletely digested pieces of myocardium were gently dissociated using a syringe plunger. Cells were then washed in 50ml 4°C PBS containing 1% Fetal Bovine Serum (centrifugation 8 minutes at 300rcf).

Cells were isolated from the heart of mice at 1 (n=5), 3 (n=5) and 5 days post-MI (n=3) as described above. Fc-Block was performed (10µg/ml purified rat anti-mouse CD16/CD32, Biolegend TruStain FcX anti-mouse), after which cells were stained with Fixable Viability Staining e780 (1:1000) and CD11B (M1/70) antibodies: CD11B-BV510 (Day 1 cells, 1:300, BD Biosciences), CD11B-Percp-Cy5.5 (Day 3 cells, 1:300, BD Biosciences) or CD11B-PE (Day 5 cells, 1:300, BD Biosciences) for 25 minutes at 4°C in PBS containing 1% FCS. Cells were washed once, resuspended in 1ml of PBS-1%FCS, pooled and centrifuged once more (8’, 350rcf). Viable cells labeled with CD11B-BV510, CD11B-Percp-Cy5.5 and CD11B-PE were separately sorted using a FACS Aria III (BD Biosciences) (See **methods** Figure 1). After sorting, cells were washed once in in 1ml of PBS-1%FCS (8’, 350rcf).Cells were then separately labeled with hashtag antibodies (Biolegend TotalSeq-A antibodies, Day 1: Hashtag 1, Day 3: Hashtag 2, Day 5: Hashtag 3) and CITE-Seq antibodies (Biolegend TotalSeq-A against mouse LY6C, LY6G, CD64, F4/80, CD11c, MSR1, IA/IE, TIM4, CX3CR1, and CCR2). Labeling was performed for 30 minutes at 4°C in 150µl PBS/1%FCS containing 1.5µg of each antibody. Cells were washed with 2ml PBS/1%FCS once (8’, 350rcf), resuspended in PBS supplemented with 0.04% BSA, pooled and centrifuged again (8’, 350rcf). The cells were resuspended in 150µl PBS/0.04% BSA, filtered through a 40µm cell strainer and counted before loading in the 10x Genomics Chromium.

**Methods Figure 1:**
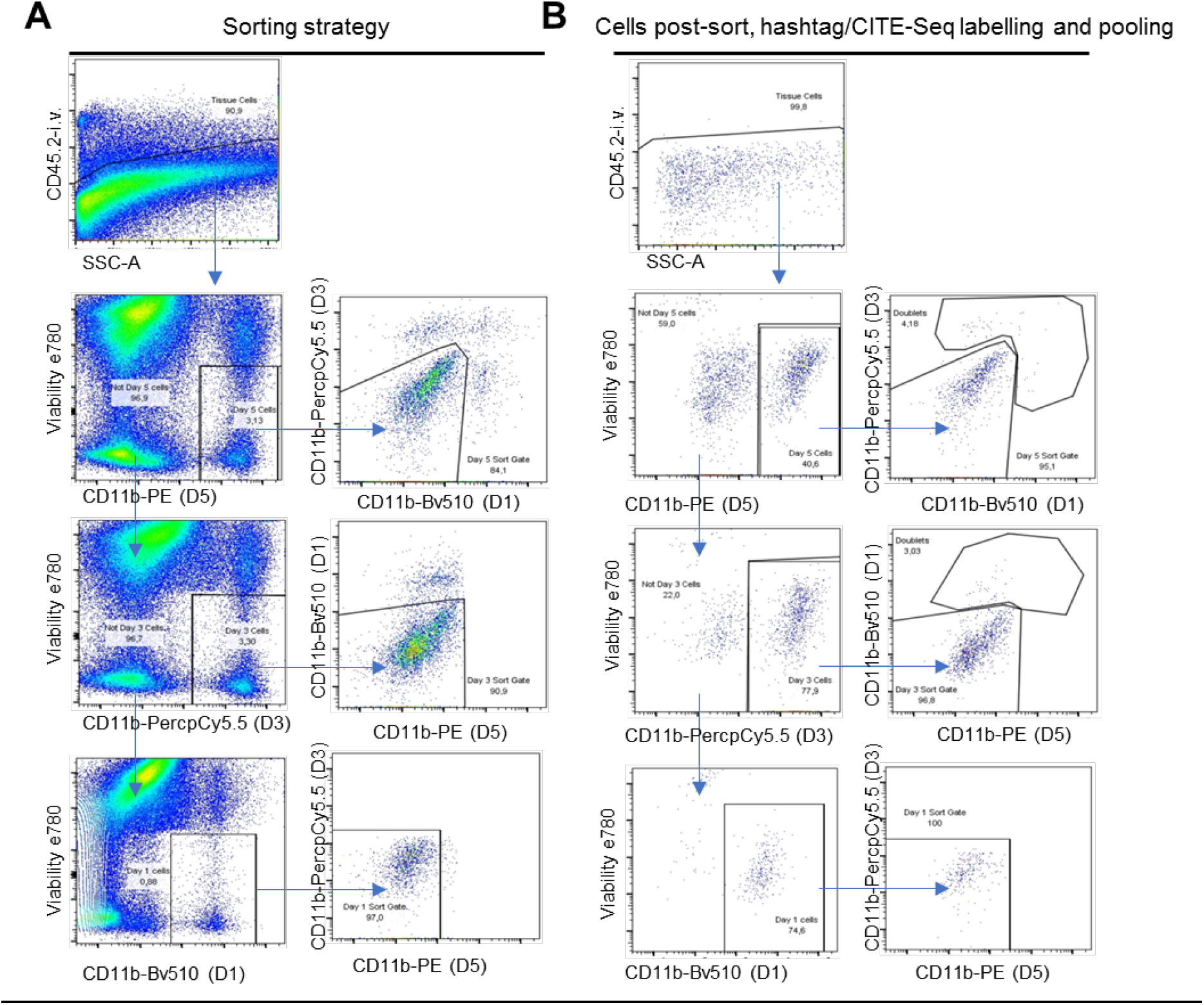
**A)** sorting strategy and **B)** purity of the cells loaded for scRNA-seq analysis for the multiplexing experiment at 1, 3 and 5 days post-MI

### Cell sorting: scRNA-seq kinetics day 0-1-3-5-7

Cardiac cells were collected at different time points after myocardial infarction (no infarct/Day 0: n=5; Day 1: n=5; Day 3: n=5, Day 5: n=5; Day 7: n=3) as described above. Cell suspensions were incubated for 10 minutes at 4°C with 10μg/ml Fc Block (purified rat anti-mouse CD16/CD32, Biolegend TruStain FcX anti-mouse) to block non-specific binding of antibodies to Fc Receptors and then incubated at 4°C for 30 minutes with anti-CD45-PE (1:300, Thermofisher Scientific, Clone 30-F11), anti-LY6G-V450 (1:300, BD Biosciences, Clone 1A8) and Fixable Viability Staining e780 (1:1000, Thermofisher Scientific). All staining and washing steps were performed in PBS containing 1% FCS, except the last wash and sorting that were made in PBS supplemented with 0.04% BSA (Sigma Aldrich, SRE0036). Around 10% of the cell suspension volume was transferred to flow cytometry tubes for analysis of neutrophil/monocyte/Mφ content in individual samples by flow cytometry analysis, while the rest of the cells for each time points were pooled for sorting. Right before sorting, cells were passed through a 40µm cell strainer. In this experiment, a neutrophil reduction step was performed to increase the proportion of monocyte/macrophages at early time points. Cells were sorted using a FACS-Aria III (BD Biosciences) with a 100µm nozzle, counted and loaded into the 10X Genomics Chromium.

### Mechanical dissociation of cardiac tissue for flow cytometry analysis

As we observed in pilot experiments that some epitopes (CXCR2, CD62L) were greatly affected by enzymatic digestion (not shown), we employed mechanical digestion of cardiac tissue in some experiments. Briefly, hearts were collected as described above, minced, and mechanically dissociated using a 12-Well Tissue Disaggregator plate (ScienceWare, Bel-Art Products). Cells were then washed and filtered using a 70µm cell strainer before being processed for flow cytometry labeling.

### Flow cytometry analysis

Cells were incubated for 10 minutes at 4°C with 10μg/ml of purified rat anti-mouse CD16/CD32 (Biolegend TruStain FcX anti-mouse) to block non-specific binding of antibodies to Fc Receptors and then incubated at 4°C for 30 minutes with Fixable Viability Staining e780 and fluorochrome/biotin conjugated antibodies (purchased from BD Bioscience, Biolegend, Thermofisher Scientific or Miltenyi biotec, and employed at concentrations recommended by the manufacturer) against mouse CD45 (clone 30-F11), anti-CD11B (M1/70), LY6C (HK1.4), LY6G (1A8), ICAM1 (YN1/1.7.4), SIGLECF (1RNM44N), CD49d (R1-2), CXCR4 (REA107), CXCR2 (SA044G4), CD62L (MEL-14), CD64 (X54-5/7.1). Specific fluorochromes conjugated to antibodies are indicated on flow cytometry panels in the figures. Data was acquired using a FACS Celesta (BD Biosciences) and analyzed with FlowJo.

### SIGLECF immunostaining

Hearts were perfused with PBS to removed excess blood, embedded in TissueTek OCT compound and snap frozen consecutively in liquid nitrogen cooled 2-Methylbutane and liquid nitrogen. Hearts were processed into 7µm cryosections (Leica CM3050 S) on SuperFrost Plus (Thermo Fisher Scientific) slides. Sections were incubated for 20 minutes with 10% normal goat serum to block nonspecific antibody binding, after which they were incubated for 2 hours with rat anti-mouse SIGLECF (Thermo Fisher Scientific, clone 1RNM44N, final concentration 10µg/ml). As a negative control, sections were incubated with the relevant isotype antibody at the same concentration (Rat IgG2a, kappa). After washing, sections were incubated with Alexa555 Goat anti-Rat IgG (1:300, Invitrogen) secondary antibody. Fluorescence microscopy imaging was performed with a Leica DM4000 B LED Fluorescence microscope and analyzed using the DISKUS software.

### Single-cell transcriptomics

Cells were loaded in the 10x Genomics Chromium at concentrations recommended by the manufacturer. Libraries were prepared with Chromium Single Cell 3’ Reagents Kit v2 Chemistry (day 0-1-3-5-7 kinetics dataset) or v3 Chemistry (main dataset: CITE-Seq/Hashing day 1-3-5 multiplexed post-MI dataset). The v2 chemistry samples were processed following the standard 10xGenomics protocol. The v3 sample was prepared for CITE-Seq/Hashing according to the manual until the cDNA amplification step in which 1 μl (0.2 μM) ADT PCR additive primer to capture Antibody-derived tags (ADTs) and 1 μl (0.1 μM) HTO PCR additive primer to capture Hashtag oligos (HTOs) were added. After cDNA amplification 60 μl (0.6x) SPRI beads (Beckman Coulter) were added to separate the supernatant fraction that contains the ADT/HTO-derived cDNAs (<180bp) and the bead fraction that contains the mRNA-derived cDNAs (>300bp). The mRNA-derived cDNAs (bead fraction) were processed following the standard 10xGenomics protocol. The ADT/HTO-derived cDNAs (supernatant fraction) were purified twice with 2x SPRI. After the two purifications half amount of the eluted cDNAs were amplified for ADTs and half for hastags. In the same reactions, the ADTs were indexed using TruSeq Small RNA primers and the hashtags using modified TruSeq DNA primers. The libraries were purified once more with 1.6x SPRI (160 μl). Ths method has been developed by Stoeckius et al., 2018 and is described in detail in reference (17). Additionally, the detailed protocol including the oligo sequences can be accessed here: https://cite-seq.com/protocol. All libraries were quantified by QubitTM 3.0 Fluometer (ThermoFisher) and quality was checked using 2100 Bioanalyzer with High Sensitivity DNA kit (Agilent). Sequencing was performed with S1 or S2 100bp flowcell with Novaseq 6000 platform (Illumina) and the reads for CITE-Seq/Hashing sample were allocated as follows: 5% for the hashtags, 10% for the ADTs and 85% for the mRNAs.

### Single-cell transcriptomics: data analysis

10X Genomics data was demultiplexed using Cell Ranger software (version 3.0.1) and HTO and ADT libraries were demultiplexed using a homemade script. Mouse GRCm38 reference genome was used for the alignment and counting steps. To evaluate the expression of the cell surface proteins alongside the transcriptome level in our main data set, --feature-ref flag of Cell Ranger software was used which creates a matrix that contains gene expression counts alongside the expression of cell surface proteins. The gene-barcode matrix obtained from Cell Ranger was further analyzed using Seurat (version 2.3.4). For the day 0-1-3-5-7 kinetics dataset, data obtained from each time point was integrated using Cell Ranger. To reduce the batch effect introduced by different sequencing depth, the read depth was equalized between libraries before integrating them.

In the main dataset (including cell hashing (17) and CITE-seq (19)), quality control was performed as follows: cells with less than 500 or more than 7500 detected genes per cell were filtered out. Moreover, cells with more than 5% UMIs mapped to mitochondrial genes were filtered out as well. The transcriptome of remaining cells was normalized and log transformed and highly variable genes were identified using the default settings of Seurat. Dimension reduction and clustering analysis were performed using the first 20 principal components. Clustering analysis was performed on gene expression data alone, at which point the cell hashing and CITE-seq signal were added to the data. A time point identifier was appended to each individual cell based on Hashtag antibody signal. Cells with positive signal for two hashtag antibodies, and outlier cells with extreme Hashtag antibody signal were discarded.

The major myeloid cell populations (i.e. monocyte/macrophages, dendritic cells, NK cells and neutrophils) were identified based on expression of canonical transcripts and CITE-seq signal of known surface markers, and cells corresponding to neutrophils were subset into a new Seurat object (see main text), where highly variable genes were identified using the FindVariableGenes function. As a preliminary analysis indicated a strong enrichment of ribosomal protein encoding genes (starting with Rpl- or Rps-) in some neutrophils, these genes were removed from the variable gene list before performing principal component analysis (PCA). Clustering was performed at a 0.8 resolution using the 15 first principal components. Markers genes were identified using the FindAllMarkers function in Seurat with default settings.

### SCENIC analysis

SCENIC was applied on the main dataset as described in (34). For the analysis, mm9 motif ranking database supplied in SCENIC package was used to score potential regulons. The input matrix genes that were not present in motif ranking database were excluded from further analysis. Target genes that did not show a positive correlation based on the GENIE3 co-expression algorithm (> 0.03) in each module were filtered out. TF-modules having less than 20 genes were filtered out and the remaining TF-modules were further examined.

### Velocity analysis

To count the spliced and unspliced UMIs in our data set, the velocyto package (33) was used. To be able to infer the future state of cells in our data set, we employed scvelo (47). Normalized and log transformed data was used to calculate first and second order moments for each cell across its nearest neighbors. Next, the velocities were estimated and the velocity graph constructed using the scvelo.tl.velocity() with the mode set to ‘stochastic’ and scvelo.tl.velocity_graph() functions. Velocities were visualized on top of the previously calculated tSNE coordinates using Seurat.

---

**Figure S1:**
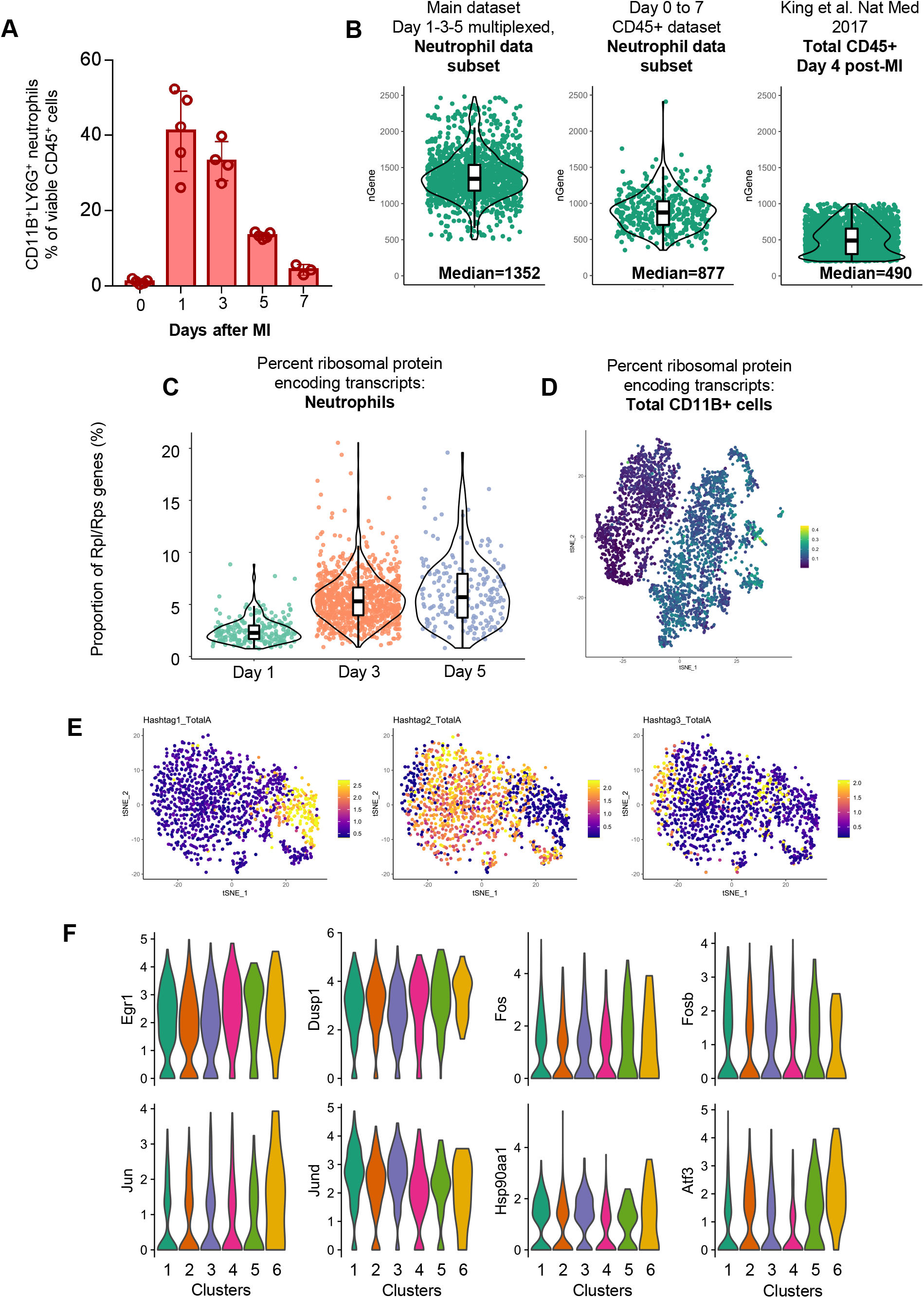
General data and scRNA-seq metrics. **A)** Proportion of CD11B^+^LY6G^+^ neutrophils among total cardiac CD45+ leukocytes at 0 (control heart) and 1, 3, 5 and 7 days after myocardial infarction (MI); **B)** violin plot indicating the number of genes detected in each cell in the datasets analyzed in this report; **C)** percent of transcripts encoding ribosomal proteins in neutrophils according to their time point of origin; **D)** percent of transcripts encoding ribosomal proteins (name starting with Rps- or Rpl-) projected onto the tSNE plot of the global CD11B+ cells dataset shows higher proportion of these transcripts in non-neutrophil CD11B+ cells; **E)** hashtag antibody signal used to determine time point of origin of individual neutrophils (for clarity, minimum and maximum expression cutoffs have been applied); **F)** violin plot showing log normalized expression of characteristic tissue dissociation induced immediate early gene in neutrophil clusters.

**Figure S2:**
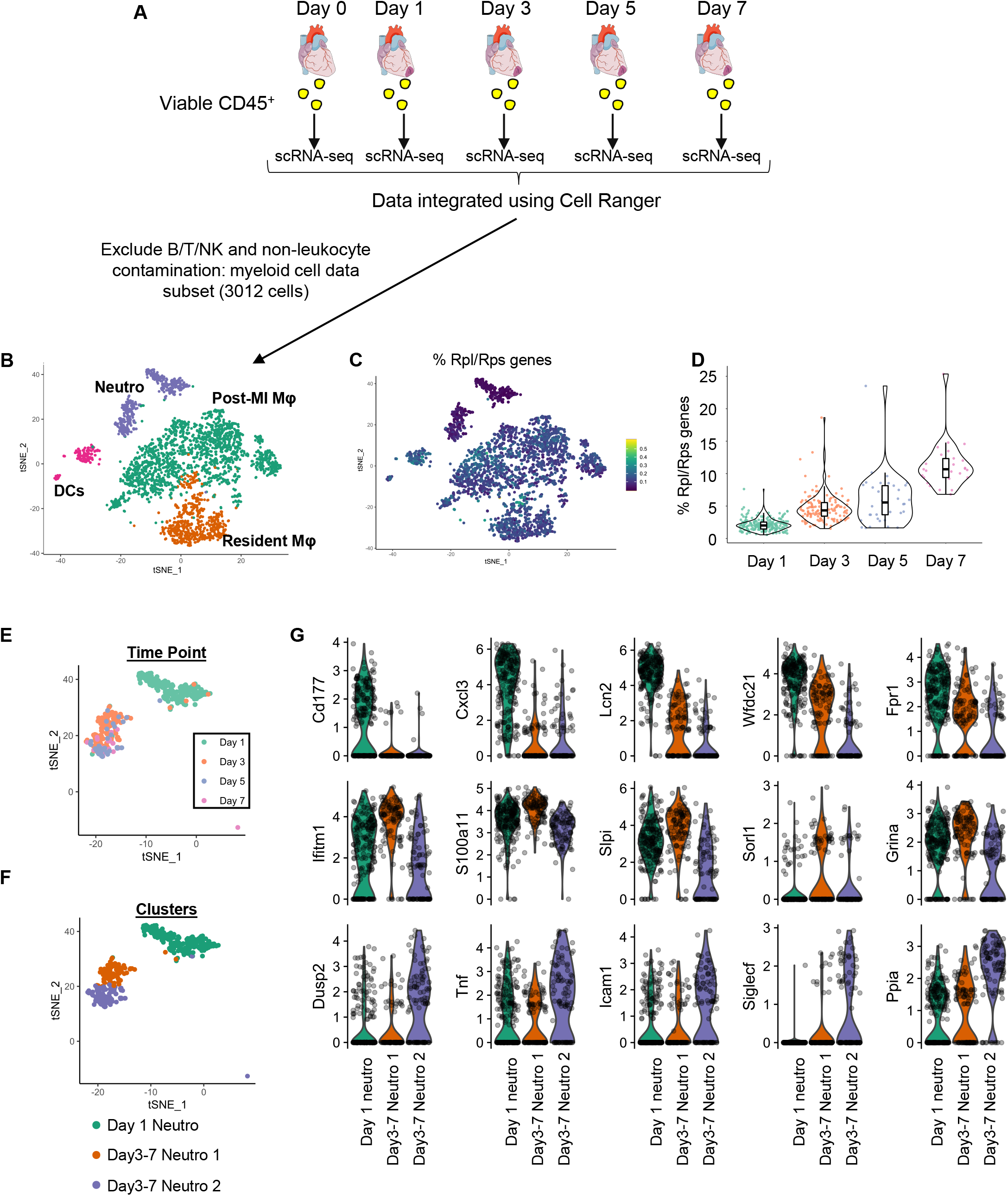
Analysis of time series scRNA-seq of total cardiac CD45+ cells, day 0 to 7 post-MI. **A)** Summary of the experimental design; **B)** identification of major myeloid cell lineages in the dataset projected onte the tSNE plot; **C)** Percent of transcripts encoding ribosomal proteins projected onto the tSNE plot of total myeloid cells; **D)** violin plot showing percent of transcripts encoding ribosomal proteins in neutrophils, according to time point of origin; **E)** time point of origin of single neutrophils projected onto the tSNE plot; **F)** identification of neutrophil clusters on the tSNE plot; **G)** violin plot showing log normalized expression of the indicated transcripts in neutrophils, according to cluster.

**Figure S3:**
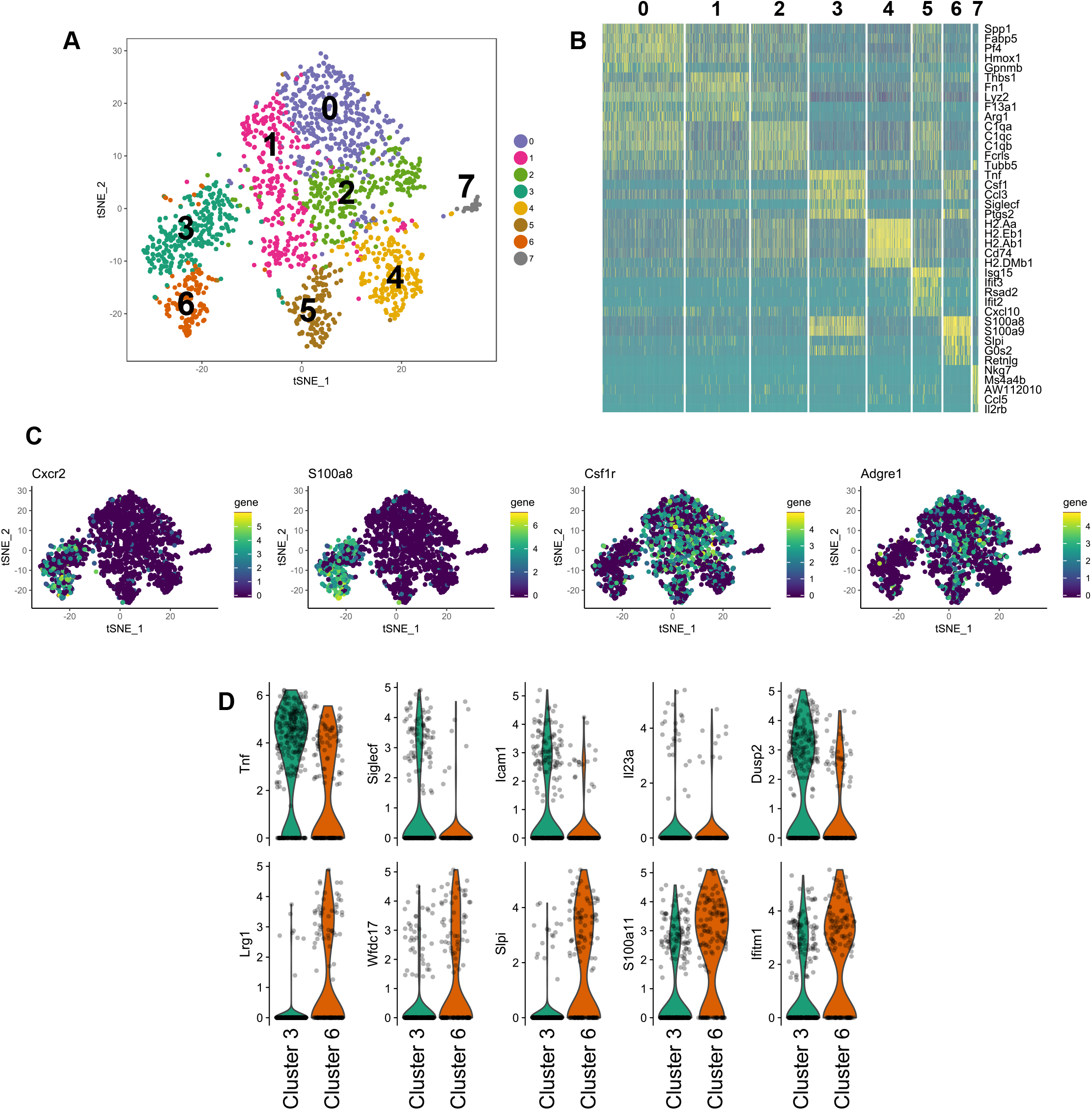
Reanalysis of King et al., scRNA-seq of CD45+ cells at day 4 after MI (King et al. Nat Med 2017). **A)** tSNE representation of scRNA-seq gene expression data and clustering analysis; **B)** heatmap of the top 5 marker genes (ordered by Log2 fold change) in each cluster; **C)** expression of the indicated markers identifies cells corresponding to neutrophils (*S100a8+*, *Cxcr2+*, no expression of *Adgre1* or *Csf1r*) and **D)** expression of the indicated genes in the two neutrophil clusters (Cluster 3 and 6 on the tSNE plot/heatmap, panel **A**).

**Figure S4:**
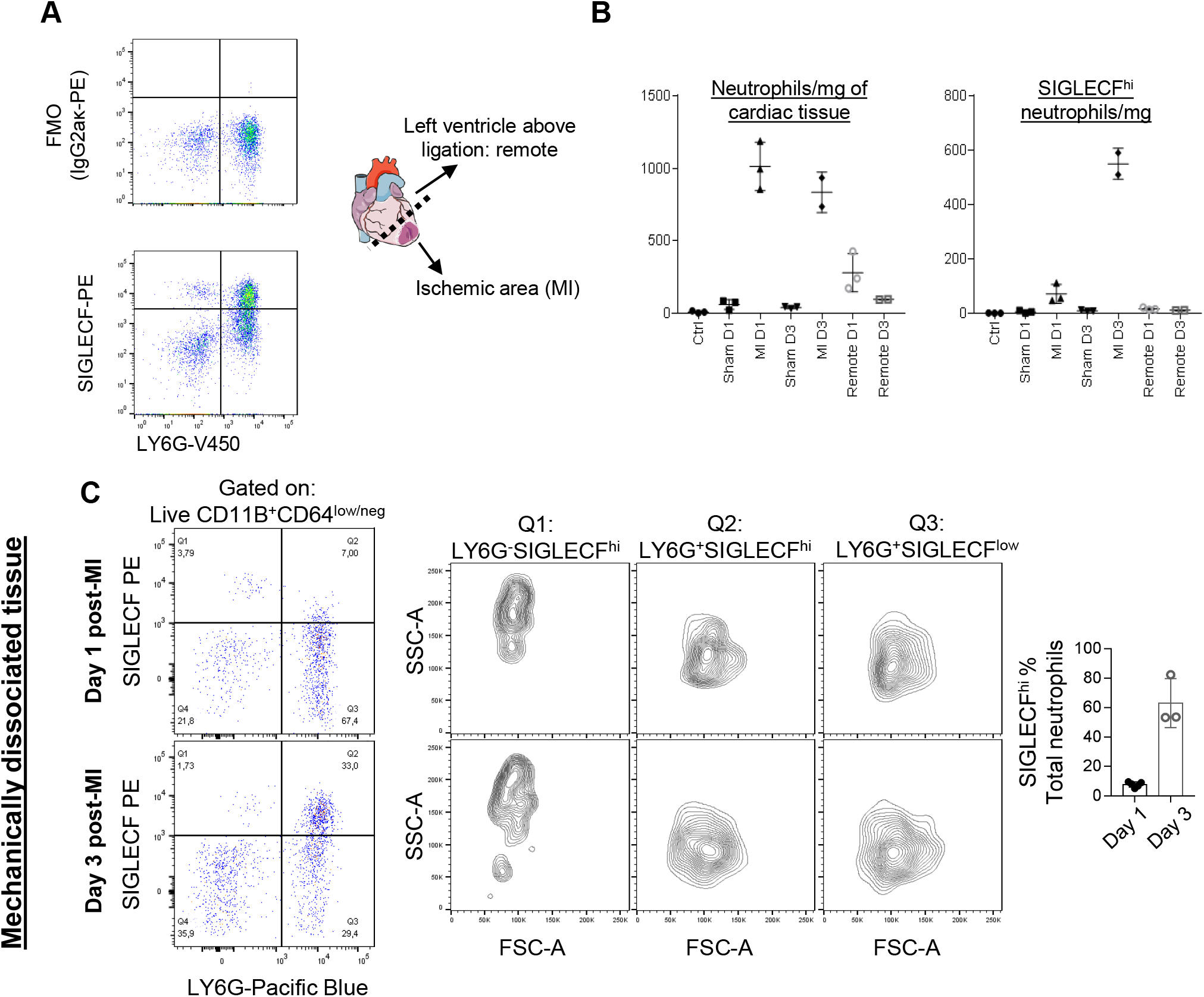
Additional flow cytometry analyses. **A)** Fluorescence minus one (FMO) validation of SIGLECF expression by cardiac neutrophils at Day 3 after MI; **B)** number of total neutrophils and SIGLECF^hi^ neutrophils in cardiac tissue from non-operated controls (Ctrl), sham and MI operated mice at Day 1 and 3 after myocardial infarction. To analyze remote, non ischemic myocardium, cardiac tissue was sampled above the ligation; **C)** detection of LY6G^neg^SIGLECF^hi^ eosinophils, LY6G^+^SIGLECF^hi^ neutrophils and LY6G^+^SIGLECF^low^ neutrophils in mechanically digested cardiac tissue at 1 and 3 days post MI (complementary to enzymatic digestion data in Figure 2).

**Figure S5:**
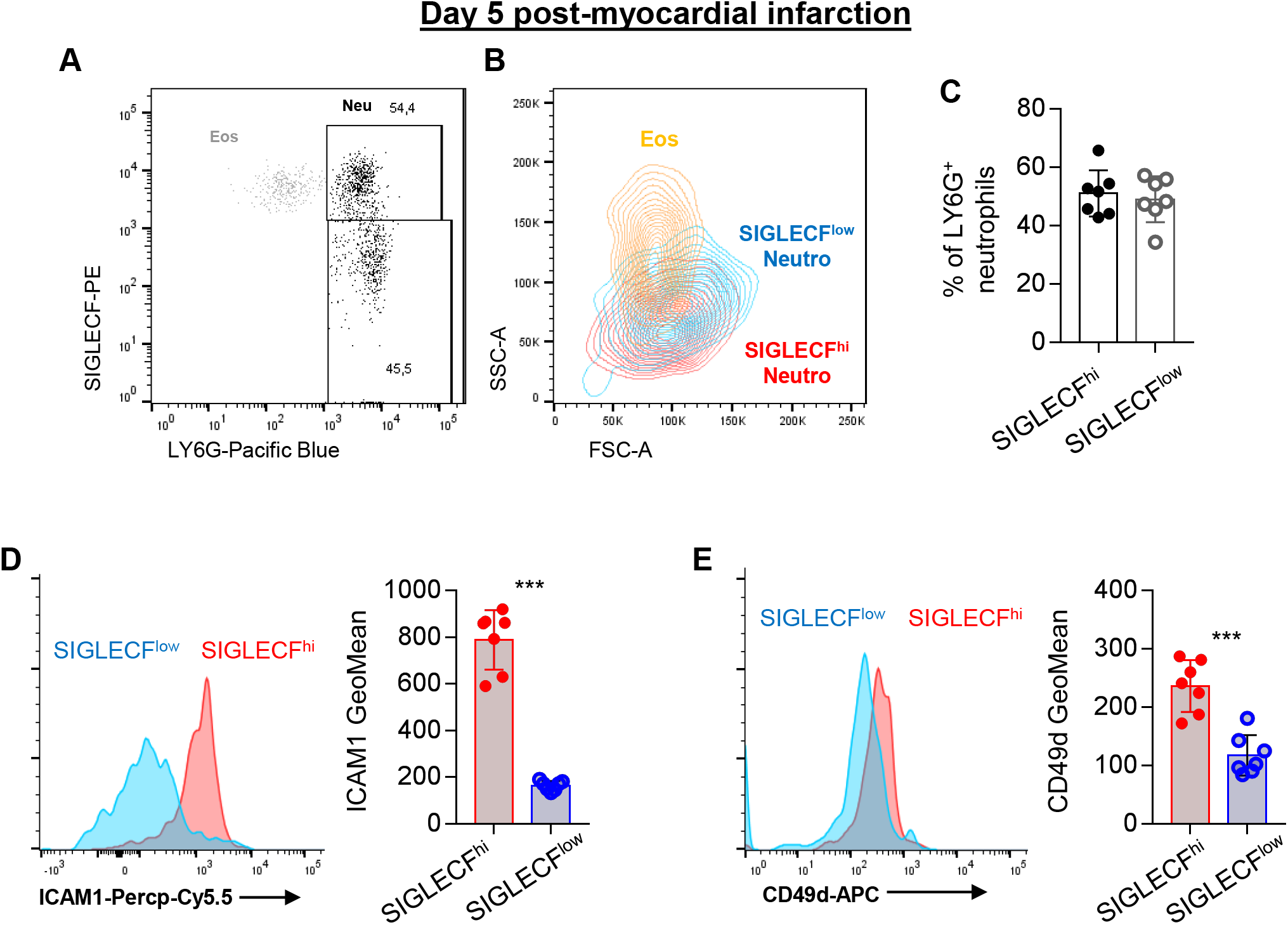
Flow cytometry analysis of neutrophil susbets in the heart 5 days after myocardial infarction. **A)** Representative dot plot showing expression of SIGLECF on eosinophils (Eos, grey) and neutrophils (Neu, black); **B)** SSC/FSC profile of neutrophil subsets and eosinophils at 5 days after myocardial infarction; **C)** proportion of SIGLECF^hi^ and SIGLECF^low^ neutrophils among total LY6G+ neutrophils; surface level of **D)** ICAM1 and **E)** CD49d in neutrophils subsets; ***p<0.001.

**Figure S6:**
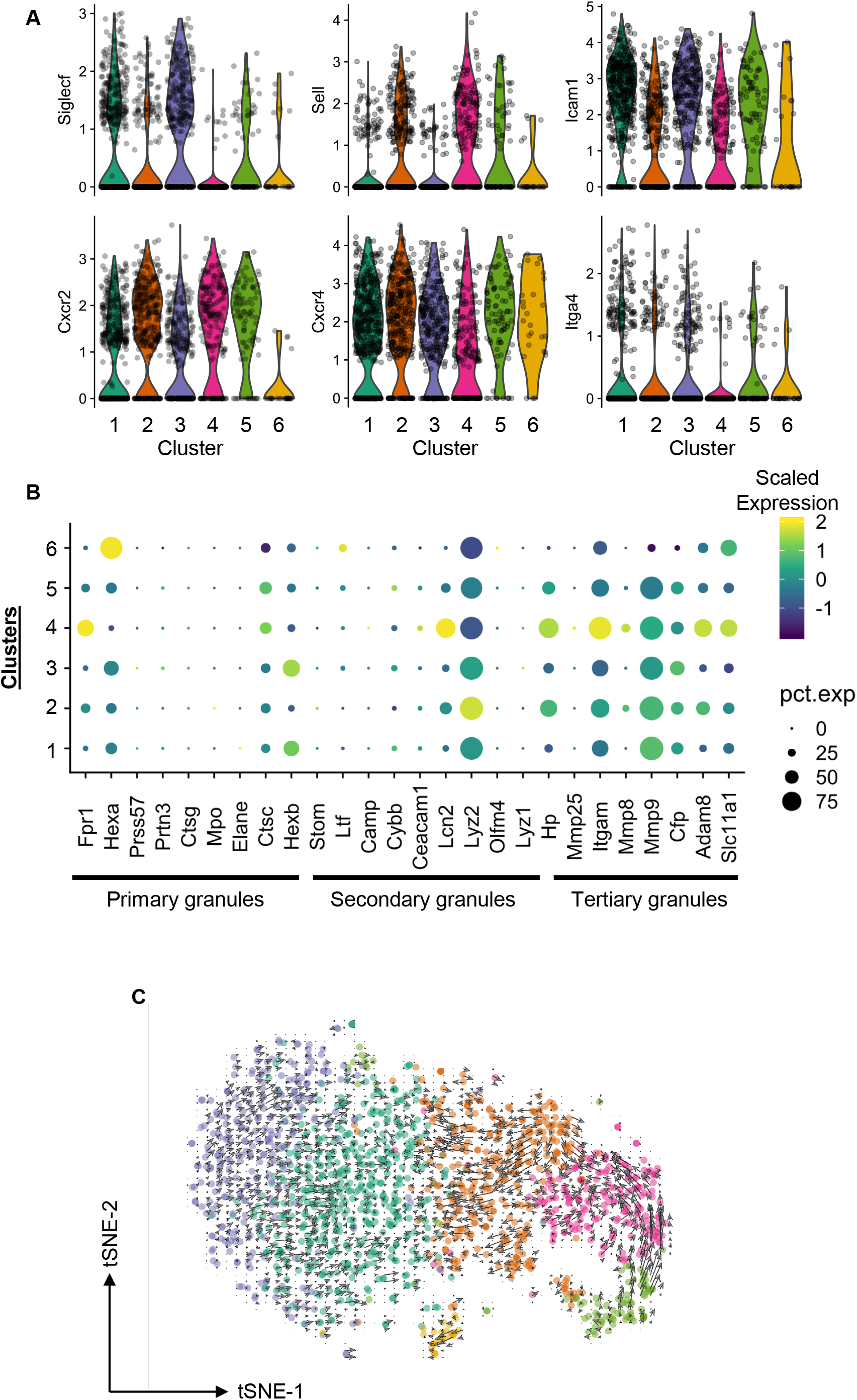
Expression of ageing/activation markers, neutrophil granule protein encoding genes and RNA-velocity analysis. **A)** Violin plots of log transformed expression of the indicated transcripts in the 6 neutrophil clusters (Figure 1); **B)** dotplot showing scaled expression (color scale) and proportion of cells (in percent, dot radius) expressing the indicated transcript in the 6 clusters; list of granule protein encoding genes adapted from Evrard et al. Immunity 2018; **C)** RNA-velocity analysis projected on the Seurat tSNE plot, with clusters color coded as in Figure 1.

**Figure S7:**
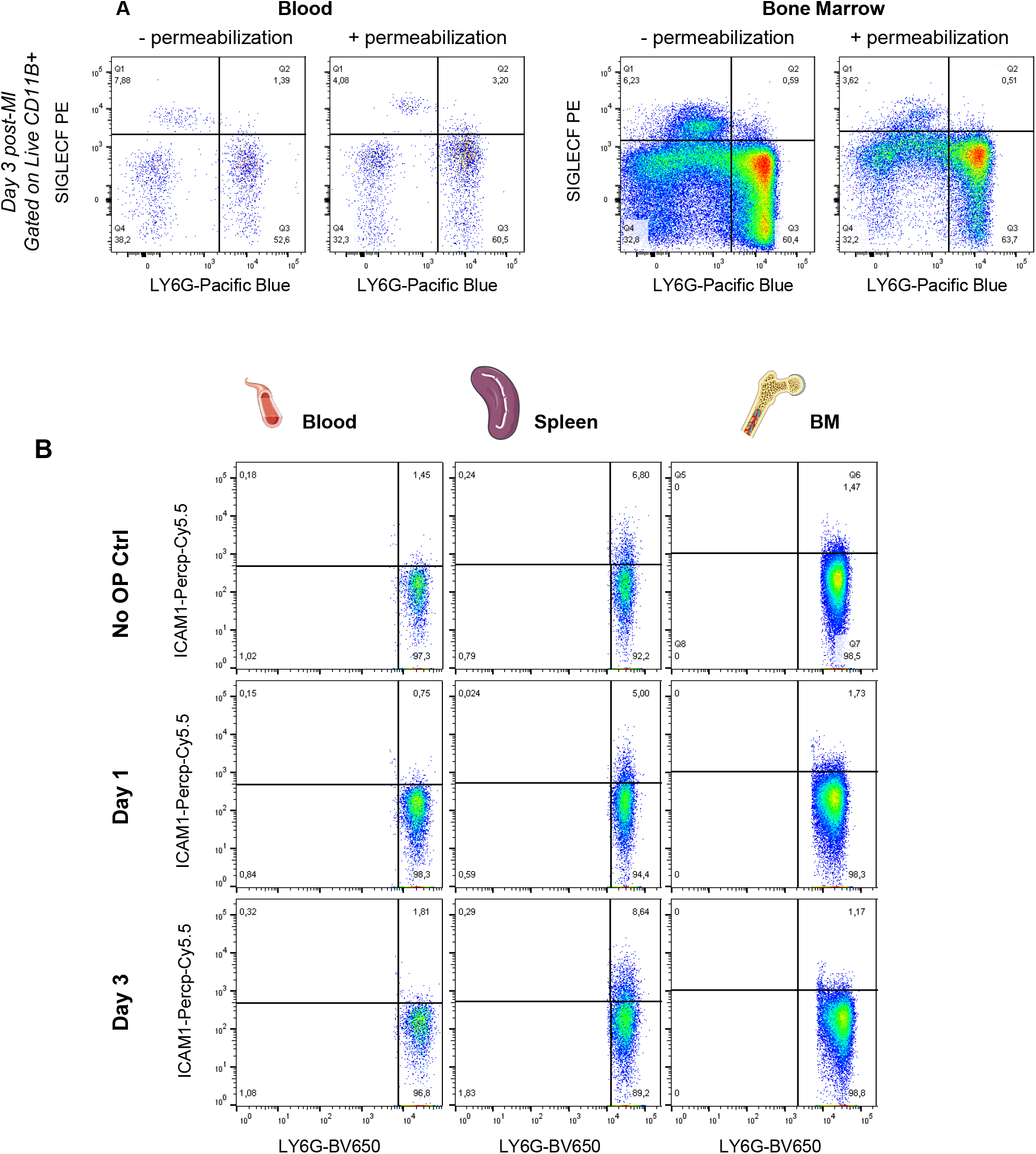
SIGLECF expression in blood and bone marrow neutrophils after permeabilization and ICAM1 expression in peripheral neutrophils. **A)** Representative flow cytometry plots of blood and bone marrow cells without (left) and with (right) permeabilization, cells were sampled in mice at 3 days post MI; **B)** ICAM1 expression in the indicated organs and treatment groups.

**Figure S8:**
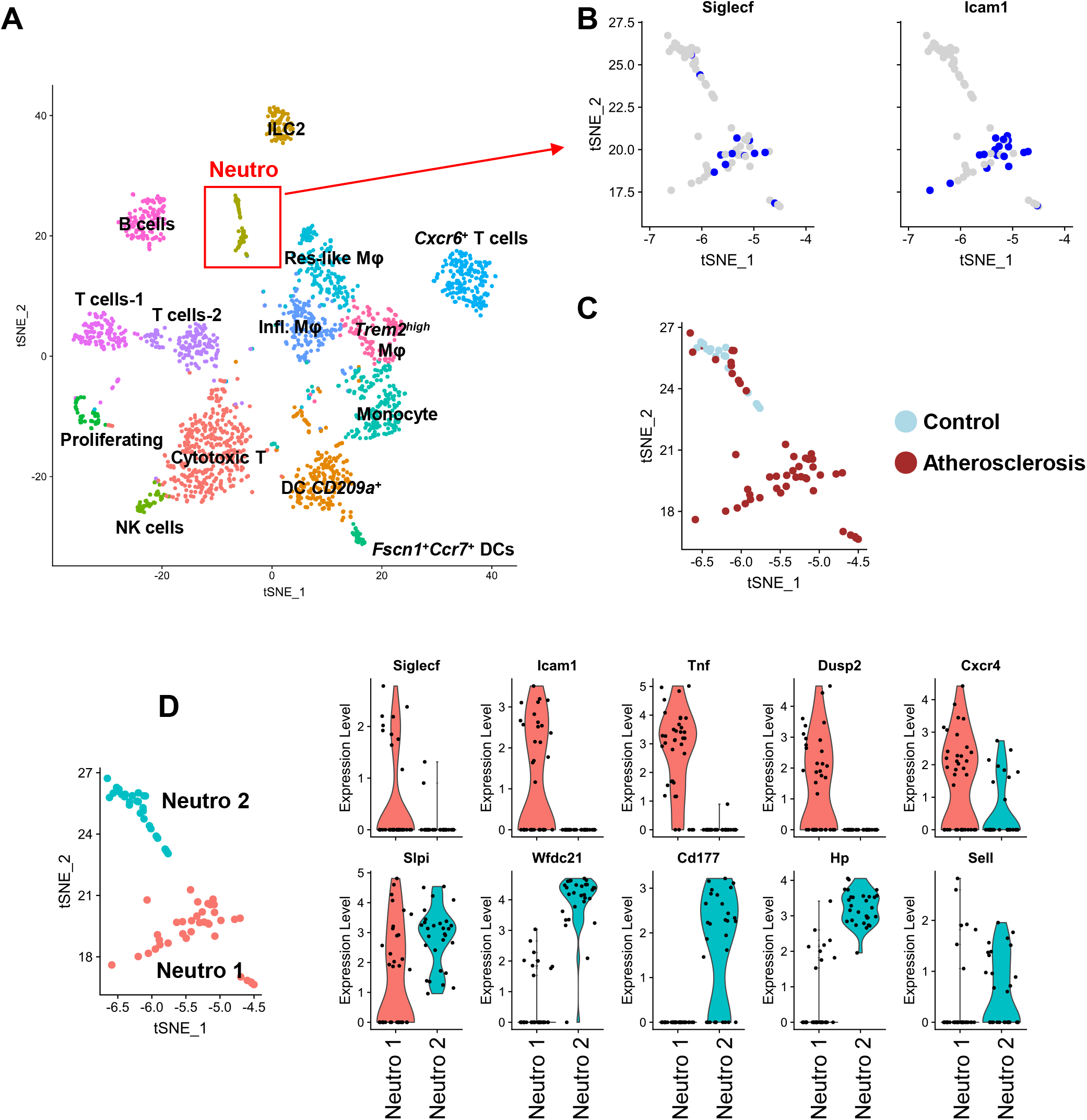
scRNA-seq of control and atherosclerotic aortas reveals two distinct neutrophil subsets. **A)** tSNE representation of single-cell RNA-seq gene expression data and clustering analysis of 2106 CD45^+^ cells isolated from control and atherosclerotic aortas from *Ldlr^−/−^* mice with identification of the major cell lineages; **B)** expression of *Siglecf* and *Icam1* in cells corresponding to neutrophils projected onto the tSNE plot (for clarity, an expression cutoff has been applied; blue= transcript detected; grey=transcript not detected); **C)** experimental condition of origin of neutrophils projected onto the tSNE plot; **D)** violin plot showing log normalized expression of the indicated genes in the two neutrophil populations.

