## Supplementary Table for "Time-resolved single-cell transcriptomics uncovers dynamics of cardiac neutrophil diversity in murine myocardial infarction"

### Marker genes for all clusters

| Column1 | Column2 | Column3 | Column4 | Column5 | Column6 | Column7 | Column8 |
|---|---|---|---|---|---|---|---|
| | p_val | avg_logFC | pct.1 | pct.2 | p_val_adj | cluster | gene |
| AA467197 | 1.07746511419719e-07 | 0.426232362789109 | 0.849 | 0.735 | 0.00334585241911652 | Cluster 1 | AA467197 |
| Nfkbia | 1.34538496024005e-07 | 0.402824073262992 | 0.994 | 0.977 | 0.00417782391703342 | Cluster 1 | Nfkbia |
| Tnf | 1.60435457677997e-07 | 0.785861835436737 | 0.959 | 0.801 | 0.00498200226727485 | Cluster 1 | Tnf |
| Cxcl2 | 8.28805519316247e-07 | 0.31496131206177 | 0.998 | 0.996 | 0.0257368977913274 | Cluster 1 | Cxcl2 |
| Rps19 | 1.5558607843176e-06 | 0.406626200160152 | 0.925 | 0.754 | 0.0483141449354144 | Cluster 1 | Rps19 |
| Ptgs2 | 2.8818522227558e-06 | 0.548152780488264 | 0.905 | 0.816 | 0.0894901570732359 | Cluster 1 | Ptgs2 |
| Rplp1 | 2.9409353648233e-06 | 0.375652622949679 | 0.973 | 0.902 | 0.091324865883858 | Cluster 1 | Rplp1 |
| Il23a | 3.07784500156183e-06 | 0.967582833266112 | 0.415 | 0.195 | 0.0955763208334994 | Cluster 1 | Il23a |
| Cst3 | 3.79649477390906e-06 | 0.256358696294138 | 0.977 | 0.931 | 0.117892552214198 | Cluster 1 | Cst3 |
| Eef1a1 | 5.40011358057686e-06 | 0.333527147577683 | 0.985 | 0.934 | 0.167689727017653 | Cluster 1 | Eef1a1 |
| Scimp | 5.83810903443694e-06 | 0.391134480311351 | 0.359 | 0.224 | 0.18129079984637 | Cluster 1 | Scimp |
| Dusp2 | 1.057861431416e-05 | 0.717223565853666 | 0.815 | 0.571 | 0.32849771029761 | Cluster 1 | Dusp2 |
| Mrpl52 | 1.22176363975199e-05 | 0.461194672283339 | 0.737 | 0.492 | 0.379394263052186 | Cluster 1 | Mrpl52 |
| Ppp1r15a | 1.34828684756645e-05 | 0.389921090692789 | 0.886 | 0.789 | 0.418683514774808 | Cluster 1 | Ppp1r15a |
| Rpsa | 1.38824077406515e-05 | 0.388787857688613 | 0.884 | 0.685 | 0.43109040757045 | Cluster 1 | Rpsa |
| Gpr84 | 1.5629754027553e-05 | 0.657425158277225 | 0.612 | 0.417 | 0.485350751817603 | Cluster 1 | Gpr84 |
| Rps8 | 1.87647885495739e-05 | 0.316876362134717 | 0.946 | 0.878 | 0.582702978829918 | Cluster 1 | Rps8 |
| Creb5 | 2.86006413571548e-05 | 0.45698435802995 | 0.425 | 0.267 | 0.888135716063728 | Cluster 1 | Creb5 |
| Ppia | 2.92457562520452e-05 | 0.398814111400489 | 0.919 | 0.723 | 0.908168468894761 | Cluster 1 | Ppia |
| Rps15 | 3.39130839475101e-05 | 0.347738992825893 | 0.896 | 0.733 | 1 | Cluster 1 | Rps15 |
| Icam1 | 3.8794400997916e-05 | 0.590715919089652 | 0.907 | 0.661 | 1 | Cluster 1 | Icam1 |
| Rpl8 | 5.19055157613071e-05 | 0.314575819737636 | 0.94 | 0.846 | 1 | Cluster 1 | Rpl8 |
| Snrpd3 | 5.65917744314633e-05 | 0.278831504741348 | 0.409 | 0.248 | 1 | Cluster 1 | Snrpd3 |
| Ptma | 7.9120897856531e-05 | 0.364811779960666 | 0.876 | 0.713 | 1 | Cluster 1 | Ptma |
| Clec4n | 0.000102071546330709 | 0.253036308860431 | 0.863 | 0.738 | 1 | Cluster 1 | Clec4n |
| Rplp2 | 0.000136870959992482 | 0.30420402556124 | 0.954 | 0.866 | 1 | Cluster 1 | Rplp2 |
| Tnfaip3 | 0.000143952200409666 | 0.550291473463288 | 0.923 | 0.821 | 1 | Cluster 1 | Tnfaip3 |
| Hexb | 0.000144280547166055 | 0.439102754966127 | 0.591 | 0.368 | 1 | Cluster 1 | Hexb |
| Ndufs7 | 0.000156069664996335 | 0.27914151171951 | 0.461 | 0.31 | 1 | Cluster 1 | Ndufs7 |
| Dynll1 | 0.000157625435919219 | 0.272555737604956 | 0.873 | 0.712 | 1 | Cluster 1 | Dynll1 |
| Socs3 | 0.000253077260751 | 0.531308134756251 | 0.703 | 0.555 | 1 | Cluster 1 | Socs3 |
| Rpl38 | 0.000340095329973267 | 0.36672249004735 | 0.936 | 0.801 | 1 | Cluster 1 | Rpl38 |
| Rps2 | 0.000344016993207447 | 0.317651470536516 | 0.905 | 0.783 | 1 | Cluster 1 | Rps2 |
| Rps28 | 0.000370522145390048 | 0.360630879376308 | 0.917 | 0.781 | 1 | Cluster 1 | Rps28 |
| Rps17 | 0.000423612077641998 | 0.250630743163015 | 0.701 | 0.541 | 1 | Cluster 1 | Rps17 |
| Snrpg | 0.000486913300382964 | 0.330872021593592 | 0.527 | 0.33 | 1 | Cluster 1 | Snrpg |
| Rplp0 | 0.000526966541107233 | 0.322224521584119 | 0.915 | 0.83 | 1 | Cluster 1 | Rplp0 |
| Rpl36 | 0.000548897638881645 | 0.356567058611313 | 0.886 | 0.703 | 1 | Cluster 1 | Rpl36 |
| Park7 | 0.000550688652153244 | 0.272638163630356 | 0.502 | 0.339 | 1 | Cluster 1 | Park7 |
| Kdm6b | 0.000589461261078344 | 0.277689363219109 | 0.973 | 0.923 | 1 | Cluster 1 | Kdm6b |
| Pfdn2 | 0.000651528030480848 | 0.27234019848055 | 0.531 | 0.386 | 1 | Cluster 1 | Pfdn2 |
| Gngt2 | 0.000661825422548764 | 0.351704492012859 | 0.786 | 0.592 | 1 | Cluster 1 | Gngt2 |
| Gm2a | 0.000738424157960473 | 0.275661972465368 | 0.757 | 0.654 | 1 | Cluster 1 | Gm2a |
| Ccrl2 | 0.000765270291263289 | 0.309942561430649 | 0.954 | 0.87 | 1 | Cluster 1 | Ccrl2 |
| G3bp1 | 0.000802517735530024 | 0.255666555083279 | 0.465 | 0.358 | 1 | Cluster 1 | G3bp1 |
| Rpl35a | 0.00102044581052598 | 0.265893616504243 | 0.965 | 0.925 | 1 | Cluster 1 | Rpl35a |
| B4galt1 | 0.00111556188709652 | 0.39873490123617 | 0.417 | 0.263 | 1 | Cluster 1 | B4galt1 |
| Nop10 | 0.00122465730920354 | 0.345751390808912 | 0.465 | 0.284 | 1 | Cluster 1 | Nop10 |
| Emc6 | 0.00141764146841542 | 0.284153324165772 | 0.396 | 0.263 | 1 | Cluster 1 | Emc6 |
| Rps26 | 0.00154205348200595 | 0.264758512751333 | 0.925 | 0.824 | 1 | Cluster 1 | Rps26 |
| N4bp1 | 0.0015486175049064 | 0.35240547541601 | 0.83 | 0.723 | 1 | Cluster 1 | N4bp1 |
| Nlrp3 | 0.0017587562092015 | 0.306897166092845 | 0.902 | 0.872 | 1 | Cluster 1 | Nlrp3 |
| Rps11 | 0.00183271527461443 | 0.290667635049645 | 0.917 | 0.795 | 1 | Cluster 1 | Rps11 |
| Uqcc2 | 0.00220187824563291 | 0.303163163457848 | 0.332 | 0.191 | 1 | Cluster 1 | Uqcc2 |
| Arf2 | 0.00232670255413717 | 0.377740050905511 | 0.473 | 0.327 | 1 | Cluster 1 | Arf2 |

| | | | | | | | |
|---|---|---|---|---|---|---|---|
| Atp5g2 | 0.00261014094229869 | 0.262415545271093 | 0.444 | 0.33 | 1 | Cluster 1 | Atp5g2 |
| Nme1 | 0.00266666877991743 | 0.394930242599663 | 0.498 | 0.319 | 1 | Cluster 1 | Nme1 |
| Clec5a | 0.00320345452279302 | 0.26004957396536 | 0.66 | 0.59 | 1 | Cluster 1 | Clec5a |
| Rps21 | 0.00349412451804139 | 0.295866276255663 | 0.944 | 0.845 | 1 | Cluster 1 | Rps21 |
| Rps18 | 0.00351143952163509 | 0.278930070380599 | 0.803 | 0.597 | 1 | Cluster 1 | Rps18 |
| Rpl12 | 0.00353427619001374 | 0.404864795050484 | 0.745 | 0.569 | 1 | Cluster 1 | Rpl12 |
| Atf4 | 0.0035783567216438 | 0.305299206770421 | 0.732 | 0.6 | 1 | Cluster 1 | Atf4 |
| Ptgs1 | 0.00451669937372853 | 0.345033551303347 | 0.444 | 0.28 | 1 | Cluster 1 | Ptgs1 |
| Snrpb | 0.00472825142858558 | 0.267519152154571 | 0.602 | 0.479 | 1 | Cluster 1 | Snrpb |
| Ahnak | 0.00475244205427365 | 0.26604336267272 | 0.263 | 0.207 | 1 | Cluster 1 | Ahnak |
| Ifrd1 | 0.00485315622333524 | 0.349449207767136 | 0.929 | 0.847 | 1 | Cluster 1 | Ifrd1 |
| Rpl36al | 0.00508638161274313 | 0.252869997555697 | 0.544 | 0.401 | 1 | Cluster 1 | Rpl36al |
| Pcbp1 | 0.00530925588453954 | 0.280250830068844 | 0.668 | 0.56 | 1 | Cluster 1 | Pcbp1 |
| Bcl2a1b | 0.00550680263796549 | 0.311714269527191 | 0.961 | 0.908 | 1 | Cluster 1 | Bcl2a1b |
| Cd81 | 0.00678805326495278 | 0.261633868439101 | 0.367 | 0.241 | 1 | Cluster 1 | Cd81 |
| Clec4e | 0.00687287414970645 | 0.268428363957768 | 0.967 | 0.948 | 1 | Cluster 1 | Clec4e |
| Tnfaip2 | 0.00718658632896282 | 0.309282491429228 | 0.915 | 0.852 | 1 | Cluster 1 | Tnfaip2 |
| Slpi1 | 1.09996386488299e-16 | 1.13678277148422 | 0.845 | 0.514 | 3.41571778962114e-12 | Cluster 2 | Slpi |
| S100a111 | 7.01547119368234e-12 | 0.665485922378742 | 0.978 | 0.957 | 2.17851426977418e-07 | Cluster 2 | S100a11 |
| Ifitm1 | 8.36158030037839e-12 | 0.909180419568715 | 0.923 | 0.752 | 2.5965215306765e-07 | Cluster 2 | Ifitm1 |
| Sorl1 | 4.23844537038078e-11 | 0.619005633691049 | 0.842 | 0.659 | 1.31616444086434e-06 | Cluster 2 | Sorl1 |
| Tyrobp | 4.38763515000378e-11 | 0.355019151702212 | 1 | 0.996 | 1.36249234313067e-06 | Cluster 2 | Tyrobp |
| Msrb11 | 4.62968261966709e-11 | 0.59593206966782 | 0.954 | 0.889 | 1.43765534388522e-06 | Cluster 2 | Msrb1 |
| Actg11 | 1.38095359115551e-10 | 0.529648947940327 | 0.994 | 0.984 | 4.28827518661522e-06 | Cluster 2 | Actg1 |
| Csf3r | 2.01895963859408e-10 | 0.460306487903435 | 0.978 | 0.915 | 6.2694753657262e-06 | Cluster 2 | Csf3r |
| S100a9 | 2.76699573387243e-10 | 0.6759438704346 | 0.963 | 0.925 | 8.59235185239405e-06 | Cluster 2 | S100a9 |
| Grina1 | 3.62341861107643e-10 | 0.604483332516927 | 0.913 | 0.743 | 1.12518018129756e-05 | Cluster 2 | Grina |
| Rnf149 | 4.46742296920846e-10 | 0.583422440840422 | 0.941 | 0.86 | 1.3872688546283e-05 | Cluster 2 | Rnf149 |
| Wfdc171 | 4.71709220026133e-10 | 0.804604764904323 | 0.783 | 0.507 | 1.46479864094715e-05 | Cluster 2 | Wfdc17 |
| Selplg1 | 9.58486240875036e-10 | 0.573539899406554 | 0.851 | 0.67 | 2.97638732378925e-05 | Cluster 2 | Selplg |
| Txn1 | 2.77504585457319e-09 | 0.419709469924447 | 0.929 | 0.851 | 8.61734989220614e-05 | Cluster 2 | Txn1 |
| Hp1 | 2.88570530433539e-09 | 0.576491286724817 | 0.659 | 0.408 | 8.96098068155268e-05 | Cluster 2 | Hp |
| Ifitm2 | 3.57305544340369e-09 | 0.58316455375773 | 0.929 | 0.924 | 0.000110954090684015 | Cluster 2 | Ifitm2 |
| S100a8 | 1.06531544687852e-08 | 0.694105971002742 | 0.978 | 0.909 | 0.000330812405719187 | Cluster 2 | S100a8 |
| Lsp1 | 1.5003130025245e-08 | 0.392497416768355 | 0.851 | 0.757 | 0.000465892196673933 | Cluster 2 | Lsp1 |
| Map1lc3b1 | 1.74900874326389e-08 | 0.361564755058824 | 0.882 | 0.83 | 0.000543119685045737 | Cluster 2 | Map1lc3b |
| Cxcr21 | 2.56428214598414e-08 | 0.381096145660024 | 0.799 | 0.606 | 0.000796286534792456 | Cluster 2 | Cxcr2 |
| Stfa2l1 | 2.60869276780052e-08 | 0.768567406743353 | 0.415 | 0.201 | 0.000810077365185095 | Cluster 2 | Stfa2l1 |
| S100a61 | 3.51827672832871e-08 | 0.586618701484635 | 0.932 | 0.78 | 0.00109253047244791 | Cluster 2 | S100a6 |
| Hdc | 7.70671127699777e-08 | 0.436260419829388 | 0.889 | 0.778 | 0.00239316505284612 | Cluster 2 | Hdc |
| Ctsd1 | 1.47636083863098e-07 | 0.463071617133214 | 0.981 | 0.921 | 0.00458454331220077 | Cluster 2 | Ctsd |
| Acod1 | 2.67037195297515e-07 | 0.595316359064902 | 0.944 | 0.792 | 0.00829230602557374 | Cluster 2 | Acod1 |
| Mxd11 | 5.75687739280443e-07 | 0.405048153242888 | 0.95 | 0.854 | 0.0178768313678756 | Cluster 2 | Mxd1 |
| Actb | 6.0894126046991e-07 | 0.357658704901226 | 1 | 1 | 0.0189094529613721 | Cluster 2 | Actb |
| Mcemp11 | 7.31400682706989e-07 | 0.586709458289661 | 0.659 | 0.394 | 0.0227121854001001 | Cluster 2 | Mcemp1 |
| Lmnb1 | 1.32331460000983e-06 | 0.399570300305577 | 0.95 | 0.897 | 0.0410928882741052 | Cluster 2 | Lmnb1 |
| Slc40a1 | 1.40294704499884e-06 | 0.494895763920314 | 0.254 | 0.093 | 0.0435657145883491 | Cluster 2 | Slc40a1 |
| Myl12b | 1.51558374516242e-06 | 0.427292178064563 | 0.907 | 0.79 | 0.0470634220385286 | Cluster 2 | Myl12b |
| Samsn1 | 1.61928108517062e-06 | 0.40592863565251 | 0.864 | 0.826 | 0.0502835355378032 | Cluster 2 | Samsn1 |
| Lrg11 | 2.16958151553135e-06 | 0.875645813336066 | 0.582 | 0.267 | 0.0673720148017951 | Cluster 2 | Lrg1 |
| Lyst | 2.57580745372552e-06 | 0.432365928432324 | 0.709 | 0.586 | 0.0799865488605386 | Cluster 2 | Lyst |
| Vps37b1 | 2.83359621664991e-06 | 0.857511604341725 | 0.681 | 0.413 | 0.0879916633156297 | Cluster 2 | Vps37b |
| Selenon1 | 3.50857109005097e-06 | 0.589646061251675 | 0.526 | 0.306 | 0.108951658059353 | Cluster 2 | Selenon |
| Asprv11 | 3.64326022515362e-06 | 1.12850345578452 | 0.458 | 0.162 | 0.113134159771695 | Cluster 2 | Asprv1 |
| Gda1 | 3.70295212731516e-06 | 0.59997719290973 | 0.492 | 0.274 | 0.114987772409518 | Cluster 2 | Gda |
| Tmsb4x | 4.07225261219704e-06 | 0.321032316961278 | 1 | 1 | 0.126455660366555 | Cluster 2 | Tmsb4x |
| Tpd52 | 4.27067075831854e-06 | 0.4010841982362 | 0.926 | 0.89 | 0.132617139058066 | Cluster 2 | Tpd52 |
| H3f3a | 4.79145030617242e-06 | 0.29184989764607 | 0.991 | 0.989 | 0.148788906357572 | Cluster 2 | H3f3a |
| Retnlg1 | 4.87101607862221e-06 | 0.310195123009392 | 0.44 | 0.166 | 0.151259662289456 | Cluster 2 | Retnlg |
| Slc16a31 | 5.2956947584987e-06 | 0.484275915740122 | 0.833 | 0.677 | 0.16444720933566 | Cluster 2 | Slc16a3 |
| Notch2 | 5.31278940748332e-06 | 0.390401071603828 | 0.406 | 0.265 | 0.16497804947058 | Cluster 2 | Notch2 |

| | | | | | | | |
|---|---|---|---|---|---|---|---|
| H2-D11 | 7.78932697354284e-06 | 0.310966227214542 | 0.972 | 0.957 | 0.241881970509426 | Cluster 2 | H2-D1 |
| Clec4d1 | 8.31474322574908e-06 | 0.487185239395 | 0.972 | 0.93 | 0.258197721389186 | Cluster 2 | Clec4d |
| Taldo11 | 9.22684542985164e-06 | 0.465713095679348 | 0.768 | 0.668 | 0.286521231133183 | Cluster 2 | Taldo1 |
| Rnf111 | 9.85044181253431e-06 | 0.396521206238754 | 0.489 | 0.335 | 0.305885769604628 | Cluster 2 | Rnf11 |
| Hmgb2 | 1.05328470805001e-05 | 0.294645082039449 | 0.799 | 0.728 | 0.32707650039077 | Cluster 2 | Hmgb2 |
| AC110211.11 | 1.39092978761454e-05 | 0.454170301656592 | 0.464 | 0.232 | 0.431925426947942 | Cluster 2 | AC110211.1 |
| Sirpb1c | 1.55843399763118e-05 | 0.292837199228801 | 0.622 | 0.51 | 0.483940509284409 | Cluster 2 | Sirpb1c |
| Ccl61 | 1.6598691263151e-05 | 0.364026671941972 | 0.693 | 0.498 | 0.515439159794629 | Cluster 2 | Ccl6 |
| Smox1 | 1.99503183711323e-05 | 0.343309685789588 | 0.659 | 0.491 | 0.61951723637877 | Cluster 2 | Smox |
| mt-Co11 | 2.2569756426942e-05 | 0.466848002840225 | 0.944 | 0.89 | 0.700858646325829 | Cluster 2 | mt-Co1 |
| Pnrc1 | 2.59845252183991e-05 | 0.297061504138946 | 0.957 | 0.884 | 0.806897461606946 | Cluster 2 | Pnrc1 |
| Gm5150 | 2.80179566408938e-05 | 0.462308242811918 | 0.486 | 0.284 | 0.870041607569677 | Cluster 2 | Gm5150 |
| Pla2g7 | 2.91537627922237e-05 | 0.566415709502097 | 0.663 | 0.469 | 0.905311795986924 | Cluster 2 | Pla2g7 |
| Itgal | 3.00348430003351e-05 | 0.398638143068563 | 0.44 | 0.302 | 0.932671979689406 | Cluster 2 | Itgal |
| Tgfbi | 3.03151335934897e-05 | 0.408824989207561 | 0.749 | 0.612 | 0.941375843478635 | Cluster 2 | Tgfbi |
| Rassf3 | 3.21548596305364e-05 | 0.502064211611011 | 0.362 | 0.182 | 0.998504856107048 | Cluster 2 | Rassf3 |
| Adam81 | 3.22831301581414e-05 | 0.473300362124591 | 0.514 | 0.337 | 1 | Cluster 2 | Adam8 |
| Sfxn5 | 3.35771215650453e-05 | 0.284745082289773 | 0.288 | 0.177 | 1 | Cluster 2 | Sfxn5 |
| Gpr35 | 3.76194855267439e-05 | 0.32001009840144 | 0.567 | 0.403 | 1 | Cluster 2 | Gpr35 |
| Sirpb1b | 4.15457321997205e-05 | 0.386514530563846 | 0.653 | 0.515 | 1 | Cluster 2 | Sirpb1b |
| Fam32a | 4.946736789906e-05 | 0.415254457355644 | 0.622 | 0.48 | 1 | Cluster 2 | Fam32a |
| Pirb | 5.00483345890905e-05 | 0.304774047773445 | 0.619 | 0.506 | 1 | Cluster 2 | Pirb |
| Entpd11 | 5.01597006230305e-05 | 0.573032936792071 | 0.684 | 0.506 | 1 | Cluster 2 | Entpd1 |
| Tnrc18 | 5.32901252894661e-05 | 0.284259821758793 | 0.461 | 0.36 | 1 | Cluster 2 | Tnrc18 |
| Por | 5.93334212706481e-05 | 0.461996114830487 | 0.458 | 0.307 | 1 | Cluster 2 | Por |
| Kpna4 | 6.95585724343315e-05 | 0.415456026018129 | 0.669 | 0.578 | 1 | Cluster 2 | Kpna4 |
| Il15 | 7.76082112061727e-05 | 0.337390024476643 | 0.186 | 0.066 | 1 | Cluster 2 | Il15 |
| Slc15a31 | 9.62974515708793e-05 | 0.269816089210458 | 0.715 | 0.625 | 1 | Cluster 2 | Slc15a3 |
| Sell1 | 9.96701924794698e-05 | 0.552346230106091 | 0.511 | 0.25 | 1 | Cluster 2 | Sell |
| March7 | 0.000103489464014914 | 0.290643462345735 | 0.749 | 0.683 | 1 | Cluster 2 | March7 |
| Pira2 | 0.000106580255978686 | 0.2774165752241 | 0.409 | 0.297 | 1 | Cluster 2 | Pira2 |
| Cd300ld | 0.000107688769906223 | 0.402567488493852 | 0.786 | 0.648 | 1 | Cluster 2 | Cd300ld |
| Rnf144a | 0.000117650554120492 | 0.285163842041921 | 0.149 | 0.032 | 1 | Cluster 2 | Rnf144a |
| AB124611 | 0.000135639282559449 | 0.506883215365431 | 0.492 | 0.287 | 1 | Cluster 2 | AB124611 |
| Cd52 | 0.000157432699243793 | 0.305028707045352 | 0.994 | 0.975 | 1 | Cluster 2 | Cd52 |
| Fyb | 0.000159684651704733 | 0.45555386224628 | 0.557 | 0.367 | 1 | Cluster 2 | Fyb |
| Efhd2 | 0.000160015459602468 | 0.266808551561118 | 0.861 | 0.798 | 1 | Cluster 2 | Efhd2 |
| Abr | 0.000184164463018243 | 0.334749787713288 | 0.511 | 0.392 | 1 | Cluster 2 | Abr |
| H2-Q10 | 0.000197179494149499 | 0.391048370240272 | 0.526 | 0.433 | 1 | Cluster 2 | H2-Q10 |
| Slc7a11 | 0.000219612231772208 | 0.342771676781173 | 0.932 | 0.833 | 1 | Cluster 2 | Slc7a11 |
| Fmnl11 | 0.000227345264251468 | 0.343305947010694 | 0.576 | 0.471 | 1 | Cluster 2 | Fmnl1 |
| Slc22a15 | 0.000243662164373007 | 0.455171875271327 | 0.307 | 0.181 | 1 | Cluster 2 | Slc22a15 |
| Fam129a | 0.000246156276283765 | 0.353969401475147 | 0.31 | 0.2 | 1 | Cluster 2 | Fam129a |
| Btg1 | 0.000246451650457695 | 0.284627699401775 | 0.988 | 0.979 | 1 | Cluster 2 | Btg1 |
| Hif1a1 | 0.00025248161465344 | 0.378947002795304 | 0.477 | 0.37 | 1 | Cluster 2 | Hif1a |
| Sema4a | 0.000253473870454476 | 0.515845665408783 | 0.567 | 0.343 | 1 | Cluster 2 | Sema4a |
| Ankrd33b | 0.000258946267887724 | 0.393297325116969 | 0.7 | 0.577 | 1 | Cluster 2 | Ankrd33b |
| Sh2d3c1 | 0.000301441706813341 | 0.424394189470592 | 0.257 | 0.134 | 1 | Cluster 2 | Sh2d3c |
| Ube2b | 0.000316000701184202 | 0.267553339976429 | 0.774 | 0.697 | 1 | Cluster 2 | Ube2b |
| Ncf2 | 0.000395350832486464 | 0.442376756700418 | 0.786 | 0.637 | 1 | Cluster 2 | Ncf2 |
| Alox5ap | 0.000424360821696297 | 0.270339682993762 | 0.954 | 0.929 | 1 | Cluster 2 | Alox5ap |
| Cnn21 | 0.000472536890294284 | 0.478874760380561 | 0.399 | 0.221 | 1 | Cluster 2 | Cnn2 |
| Sat11 | 0.00048152530084281 | 0.377018206328694 | 0.759 | 0.718 | 1 | Cluster 2 | Sat1 |
| Pglyrp11 | 0.00049024850697517 | 0.499749769971269 | 0.628 | 0.483 | 1 | Cluster 2 | Pglyrp1 |
| Coro1a | 0.000504562727337199 | 0.266502778069595 | 0.923 | 0.901 | 1 | Cluster 2 | Coro1a |
| Klra2 | 0.000591895253617288 | 0.279239827602268 | 0.266 | 0.172 | 1 | Cluster 2 | Klra2 |
| Gm26740 | 0.000725224747526403 | 0.339539112174642 | 0.573 | 0.45 | 1 | Cluster 2 | Gm26740 |
| Nudt4 | 0.000734130892300938 | 0.269308640966737 | 0.687 | 0.643 | 1 | Cluster 2 | Nudt4 |
| Svil | 0.000742504941780134 | 0.32659304923823 | 0.359 | 0.235 | 1 | Cluster 2 | Svil |
| Cpne2 | 0.000742755452261896 | 0.498028096986798 | 0.341 | 0.191 | 1 | Cluster 2 | Cpne2 |
| mt-Co3 | 0.000814547766365691 | 0.415569455258724 | 0.969 | 0.952 | 1 | Cluster 2 | mt-Co3 |

| | | | | | | | |
|---|---|---|---|---|---|---|---|
| Osgin1 | 0.000832460222074312 | 0.509723600607553 | 0.325 | 0.174 | 1 | Cluster 2 | Osgin1 |
| Ripor21 | 0.000873245701600644 | 0.347632895381468 | 0.307 | 0.163 | 1 | Cluster 2 | Ripor2 |
| Spag9 | 0.000919543074306106 | 0.270377396710089 | 0.864 | 0.841 | 1 | Cluster 2 | Spag9 |
| Dhrs7 | 0.000936741723271207 | 0.263141122445055 | 0.536 | 0.44 | 1 | Cluster 2 | Dhrs7 |
| Impact | 0.000942330758571201 | 0.33928090234921 | 0.198 | 0.1 | 1 | Cluster 2 | Impact |
| Gm34084 | 0.00101269110794444 | 0.442518582810234 | 0.474 | 0.341 | 1 | Cluster 2 | Gm34084 |
| Sgms21 | 0.00109064343891624 | 0.386236676364076 | 0.483 | 0.312 | 1 | Cluster 2 | Sgms2 |
| Litaf | 0.00118389650406162 | 0.28957629657025 | 0.969 | 0.948 | 1 | Cluster 2 | Litaf |
| Steap4 | 0.00123176805885791 | 0.453276432469548 | 0.195 | 0.077 | 1 | Cluster 2 | Steap4 |
| Slfn1 | 0.0013879490571325 | 0.279048822990151 | 0.672 | 0.522 | 1 | Cluster 2 | Slfn1 |
| Ndel1 | 0.00148233642897891 | 0.33167572933598 | 0.848 | 0.777 | 1 | Cluster 2 | Ndel1 |
| Xylt1 | 0.00159876906583634 | 0.287408011844054 | 0.396 | 0.306 | 1 | Cluster 2 | Xylt1 |
| Hcls1 | 0.00163194909993542 | 0.29233638625817 | 0.672 | 0.544 | 1 | Cluster 2 | Hcls1 |
| Emb | 0.00173572030481754 | 0.365012462467355 | 0.653 | 0.498 | 1 | Cluster 2 | Emb |
| Trps1 | 0.00196042292503163 | 0.332804270601071 | 0.344 | 0.23 | 1 | Cluster 2 | Trps1 |
| Tlr21 | 0.00202613288468162 | 0.495737809614462 | 0.542 | 0.376 | 1 | Cluster 2 | Tlr2 |
| Gm39321 | 0.00206144073911898 | 0.358200631627359 | 0.285 | 0.149 | 1 | Cluster 2 | Gm39321 |
| Rhov | 0.00244093236057068 | 0.409727219866136 | 0.266 | 0.147 | 1 | Cluster 2 | Rhov |
| Pag1 | 0.00249604211895716 | 0.286056020959238 | 0.344 | 0.26 | 1 | Cluster 2 | Pag1 |
| Trem3 | 0.00251738971726384 | 0.386317271294575 | 0.464 | 0.35 | 1 | Cluster 2 | Trem3 |
| Ftl1 | 0.00260400400272695 | 0.292748518010882 | 1 | 1 | 1 | Cluster 2 | Ftl1 |
| Zyx | 0.002729899532668 | 0.342004812577763 | 0.675 | 0.591 | 1 | Cluster 2 | Zyx |
| Pbx1 | 0.00304734829969412 | 0.300452597161885 | 0.337 | 0.208 | 1 | Cluster 2 | Pbx1 |
| Trpm2 | 0.0030886773080784 | 0.285467315743817 | 0.164 | 0.062 | 1 | Cluster 2 | Trpm2 |
| Adipor1 | 0.00328092187324165 | 0.289612001703426 | 0.762 | 0.694 | 1 | Cluster 2 | Adipor1 |
| 1600010M07Rik | 0.00342373764715376 | 0.37567515652778 | 0.464 | 0.307 | 1 | Cluster 2 | 1600010M07Rik |
| Rps9 | 0.00367049325655096 | 0.306134039950372 | 0.975 | 0.97 | 1 | Cluster 2 | Rps9 |
| Ttc7 | 0.00381696277144477 | 0.25037695363342 | 0.276 | 0.179 | 1 | Cluster 2 | Ttc7 |
| Nktr | 0.00419165318662164 | 0.343215327843218 | 0.61 | 0.57 | 1 | Cluster 2 | Nktr |
| Ccr1 | 0.00478744105716813 | 0.330410675331984 | 0.944 | 0.899 | 1 | Cluster 2 | Ccr1 |
| Tspan13 | 0.00489755023496894 | 0.306505374068448 | 0.353 | 0.246 | 1 | Cluster 2 | Tspan13 |
| Chil11 | 0.00495553247042272 | 0.395670934456937 | 0.307 | 0.195 | 1 | Cluster 2 | Chil1 |
| Suco | 0.00556423460483505 | 0.279604214462568 | 0.35 | 0.273 | 1 | Cluster 2 | Suco |
| Mrpl331 | 0.00583345675905631 | 0.417211982876265 | 0.653 | 0.568 | 1 | Cluster 2 | Mrpl33 |
| Samhd11 | 0.00614789116538828 | 0.255147527750626 | 0.697 | 0.595 | 1 | Cluster 2 | Samhd1 |
| Sp1401 | 0.0061757959607001 | 0.256906129197785 | 0.415 | 0.321 | 1 | Cluster 2 | Sp140 |
| Nedd91 | 0.00618051549880362 | 0.259636014686968 | 0.26 | 0.155 | 1 | Cluster 2 | Nedd9 |
| Lyz2 | 0.00667790336077357 | 0.258988359613323 | 0.935 | 0.911 | 1 | Cluster 2 | Lyz2 |
| Arpc3 | 0.0077293558026821 | 0.293424517288422 | 0.944 | 0.936 | 1 | Cluster 2 | Arpc3 |
| Dhrs9 | 0.00776438344099211 | 0.260787518459238 | 0.254 | 0.154 | 1 | Cluster 2 | Dhrs9 |
| Hnrnph2 | 0.0081359795879238 | 0.304209151455502 | 0.576 | 0.485 | 1 | Cluster 2 | Hnrnph2 |
| Jdp2 | 0.00862654856383312 | 0.317844735877289 | 0.749 | 0.653 | 1 | Cluster 2 | Jdp2 |
| Sephs2 | 0.00868523155829351 | 0.334524106281115 | 0.238 | 0.137 | 1 | Cluster 2 | Sephs2 |
| Tmcc1 | 0.00877901134692887 | 0.440398629856615 | 0.464 | 0.318 | 1 | Cluster 2 | Tmcc1 |
| Tnfrsf23 | 0.00998191396666739 | 0.306683761726534 | 0.502 | 0.439 | 1 | Cluster 2 | Tnfrsf23 |
| Cox7a21 | 6.5614068633359e-17 | 0.646530648697613 | 0.934 | 0.638 | 2.0375136732717e-12 | Cluster 3 | Cox7a2 |
| Snrpe1 | 1.45482660480381e-14 | 0.639641059660128 | 0.707 | 0.304 | 4.51767305589727e-10 | Cluster 3 | Snrpe |
| Ppia2 | 3.47619825406233e-14 | 0.583048252344196 | 0.976 | 0.742 | 1.07946384383398e-09 | Cluster 3 | Ppia |
| Rpl191 | 1.15313178838209e-13 | 0.495823188318264 | 0.976 | 0.838 | 3.58082014246289e-09 | Cluster 3 | Rpl19 |
| Rgs101 | 1.67087130857525e-13 | 0.59033737858887 | 0.557 | 0.199 | 5.18855667451874e-09 | Cluster 3 | Rgs10 |
| Rpl35a2 | 2.04128118277881e-13 | 0.388102011946833 | 0.997 | 0.924 | 6.33879045688305e-09 | Cluster 3 | Rpl35a |
| Rps27 | 4.56789734393067e-13 | 0.568719370718437 | 0.979 | 0.785 | 1.41846916221079e-08 | Cluster 3 | Rps2 |
| Rps241 | 4.90207407328294e-13 | 0.467497507360235 | 0.941 | 0.816 | 1.52224106197655e-08 | Cluster 3 | Rps24 |
| Rpl321 | 6.06248116932769e-13 | 0.553974768788973 | 0.958 | 0.772 | 1.88258227751133e-08 | Cluster 3 | Rpl32 |
| Rpl151 | 9.32100523161901e-13 | 0.53298511276364 | 0.875 | 0.574 | 2.89445175457465e-08 | Cluster 3 | Rpl15 |
| Rps182 | 1.09847846664965e-12 | 0.612191284575615 | 0.906 | 0.606 | 3.41110518248716e-08 | Cluster 3 | Rps18 |
| Rpl181 | 1.35872560094919e-12 | 0.52336444524199 | 0.979 | 0.819 | 4.21925060862751e-08 | Cluster 3 | Rpl18 |
| Rpl221 | 1.3731260515569e-12 | 0.588562526232624 | 0.944 | 0.635 | 4.26396832789963e-08 | Cluster 3 | Rpl22 |
| Rps262 | 1.40651791186214e-12 | 0.529886957432252 | 0.979 | 0.828 | 4.36766007170551e-08 | Cluster 3 | Rps26 |
| Rps231 | 2.44092210456078e-12 | 0.534672124953177 | 0.979 | 0.753 | 7.57979541129258e-08 | Cluster 3 | Rps23 |
| Rpl362 | 3.63609067617573e-12 | 0.583733621413026 | 0.969 | 0.714 | 1.12911523767285e-07 | Cluster 3 | Rpl36 |

| | | | | | | | |
|---|---|---|---|---|---|---|---|
| Rps192 | 4.95053483376858e-12 | 0.598409236972505 | 0.979 | 0.77 | 1.53728958193016e-07 | Cluster 3 | Rps19 |
| Ybx11 | 4.953061760519e-12 | 0.493458522436862 | 0.875 | 0.581 | 1.53807426849397e-07 | Cluster 3 | Ybx1 |
| Rpl261 | 7.32531919617706e-12 | 0.507743727232499 | 0.948 | 0.812 | 2.27473136998886e-07 | Cluster 3 | Rpl26 |
| Rpl281 | 1.66507088389248e-11 | 0.530046893957429 | 0.955 | 0.771 | 5.17054461575131e-07 | Cluster 3 | Rpl28 |
| Rpl382 | 2.87128488319884e-11 | 0.450915157492334 | 0.969 | 0.816 | 8.91620094779737e-07 | Cluster 3 | Rpl38 |
| Btf31 | 2.95009445585495e-11 | 0.42264047779018 | 0.93 | 0.683 | 9.16092831376637e-07 | Cluster 3 | Btf3 |
| Cox6a11 | 4.25745859926945e-11 | 0.454557901858432 | 0.857 | 0.598 | 1.32206861883114e-06 | Cluster 3 | Cox6a1 |
| Rpsa2 | 1.01630068349547e-10 | 0.574991581813253 | 0.944 | 0.704 | 3.15591851245848e-06 | Cluster 3 | Rpsa |
| Rplp02 | 1.26819026673488e-10 | 0.516942831393413 | 0.986 | 0.826 | 3.93811123529182e-06 | Cluster 3 | Rplp0 |
| Mrps141 | 1.37313609347193e-10 | 0.518097229439345 | 0.505 | 0.184 | 4.2639995110584e-06 | Cluster 3 | Mrps14 |
| Rplp12 | 1.50610737345698e-10 | 0.473729188519305 | 0.983 | 0.912 | 4.67691522679596e-06 | Cluster 3 | Rplp1 |
| Chchd21 | 2.1108843535529e-10 | 0.396637037332561 | 0.976 | 0.856 | 6.55492918308781e-06 | Cluster 3 | Chchd2 |
| Rplp22 | 2.13235908129938e-10 | 0.40240093834436 | 0.986 | 0.873 | 6.62161465515897e-06 | Cluster 3 | Rplp2 |
| Rbx11 | 3.15565328079251e-10 | 0.575662428227724 | 0.854 | 0.503 | 9.79925013284499e-06 | Cluster 3 | Rbx1 |
| Rps172 | 4.48183595606778e-10 | 0.484074314232631 | 0.829 | 0.536 | 1.39174451943773e-05 | Cluster 3 | Rps17 |
| Ebna1bp2 | 6.31989144994392e-10 | 0.380924152395424 | 0.39 | 0.137 | 1.96251589195109e-05 | Cluster 3 | Ebna1bp2 |
| Rps3 | 6.94653111465146e-10 | 0.452253591371756 | 0.976 | 0.8 | 2.15710630703272e-05 | Cluster 3 | Rps3 |
| Dad11 | 7.15950837542249e-10 | 0.450081355413606 | 0.638 | 0.344 | 2.22324213581995e-05 | Cluster 3 | Dad1 |
| Rpl34 | 1.26431112705631e-09 | 0.438937838573738 | 0.976 | 0.833 | 3.92606534284794e-05 | Cluster 3 | Rpl34 |
| Siglecf1 | 1.60011718512897e-09 | 0.60565269586993 | 0.672 | 0.328 | 4.96884389498098e-05 | Cluster 3 | Siglecf |
| Cnbp1 | 3.56383256247556e-09 | 0.540564317726775 | 0.826 | 0.493 | 0.000110667692562554 | Cluster 3 | Cnbp |
| Ndufa131 | 4.10510261734151e-09 | 0.46706838944814 | 0.822 | 0.508 | 0.000127475751576306 | Cluster 3 | Ndufa13 |
| Tomm7 | 4.62987052412963e-09 | 0.383926174265138 | 0.836 | 0.603 | 0.000143771369385798 | Cluster 3 | Tomm7 |
| Rpl18a | 5.11158367193978e-09 | 0.40558487133445 | 0.969 | 0.862 | 0.000158730007764746 | Cluster 3 | Rpl18a |
| Rpl82 | 5.52203367436465e-09 | 0.409364320866935 | 0.983 | 0.852 | 0.000171475711690046 | Cluster 3 | Rpl8 |
| Rps12 | 8.21813906475006e-09 | 0.400533262191256 | 0.993 | 0.911 | 0.000255197872377684 | Cluster 3 | Rps12 |
| Cox7b1 | 8.37139682927907e-09 | 0.522959170053083 | 0.721 | 0.376 | 0.000259956985739603 | Cluster 3 | Cox7b |
| Ptma2 | 1.02170249593666e-08 | 0.47195728386834 | 0.955 | 0.721 | 0.000317269276063212 | Cluster 3 | Ptma |
| Atp5o.11 | 1.07372837513253e-08 | 0.431715874602035 | 0.502 | 0.209 | 0.000333424872329905 | Cluster 3 | Atp5o.1 |
| Ndufa111 | 1.24168665867936e-08 | 0.526568276776146 | 0.352 | 0.074 | 0.0003855809581197 | Cluster 3 | Ndufa11 |
| Rps112 | 1.70014458620559e-08 | 0.46163489200465 | 0.958 | 0.806 | 0.000527945898354421 | Cluster 3 | Rps11 |
| Gngt21 | 1.88386914453746e-08 | 0.332431952884984 | 0.857 | 0.607 | 0.000584997885453216 | Cluster 3 | Gngt2 |
| Rps51 | 2.45217776520991e-08 | 0.515102963357546 | 0.944 | 0.669 | 0.000761474761430633 | Cluster 3 | Rps5 |
| Rps212 | 2.78314048276849e-08 | 0.491774877393155 | 0.976 | 0.854 | 0.0008642486141141 | Cluster 3 | Rps21 |
| Snu13 | 3.41026145851968e-08 | 0.460831319165179 | 0.634 | 0.304 | 0.00105898849071412 | Cluster 3 | Snu13 |
| Rps291 | 3.89544656687361e-08 | 0.424466124388999 | 0.997 | 0.961 | 0.00120965302241126 | Cluster 3 | Rps29 |
| Rpl27a | 4.94744880645567e-08 | 0.390406087097628 | 0.969 | 0.853 | 0.00153633127786868 | Cluster 3 | Rpl27a |
| mt-Nd2 | 5.39887641164889e-08 | 0.51408640805195 | 0.868 | 0.654 | 0.00167651309210933 | Cluster 3 | mt-Nd2 |
| Ngdn | 5.61577102169129e-08 | 0.25505053197029 | 0.317 | 0.139 | 0.0017438653753658 | Cluster 3 | Ngdn |
| Snrpg2 | 5.92761935162609e-08 | 0.524478436028114 | 0.683 | 0.325 | 0.00184070363726045 | Cluster 3 | Snrpg |
| U2af11 | 6.01190879478743e-08 | 0.43442500419955 | 0.551 | 0.249 | 0.00186687803804534 | Cluster 3 | U2af1 |
| Srsf31 | 6.08934604023067e-08 | 0.469510350161675 | 0.756 | 0.456 | 0.00189092462587283 | Cluster 3 | Srsf3 |
| Rps131 | 6.33091478609035e-08 | 0.443673946038502 | 0.979 | 0.845 | 0.00196593896852464 | Cluster 3 | Rps13 |
| Rps27a | 6.38456295278179e-08 | 0.34531560714117 | 0.983 | 0.889 | 0.00198259833372733 | Cluster 3 | Rps27a |
| Ndufb51 | 8.05940271161547e-08 | 0.460680219030755 | 0.484 | 0.198 | 0.00250268632403795 | Cluster 3 | Ndufb5 |
| Mtdh1 | 8.18039629548915e-08 | 0.430340028755151 | 0.739 | 0.408 | 0.00254025846163824 | Cluster 3 | Mtdh |
| Ost41 | 9.16497052021096e-08 | 0.369327311795109 | 0.906 | 0.673 | 0.00284599829564111 | Cluster 3 | Ost4 |
| Krtcap21 | 9.65003929642509e-08 | 0.385855578348383 | 0.394 | 0.151 | 0.00299662670271888 | Cluster 3 | Krtcap2 |
| Rps82 | 9.86195247540507e-08 | 0.36034344482368 | 0.993 | 0.877 | 0.00306243210218754 | Cluster 3 | Rps8 |
| Pfdn6 | 1.01923556963242e-07 | 0.440444524389888 | 0.568 | 0.266 | 0.00316503221437955 | Cluster 3 | Pfdn6 |
| Rps16 | 1.05985522034373e-07 | 0.347680313145585 | 0.993 | 0.929 | 0.00329116841573338 | Cluster 3 | Rps16 |
| Cacybp | 1.07745329333153e-07 | 0.300566525293653 | 0.307 | 0.119 | 0.00334581571178241 | Cluster 3 | Cacybp |
| Rps282 | 1.16823860202635e-07 | 0.420609359685398 | 0.965 | 0.792 | 0.00362773133087243 | Cluster 3 | Rps28 |
| Hexb2 | 1.18202251203019e-07 | 0.504615881203238 | 0.669 | 0.387 | 0.00367053450660735 | Cluster 3 | Hexb |
| Psma31 | 1.2541159128307e-07 | 0.351284859012395 | 0.666 | 0.381 | 0.00389440614411318 | Cluster 3 | Psma3 |
| Minos11 | 1.41614062078539e-07 | 0.412618512061651 | 0.774 | 0.474 | 0.00439754146972488 | Cluster 3 | Minos1 |
| Rps201 | 2.19317636161198e-07 | 0.419049324588517 | 0.902 | 0.675 | 0.00681047055571369 | Cluster 3 | Rps20 |
| Cox20 | 2.30920320408848e-07 | 0.317260212502627 | 0.366 | 0.128 | 0.00717076870965597 | Cluster 3 | Cox20 |
| Rpl30 | 2.36296931096082e-07 | 0.379042612561407 | 0.941 | 0.804 | 0.00733772860132663 | Cluster 3 | Rpl30 |
| Ndufs61 | 2.37104236078787e-07 | 0.485135318781294 | 0.551 | 0.242 | 0.00736279784295457 | Cluster 3 | Ndufs6 |
| Rps152 | 2.47565843446015e-07 | 0.395103510717422 | 0.927 | 0.754 | 0.0076876621365291 | Cluster 3 | Rps15 |

| | | | | | | | |
|---|---|---|---|---|---|---|---|
| Uqcrq | 2.57831980914925e-07 | 0.309001510083051 | 0.787 | 0.561 | 0.00800645650335117 | Cluster 3 | Uqcrq |
| Atp5g11 | 2.73953857836319e-07 | 0.505505628082609 | 0.561 | 0.246 | 0.00850708914739121 | Cluster 3 | Atp5g1 |
| Acadl1 | 2.97106360390794e-07 | 0.360522610355652 | 0.348 | 0.14 | 0.00922604380921532 | Cluster 3 | Acadl |
| Arpp19 | 3.37373490876535e-07 | 0.342118004194582 | 0.836 | 0.606 | 0.0104764590121891 | Cluster 3 | Arpp19 |
| Psmb2 | 3.55926905405631e-07 | 0.330399945601142 | 0.502 | 0.25 | 0.0110525981935611 | Cluster 3 | Psmb2 |
| Serbp11 | 3.68771113619473e-07 | 0.458996054848562 | 0.725 | 0.378 | 0.0114514493912255 | Cluster 3 | Serbp1 |
| Rpl211 | 3.92135773376903e-07 | 0.404013506070076 | 0.962 | 0.818 | 0.012176992170673 | Cluster 3 | Rpl21 |
| Lamtor21 | 4.24297386829599e-07 | 0.486452134148388 | 0.679 | 0.34 | 0.0131757067532196 | Cluster 3 | Lamtor2 |
| Cops9 | 4.27981580349973e-07 | 0.336261924536428 | 0.554 | 0.304 | 0.0132901120146077 | Cluster 3 | Cops9 |
| Eif3f | 4.34669960335505e-07 | 0.299364070617181 | 0.449 | 0.228 | 0.0134978062782985 | Cluster 3 | Eif3f |
| Rpl23 | 4.53487220533331e-07 | 0.315143927199873 | 0.983 | 0.915 | 0.0140821386592215 | Cluster 3 | Rpl23 |
| Snrpd21 | 5.0629256807888e-07 | 0.472273135423369 | 0.526 | 0.223 | 0.0157219031165535 | Cluster 3 | Snrpd2 |
| Psmb3 | 5.30213567518813e-07 | 0.469826825253194 | 0.756 | 0.407 | 0.0164647219121617 | Cluster 3 | Psmb3 |
| Mrps18c | 5.81247298345663e-07 | 0.329147787615061 | 0.348 | 0.148 | 0.0180494723555279 | Cluster 3 | Mrps18c |
| Atp5h1 | 5.89924985866762e-07 | 0.456561051329716 | 0.774 | 0.485 | 0.0183189405861206 | Cluster 3 | Atp5h |
| H2afy | 6.26419315351314e-07 | 0.374147209967615 | 0.46 | 0.229 | 0.0194521989996044 | Cluster 3 | H2afy |
| Psmb1 | 6.51919902587808e-07 | 0.326643277740071 | 0.526 | 0.262 | 0.0202440687350592 | Cluster 3 | Psmb1 |
| Emg11 | 6.78224716254812e-07 | 0.354828072167163 | 0.425 | 0.189 | 0.0210609121138607 | Cluster 3 | Emg1 |
| Ndufb8 | 7.01837099614972e-07 | 0.524055847642418 | 0.582 | 0.25 | 0.0217941474543437 | Cluster 3 | Ndufb8 |
| Lsm12 | 7.26932668175355e-07 | 0.402083821844638 | 0.449 | 0.174 | 0.0225734401448493 | Cluster 3 | Lsm12 |
| Sdha | 7.56731978323255e-07 | 0.298327980726927 | 0.394 | 0.168 | 0.023498798122872 | Cluster 3 | Sdha |
| Atp5g22 | 8.05355744697071e-07 | 0.368343148628753 | 0.589 | 0.313 | 0.0250087119400781 | Cluster 3 | Atp5g2 |
| Rpl111 | 8.1163905387604e-07 | 0.468233421326627 | 0.944 | 0.719 | 0.0252038275400127 | Cluster 3 | Rpl11 |
| Eif3c1 | 8.83395504421012e-07 | 0.369608014670727 | 0.432 | 0.189 | 0.0274320805987857 | Cluster 3 | Eif3c |
| Ndufa6 | 9.44544063532901e-07 | 0.405159322196391 | 0.589 | 0.304 | 0.0293309268048872 | Cluster 3 | Ndufa6 |
| 2010107E04Rik1 | 1.02461761044942e-06 | 0.432939638019496 | 0.815 | 0.54 | 0.0318174506572858 | Cluster 3 | 2010107E04Rik |
| Cox7c1 | 1.04096027581504e-06 | 0.468485495744293 | 0.54 | 0.354 | 0.0323249394448845 | Cluster 3 | Cox7c |
| Pin4 | 1.05275902061626e-06 | 0.319929592566518 | 0.348 | 0.138 | 0.0326913258671969 | Cluster 3 | Pin4 |
| Pdap1 | 1.08706383476544e-06 | 0.309672308031517 | 0.422 | 0.199 | 0.0337565932609712 | Cluster 3 | Pdap1 |
| Rpl17 | 1.14697055135946e-06 | 0.264080872269867 | 0.972 | 0.829 | 0.0356168765313654 | Cluster 3 | Rpl17 |
| Lsm7 | 1.15910865443627e-06 | 0.3607399405444 | 0.279 | 0.094 | 0.0359938010462093 | Cluster 3 | Lsm7 |
| Sfpq | 1.17586176653352e-06 | 0.33546891009373 | 0.693 | 0.436 | 0.0365140354361654 | Cluster 3 | Sfpq |
| Exoc4 | 1.30660367715348e-06 | 0.265550965190356 | 0.209 | 0.063 | 0.0405739639866469 | Cluster 3 | Exoc4 |
| Rpl41 | 1.34294254710959e-06 | 0.356301304743767 | 0.99 | 0.975 | 0.0417023949153942 | Cluster 3 | Rpl41 |
| Ccng11 | 1.34965656126722e-06 | 0.494740339033835 | 0.425 | 0.161 | 0.041910885197031 | Cluster 3 | Ccng1 |
| Dnttip2 | 1.38067569798762e-06 | 0.38227355346458 | 0.334 | 0.133 | 0.0428741224496096 | Cluster 3 | Dnttip2 |
| Sf3b51 | 1.43151547487478e-06 | 0.3646700476313 | 0.641 | 0.415 | 0.0444528500412865 | Cluster 3 | Sf3b5 |
| Gpx1 | 1.52564402907581e-06 | 0.366392822179458 | 0.972 | 0.861 | 0.0473758240348911 | Cluster 3 | Gpx1 |
| Rpl61 | 1.72580575257882e-06 | 0.460054529788295 | 0.868 | 0.637 | 0.05359144603483 | Cluster 3 | Rpl6 |
| Selenof1 | 1.77313029845037e-06 | 0.276624884068791 | 0.599 | 0.341 | 0.0550610151577795 | Cluster 3 | Selenof |
| Atp5k | 1.78418445166772e-06 | 0.378662249125676 | 0.547 | 0.28 | 0.0554042797776376 | Cluster 3 | Atp5k |
| Rpl37a | 1.82312683167658e-06 | 0.299504709445901 | 0.969 | 0.887 | 0.0566135575040529 | Cluster 3 | Rpl37a |
| Banf11 | 2.00424367437895e-06 | 0.467358780596838 | 0.498 | 0.199 | 0.0622377788204894 | Cluster 3 | Banf1 |
| Polr2f1 | 2.01366759802038e-06 | 0.380494780656078 | 0.557 | 0.273 | 0.0625304199213268 | Cluster 3 | Polr2f |
| Rpl7 | 2.14427870472113e-06 | 0.291293975061279 | 0.878 | 0.64 | 0.0665862866177053 | Cluster 3 | Rpl7 |
| Ran1 | 2.27489140647231e-06 | 0.37329749685483 | 0.662 | 0.347 | 0.0706422028451845 | Cluster 3 | Ran |
| Rpl131 | 2.29359401546223e-06 | 0.425268682274422 | 0.951 | 0.83 | 0.0712229749621487 | Cluster 3 | Rpl13 |
| Mrpl421 | 2.50521432310132e-06 | 0.442683134326506 | 0.418 | 0.16 | 0.0777944203752654 | Cluster 3 | Mrpl42 |
| Rwdd11 | 2.54068272022964e-06 | 0.36466201679554 | 0.467 | 0.21 | 0.0788958205112911 | Cluster 3 | Rwdd1 |
| Cox4i1 | 2.54965534895477e-06 | 0.322834531190702 | 0.927 | 0.763 | 0.0791744475510923 | Cluster 3 | Cox4i1 |
| Prpf8 | 2.64886081506016e-06 | 0.36793846626022 | 0.397 | 0.161 | 0.082255074890063 | Cluster 3 | Prpf8 |
| Samm50 | 2.65232825167933e-06 | 0.298497254464015 | 0.296 | 0.109 | 0.0823627491993981 | Cluster 3 | Samm50 |
| Cox6b1 | 2.76832419471194e-06 | 0.357377598911515 | 0.882 | 0.687 | 0.0859647712183899 | Cluster 3 | Cox6b1 |
| Kif5b1 | 2.81998535888906e-06 | 0.315359721088071 | 0.641 | 0.394 | 0.0875690053495821 | Cluster 3 | Kif5b |
| Ndufa14 | 2.84744086836743e-06 | 0.32014925617191 | 0.645 | 0.394 | 0.0884215812854137 | Cluster 3 | Ndufa1 |
| Snrpb2 | 2.89081358593509e-06 | 0.413001376409543 | 0.791 | 0.452 | 0.0897684342840423 | Cluster 3 | Snrpb |
| Hint11 | 3.11203780517963e-06 | 0.408571249663388 | 0.676 | 0.386 | 0.0966381099642431 | Cluster 3 | Hint1 |
| Lsm61 | 3.22535759155269e-06 | 0.45215339639173 | 0.46 | 0.166 | 0.100157029290486 | Cluster 3 | Lsm6 |
| Atp5e | 3.24479458023979e-06 | 0.352664712321241 | 0.948 | 0.814 | 0.100760606100186 | Cluster 3 | Atp5e |
| Psma1 | 3.2452858961322e-06 | 0.318266108565579 | 0.53 | 0.309 | 0.100775862932593 | Cluster 3 | Psma1 |
| Npm11 | 3.25460648482156e-06 | 0.491375647304474 | 0.509 | 0.201 | 0.101065295173164 | Cluster 3 | Npm1 |

| | | | | | | | |
|---|---|---|---|---|---|---|---|
| Hnrnpf1 | 3.26179404183196e-06 | 0.309959470344304 | 0.934 | 0.777 | 0.101288490381008 | Cluster 3 | Hnrnpf |
| Nr4a2 | 3.26774940070593e-06 | 0.588513732381008 | 0.366 | 0.204 | 0.101473422140121 | Cluster 3 | Nr4a2 |
| Rpl36a | 3.30153762354761e-06 | 0.400649176620771 | 0.638 | 0.353 | 0.102522647824024 | Cluster 3 | Rpl36a |
| Hnrnpdl1 | 3.39014340648548e-06 | 0.3069599922412 | 0.575 | 0.302 | 0.105274123201594 | Cluster 3 | Hnrnpdl |
| Atp5a1 | 3.41073311286343e-06 | 0.356010192595556 | 0.613 | 0.343 | 0.105913495353748 | Cluster 3 | Atp5a1 |
| Rpl24 | 3.79251178088769e-06 | 0.406158961870552 | 0.854 | 0.641 | 0.117768868331906 | Cluster 3 | Rpl24 |
| Rps27l1 | 3.92569055538054e-06 | 0.585331773450397 | 0.801 | 0.486 | 0.121904468816232 | Cluster 3 | Rps27l |
| Ctsa | 3.96901624671609e-06 | 0.267454499314329 | 0.683 | 0.432 | 0.123249861509275 | Cluster 3 | Ctsa |
| Tsn | 3.9734030887482e-06 | 0.338327840773141 | 0.334 | 0.145 | 0.123386086114898 | Cluster 3 | Tsn |
| Atp5j21 | 4.1200700570151e-06 | 0.423758217102811 | 0.739 | 0.475 | 0.12794053548049 | Cluster 3 | Atp5j2 |
| Rpl9 | 4.12209352841741e-06 | 0.256460667721274 | 0.965 | 0.845 | 0.128003370337946 | Cluster 3 | Rpl9 |
| Rpl7l1 | 4.40328361452987e-06 | 0.341304029655973 | 0.352 | 0.135 | 0.136735166081996 | Cluster 3 | Rpl7l1 |
| Rps15a1 | 4.67913021267167e-06 | 0.438941234382501 | 0.951 | 0.757 | 0.145301030494093 | Cluster 3 | Rps15a |
| Spcs1 | 5.10466860057042e-06 | 0.322642814517508 | 0.463 | 0.242 | 0.158515274053513 | Cluster 3 | Spcs1 |
| Rpl51 | 5.29735308138547e-06 | 0.443346072404685 | 0.798 | 0.517 | 0.164498705236263 | Cluster 3 | Rpl5 |
| Rpl39 | 5.30776492559063e-06 | 0.30082961591395 | 0.948 | 0.791 | 0.164822024234366 | Cluster 3 | Rpl39 |
| N4bp2l2 | 5.35278846614811e-06 | 0.33434179265209 | 0.484 | 0.275 | 0.166220140239297 | Cluster 3 | N4bp2l2 |
| Pdcd6 | 6.22837587863904e-06 | 0.301923844048769 | 0.537 | 0.328 | 0.193409756159378 | Cluster 3 | Pdcd6 |
| Eif3m1 | 6.57207659809003e-06 | 0.396619714684786 | 0.383 | 0.141 | 0.20408269460049 | Cluster 3 | Eif3m |
| Rpl122 | 7.4386497648409e-06 | 0.394698500494447 | 0.84 | 0.575 | 0.230992391147605 | Cluster 3 | Rpl12 |
| Snw1 | 7.83417584713975e-06 | 0.362524640119628 | 0.404 | 0.167 | 0.243274662581231 | Cluster 3 | Snw1 |
| Rcc2 | 8.75758540971507e-06 | 0.316018524656469 | 0.282 | 0.084 | 0.271949299727882 | Cluster 3 | Rcc2 |
| Ube2e1 | 8.98713731487814e-06 | 0.288193311604996 | 0.366 | 0.161 | 0.279077575038911 | Cluster 3 | Ube2e1 |
| Ghitm | 1.0164787409925e-05 | 0.283665805114487 | 0.345 | 0.169 | 0.3156471434404 | Cluster 3 | Ghitm |
| Rps25 | 1.01700747664172e-05 | 0.324534118081614 | 0.969 | 0.837 | 0.315811331721553 | Cluster 3 | Rps25 |
| Cnot1 | 1.03565994173011e-05 | 0.284858304416372 | 0.408 | 0.191 | 0.321603481705452 | Cluster 3 | Cnot1 |
| Cuta | 1.04130149318916e-05 | 0.335771249941877 | 0.418 | 0.189 | 0.323355352680031 | Cluster 3 | Cuta |
| Fubp11 | 1.04659724582035e-05 | 0.40845355286283 | 0.474 | 0.23 | 0.324999842744595 | Cluster 3 | Fubp1 |
| Ndufc11 | 1.17936951497133e-05 | 0.349169755248665 | 0.411 | 0.173 | 0.366229615484047 | Cluster 3 | Ndufc1 |
| Psmc31 | 1.20296016065117e-05 | 0.404976248824686 | 0.38 | 0.146 | 0.373555218687006 | Cluster 3 | Psmc3 |
| Pvr | 1.26878118377939e-05 | 0.255711803365631 | 0.174 | 0.052 | 0.393994620999013 | Cluster 3 | Pvr |
| Uqcrc2 | 1.34814098981866e-05 | 0.346339517736889 | 0.317 | 0.115 | 0.418638221568388 | Cluster 3 | Uqcrc2 |
| Cox6c | 1.34880278891781e-05 | 0.500378979712176 | 0.725 | 0.47 | 0.418843730042647 | Cluster 3 | Cox6c |
| Set1 | 1.36425987517059e-05 | 0.435791087169426 | 0.599 | 0.292 | 0.423643619036724 | Cluster 3 | Set |
| Csnk2b | 1.37544675414108e-05 | 0.290003185230643 | 0.498 | 0.284 | 0.427117480563431 | Cluster 3 | Csnk2b |
| Higd2a | 1.38476062650258e-05 | 0.296253816889329 | 0.46 | 0.26 | 0.430009717347845 | Cluster 3 | Higd2a |
| Rps7 | 1.44542897802265e-05 | 0.313167098367051 | 0.85 | 0.682 | 0.448849060545374 | Cluster 3 | Rps7 |
| Mrfap1 | 1.5027488052415e-05 | 0.337725222696131 | 0.676 | 0.421 | 0.466648586491644 | Cluster 3 | Mrfap1 |
| Cct81 | 1.53049083593756e-05 | 0.348706611223705 | 0.491 | 0.242 | 0.47526331928369 | Cluster 3 | Cct8 |
| Psmc51 | 1.54601493961688e-05 | 0.309664225666976 | 0.348 | 0.13 | 0.480084019199231 | Cluster 3 | Psmc5 |
| B3gnt5 | 1.62335962182732e-05 | 0.302029448927006 | 0.244 | 0.117 | 0.504101863366037 | Cluster 3 | B3gnt5 |
| Uqcrc1 | 1.7473420057701e-05 | 0.347377072315572 | 0.373 | 0.155 | 0.542602113051789 | Cluster 3 | Uqcrc1 |
| Hnrnpd1 | 1.82703018297011e-05 | 0.402801910126039 | 0.627 | 0.343 | 0.567347682717709 | Cluster 3 | Hnrnpd |
| Rps3a1 | 1.82925221230227e-05 | 0.346234284413026 | 0.976 | 0.835 | 0.568037689486223 | Cluster 3 | Rps3a1 |
| Eif3e | 1.95243456496144e-05 | 0.307315130026381 | 0.289 | 0.106 | 0.606289505457476 | Cluster 3 | Eif3e |
| Rpl31 | 1.99503183711323e-05 | 0.37671123625704 | 0.655 | 0.406 | 0.61951723637877 | Cluster 3 | Rpl31 |
| Nsmce4a | 2.01458265904962e-05 | 0.306453331751513 | 0.366 | 0.14 | 0.625588353114677 | Cluster 3 | Nsmce4a |
| Zcrb1 | 2.05668434704454e-05 | 0.253143203449499 | 0.408 | 0.223 | 0.638662190287742 | Cluster 3 | Zcrb1 |
| Hspe11 | 2.14130668624048e-05 | 0.290515586995011 | 0.334 | 0.14 | 0.664939965278256 | Cluster 3 | Hspe1 |
| Psmd11 | 2.15037518113613e-05 | 0.312435301952988 | 0.366 | 0.207 | 0.667756004998204 | Cluster 3 | Psmd11 |
| Psmc4 | 2.19652694154643e-05 | 0.275862666048998 | 0.314 | 0.142 | 0.682087511158413 | Cluster 3 | Psmc4 |
| Mrpl30 | 2.21837016765938e-05 | 0.308881551090913 | 0.38 | 0.18 | 0.688870488163267 | Cluster 3 | Mrpl30 |
| Uqcrb1 | 2.24420574773338e-05 | 0.426577507273225 | 0.53 | 0.261 | 0.696893210843647 | Cluster 3 | Uqcrb |
| Rpl141 | 2.3570318535656e-05 | 0.33780145843435 | 0.801 | 0.531 | 0.731929101487725 | Cluster 3 | Rpl14 |
| Eprs1 | 2.41631020412992e-05 | 0.291885038291568 | 0.247 | 0.111 | 0.750336807688463 | Cluster 3 | Eprs |
| Zcchc17 | 2.56896454597774e-05 | 0.266598608446674 | 0.251 | 0.089 | 0.797740560462467 | Cluster 3 | Zcchc17 |
| Rabac1 | 2.63672635669633e-05 | 0.328482451643821 | 0.84 | 0.631 | 0.81878263554491 | Cluster 3 | Rabac1 |
| Purb | 2.64676166919227e-05 | 0.276194371583288 | 0.669 | 0.455 | 0.821898901134275 | Cluster 3 | Purb |
| Rpl36al2 | 2.65887287741776e-05 | 0.340804553114366 | 0.676 | 0.392 | 0.825659794624536 | Cluster 3 | Rpl36al |
| Sec11c | 2.82127654627565e-05 | 0.296252880427682 | 0.547 | 0.305 | 0.876091005914976 | Cluster 3 | Sec11c |
| Snrpd32 | 2.86574385358396e-05 | 0.388039184054112 | 0.505 | 0.251 | 0.889899438853426 | Cluster 3 | Snrpd3 |

| | | | | | | | |
|---|---|---|---|---|---|---|---|
| Emc62 | 2.9138544590741e-05 | 0.287942769868288 | 0.484 | 0.264 | 0.90483922517628 | Cluster 3 | Emc6 |
| Ndufb61 | 2.9383527281674e-05 | 0.328460308386937 | 0.415 | 0.221 | 0.912446672677824 | Cluster 3 | Ndufb6 |
| Slc25a5 | 3.11106962132063e-05 | 0.282739240907613 | 0.631 | 0.401 | 0.966080449508694 | Cluster 3 | Slc25a5 |
| Top1 | 3.26968391781694e-05 | 0.289727330265005 | 0.714 | 0.494 | 1 | Cluster 3 | Top1 |
| Exosc5 | 3.28846964076036e-05 | 0.310957695640598 | 0.345 | 0.14 | 1 | Cluster 3 | Exosc5 |
| Bzw1 | 3.35840478868115e-05 | 0.306728638531088 | 0.669 | 0.424 | 1 | Cluster 3 | Bzw1 |
| Tma7 | 3.41174953276793e-05 | 0.4074156757881 | 0.787 | 0.53 | 1 | Cluster 3 | Tma7 |
| Hist1h1c | 3.7766384382648e-05 | 0.600359501043774 | 0.505 | 0.226 | 1 | Cluster 3 | Hist1h1c |
| Lsm21 | 3.78546725891719e-05 | 0.295473480422584 | 0.303 | 0.114 | 1 | Cluster 3 | Lsm2 |
| Mrps33 | 3.80443875208463e-05 | 0.294101390208302 | 0.491 | 0.264 | 1 | Cluster 3 | Mrps33 |
| Snrpf1 | 4.01697730282545e-05 | 0.473470748483191 | 0.432 | 0.165 | 1 | Cluster 3 | Snrpf |
| mt-Nd3 | 4.15227930464472e-05 | 0.32311191029999 | 0.718 | 0.525 | 1 | Cluster 3 | mt-Nd3 |
| Nop102 | 4.17992979434064e-05 | 0.454127309799749 | 0.571 | 0.288 | 1 | Cluster 3 | Nop10 |
| Chchd11 | 4.18750446497105e-05 | 0.372240612386663 | 0.432 | 0.165 | 1 | Cluster 3 | Chchd1 |
| Lyl1 | 4.38901707194919e-05 | 0.371289733844062 | 0.254 | 0.069 | 1 | Cluster 3 | Lyl1 |
| Srp14 | 4.41307802280832e-05 | 0.340376144188908 | 0.735 | 0.508 | 1 | Cluster 3 | Srp14 |
| Ndufab1 | 4.49965532411349e-05 | 0.327031390401573 | 0.324 | 0.13 | 1 | Cluster 3 | Ndufab1 |
| Mrpl21 | 4.5524191478271e-05 | 0.307552015648876 | 0.237 | 0.064 | 1 | Cluster 3 | Mrpl21 |
| Cox5a1 | 4.58267576803775e-05 | 0.392012260410467 | 0.76 | 0.455 | 1 | Cluster 3 | Cox5a |
| Rex1bd1 | 4.58772128963627e-05 | 0.300687912459226 | 0.334 | 0.135 | 1 | Cluster 3 | Rex1bd |
| Mrpl522 | 4.68282333045423e-05 | 0.390576778688701 | 0.777 | 0.524 | 1 | Cluster 3 | Mrpl52 |
| Ptgs12 | 4.78775359911719e-05 | 0.372138333380761 | 0.53 | 0.286 | 1 | Cluster 3 | Ptgs1 |
| Vdac2 | 4.92619832974144e-05 | 0.279206205618145 | 0.641 | 0.42 | 1 | Cluster 3 | Vdac2 |
| Rps10 | 4.9765488520755e-05 | 0.265815413692621 | 0.965 | 0.854 | 1 | Cluster 3 | Rps10 |
| Mrpl32 | 5.31752825466537e-05 | 0.317485807641543 | 0.286 | 0.095 | 1 | Cluster 3 | Mrpl32 |
| Hnrnpl | 5.4304080052168e-05 | 0.303328618179345 | 0.331 | 0.146 | 1 | Cluster 3 | Hnrnpl |
| Man2b1 | 5.67909815086947e-05 | 0.377065697300448 | 0.627 | 0.409 | 1 | Cluster 3 | Man2b1 |
| Arglu1 | 5.71366282737114e-05 | 0.262739491005159 | 0.376 | 0.21 | 1 | Cluster 3 | Arglu1 |
| Grpel1 | 5.81714592329929e-05 | 0.291756195546562 | 0.39 | 0.199 | 1 | Cluster 3 | Grpel1 |
| Ddx46 | 5.84235244588605e-05 | 0.309858446234958 | 0.338 | 0.16 | 1 | Cluster 3 | Ddx46 |
| Cysltr1 | 5.86481753041225e-05 | 0.291889409665939 | 0.279 | 0.136 | 1 | Cluster 3 | Cysltr1 |
| Fam104a | 5.91562099603119e-05 | 0.286828955850274 | 0.547 | 0.36 | 1 | Cluster 3 | Fam104a |
| Rpf1 | 6.02293288671558e-05 | 0.268009949707914 | 0.192 | 0.045 | 1 | Cluster 3 | Rpf1 |
| Eapp | 6.51780026192039e-05 | 0.273639336958146 | 0.46 | 0.252 | 1 | Cluster 3 | Eapp |
| Mbnl1 | 6.7383521265774e-05 | 0.319765194058626 | 0.589 | 0.355 | 1 | Cluster 3 | Mbnl1 |
| Med29 | 6.78980053949125e-05 | 0.270390203755076 | 0.244 | 0.085 | 1 | Cluster 3 | Med29 |
| Sod1 | 6.83933988078167e-05 | 0.265825902900296 | 0.359 | 0.182 | 1 | Cluster 3 | Sod1 |
| Timm13 | 7.40527022554821e-05 | 0.267601583714876 | 0.387 | 0.182 | 1 | Cluster 3 | Timm13 |
| Cox5b1 | 7.44430875710788e-05 | 0.291285324664964 | 0.798 | 0.613 | 1 | Cluster 3 | Cox5b |
| Ndufa3 | 7.52607911932677e-05 | 0.310570423435587 | 0.624 | 0.386 | 1 | Cluster 3 | Ndufa3 |
| Naca1 | 7.68536267350441e-05 | 0.394530740299975 | 0.826 | 0.547 | 1 | Cluster 3 | Naca |
| Nol71 | 7.82346888246313e-05 | 0.456533697277708 | 0.46 | 0.16 | 1 | Cluster 3 | Nol7 |
| Zkscan3 | 7.90554745476708e-05 | 0.2760004175947 | 0.146 | 0.053 | 1 | Cluster 3 | Zkscan3 |
| Rpl7a1 | 8.00535331586134e-05 | 0.376518523906954 | 0.697 | 0.436 | 1 | Cluster 3 | Rpl7a |
| BC005624 | 8.22944981289685e-05 | 0.25622813531652 | 0.537 | 0.301 | 1 | Cluster 3 | BC005624 |
| Fkbp4 | 8.42193679431968e-05 | 0.259466988525974 | 0.178 | 0.051 | 1 | Cluster 3 | Fkbp4 |
| Nucks1 | 8.65620593815655e-05 | 0.28824458113424 | 0.275 | 0.103 | 1 | Cluster 3 | Nucks1 |
| Atp5b1 | 8.99000351647023e-05 | 0.385750201134615 | 0.819 | 0.53 | 1 | Cluster 3 | Atp5b |
| Dbi1 | 9.00466225820002e-05 | 0.405616493466898 | 0.463 | 0.223 | 1 | Cluster 3 | Dbi |
| Nedd8 | 9.29381667096393e-05 | 0.320695564650908 | 0.801 | 0.561 | 1 | Cluster 3 | Nedd8 |
| Ddx21 | 9.88160688718229e-05 | 0.31201292028553 | 0.334 | 0.135 | 1 | Cluster 3 | Ddx21 |
| Odc1 | 0.000102494584616222 | 0.471773880672977 | 0.481 | 0.278 | 1 | Cluster 3 | Odc1 |
| Matr3 | 0.000107075695788769 | 0.336221697835719 | 0.488 | 0.257 | 1 | Cluster 3 | Matr3 |
| Lsm8 | 0.000107450804828714 | 0.250366828936244 | 0.317 | 0.141 | 1 | Cluster 3 | Lsm8 |
| Polr2k | 0.000116887091201106 | 0.29147094762759 | 0.463 | 0.259 | 1 | Cluster 3 | Polr2k |
| Hsp90ab1 | 0.000117545494488022 | 0.421373111716953 | 0.725 | 0.468 | 1 | Cluster 3 | Hsp90ab1 |
| Ube2k | 0.000128032781221549 | 0.252301906212032 | 0.443 | 0.258 | 1 | Cluster 3 | Ube2k |
| Rps14 | 0.000130405351114281 | 0.370025880139186 | 0.965 | 0.836 | 1 | Cluster 3 | Rps14 |
| Mrpl571 | 0.000130792544901454 | 0.354906276988517 | 0.432 | 0.219 | 1 | Cluster 3 | Mrpl57 |
| Dynlrb11 | 0.000133605817027241 | 0.310306465618269 | 0.571 | 0.326 | 1 | Cluster 3 | Dynlrb1 |
| Zranb2 | 0.000140194290703095 | 0.287493577705954 | 0.376 | 0.199 | 1 | Cluster 3 | Zranb2 |

| | | | | | | | |
|---|---|---|---|---|---|---|---|
| Snrpd11 | 0.000141879379218113 | 0.27784866093576 | 0.307 | 0.132 | 1 | Cluster 3 | Snrpd1 |
| Rbm42 | 0.000142098196345145 | 0.343550562562645 | 0.467 | 0.243 | 1 | Cluster 3 | Rbm42 |
| Atp5c1 | 0.000142563010886259 | 0.333881521000813 | 0.571 | 0.347 | 1 | Cluster 3 | Atp5c1 |
| Anp32b | 0.000142753363609807 | 0.327529749563765 | 0.564 | 0.309 | 1 | Cluster 3 | Anp32b |
| Ptpn18 | 0.000144117471282392 | 0.356958487139072 | 0.582 | 0.365 | 1 | Cluster 3 | Ptpn18 |
| Sptssa | 0.000147994456213059 | 0.270255589986728 | 0.303 | 0.113 | 1 | Cluster 3 | Sptssa |
| Rpl10 | 0.000148778249866886 | 0.280428823083561 | 0.951 | 0.814 | 1 | Cluster 3 | Rpl10 |
| Atp5l | 0.000149089336357812 | 0.267352634157557 | 0.92 | 0.8 | 1 | Cluster 3 | Atp5l |
| Hdgf | 0.000157119558852143 | 0.273338468273573 | 0.394 | 0.188 | 1 | Cluster 3 | Hdgf |
| Cwc15 | 0.000157838381685821 | 0.401562875347696 | 0.46 | 0.215 | 1 | Cluster 3 | Cwc15 |
| Tmem208 | 0.000161206490321568 | 0.299440005701306 | 0.404 | 0.185 | 1 | Cluster 3 | Tmem208 |
| Cenpx1 | 0.000170202465171216 | 0.416357844757254 | 0.401 | 0.154 | 1 | Cluster 3 | Cenpx |
| Lsm51 | 0.000174522094795148 | 0.41541229598077 | 0.394 | 0.14 | 1 | Cluster 3 | Lsm5 |
| Ndufa21 | 0.000175185170395695 | 0.432921089171097 | 0.753 | 0.501 | 1 | Cluster 3 | Ndufa2 |
| Rpl27 | 0.000175286935715839 | 0.308866049121117 | 0.77 | 0.529 | 1 | Cluster 3 | Rpl27 |
| Copb1 | 0.000182714564220874 | 0.262711551434499 | 0.307 | 0.151 | 1 | Cluster 3 | Copb1 |
| Tomm22 | 0.000182930955551013 | 0.301657526627355 | 0.666 | 0.395 | 1 | Cluster 3 | Tomm22 |
| Swi5 | 0.000184759607062991 | 0.272961532972506 | 0.397 | 0.205 | 1 | Cluster 3 | Swi5 |
| Trmt1121 | 0.000186517005169387 | 0.297670180690768 | 0.571 | 0.313 | 1 | Cluster 3 | Trmt112 |
| Rpl3 | 0.000187160711365841 | 0.424718037940599 | 0.693 | 0.386 | 1 | Cluster 3 | Rpl3 |
| Etfb | 0.000194051232262861 | 0.370228155821667 | 0.376 | 0.147 | 1 | Cluster 3 | Etfb |
| Ift20 | 0.000194362834997822 | 0.271045620984317 | 0.366 | 0.166 | 1 | Cluster 3 | Ift20 |
| Eif4h | 0.000198865762557011 | 0.251365150379004 | 0.512 | 0.292 | 1 | Cluster 3 | Eif4h |
| Tomm20 | 0.000199373076608877 | 0.273105703172952 | 0.61 | 0.38 | 1 | Cluster 3 | Tomm20 |
| Dek1 | 0.000205043065131674 | 0.397932336808759 | 0.408 | 0.167 | 1 | Cluster 3 | Dek |
| Polr2m | 0.000213804448281658 | 0.302142942849059 | 0.24 | 0.072 | 1 | Cluster 3 | Polr2m |
| Usmg5 | 0.000218588885836413 | 0.322330807789388 | 0.787 | 0.593 | 1 | Cluster 3 | Usmg5 |
| Strap | 0.000220843721246878 | 0.270209276624828 | 0.467 | 0.266 | 1 | Cluster 3 | Strap |
| Ddost1 | 0.00022965461868527 | 0.302874581734214 | 0.289 | 0.128 | 1 | Cluster 3 | Ddost |
| Cd812 | 0.000238406507094447 | 0.35663325288614 | 0.446 | 0.242 | 1 | Cluster 3 | Cd81 |
| Ndufs51 | 0.00024843263072174 | 0.369902780118171 | 0.564 | 0.279 | 1 | Cluster 3 | Ndufs5 |
| Mpc2 | 0.000258611277833376 | 0.343612962217502 | 0.578 | 0.335 | 1 | Cluster 3 | Mpc2 |
| Use1 | 0.000270377689098868 | 0.256162641374376 | 0.38 | 0.222 | 1 | Cluster 3 | Use1 |
| Atg5 | 0.000272507475495404 | 0.267084951237905 | 0.286 | 0.131 | 1 | Cluster 3 | Atg5 |
| Ltc4s1 | 0.000278572148038136 | 0.529037543792619 | 0.345 | 0.109 | 1 | Cluster 3 | Ltc4s |
| Papola | 0.000279121781232728 | 0.330641950748354 | 0.456 | 0.242 | 1 | Cluster 3 | Papola |
| Cst31 | 0.000280434511681669 | 0.367830524709878 | 0.986 | 0.936 | 1 | Cluster 3 | Cst3 |
| Krit1 | 0.000280698251855286 | 0.254723279615665 | 0.272 | 0.135 | 1 | Cluster 3 | Krit1 |
| Sec62 | 0.00028933670254912 | 0.270217573023431 | 0.641 | 0.388 | 1 | Cluster 3 | Sec62 |
| 1110004F10Rik | 0.000301551843974231 | 0.358679998232848 | 0.328 | 0.128 | 1 | Cluster 3 | 1110004F10Rik |
| Ilf2 | 0.00030512337643355 | 0.411165416797415 | 0.293 | 0.081 | 1 | Cluster 3 | Ilf2 |
| Nudc | 0.000308326284801354 | 0.331947548856036 | 0.38 | 0.157 | 1 | Cluster 3 | Nudc |
| Eif2s2 | 0.000308805945317363 | 0.319746740952218 | 0.516 | 0.303 | 1 | Cluster 3 | Eif2s2 |
| Chtop | 0.000310272697435353 | 0.250693532348868 | 0.348 | 0.174 | 1 | Cluster 3 | Chtop |
| Psma6 | 0.000311549985062521 | 0.283344091238595 | 0.376 | 0.183 | 1 | Cluster 3 | Psma6 |
| Sumo2 | 0.000316394905261526 | 0.302025043634074 | 0.784 | 0.541 | 1 | Cluster 3 | Sumo2 |
| Nsf | 0.000318282546809846 | 0.283940620148768 | 0.432 | 0.239 | 1 | Cluster 3 | Nsf |
| Reep51 | 0.000322293548494034 | 0.311659264301126 | 0.756 | 0.547 | 1 | Cluster 3 | Reep5 |
| Mtch2 | 0.000324210321027197 | 0.328548635546853 | 0.38 | 0.173 | 1 | Cluster 3 | Mtch2 |
| Usp16 | 0.000329696428700215 | 0.257535108500471 | 0.47 | 0.304 | 1 | Cluster 3 | Usp16 |
| Ppig1 | 0.000338055135663352 | 0.421618595082231 | 0.502 | 0.244 | 1 | Cluster 3 | Ppig |
| Psmc6 | 0.000345780839453043 | 0.295097406569266 | 0.376 | 0.16 | 1 | Cluster 3 | Psmc6 |
| Ndufb9 | 0.000346289325163016 | 0.402443125671192 | 0.728 | 0.477 | 1 | Cluster 3 | Ndufb9 |
| Zfhx3 | 0.000365203757561112 | 0.274920860521295 | 0.338 | 0.192 | 1 | Cluster 3 | Zfhx3 |
| Prkar1a | 0.00037070464742143 | 0.280385990649576 | 0.732 | 0.496 | 1 | Cluster 3 | Prkar1a |
| Phb2 | 0.000373857031587218 | 0.273413655638468 | 0.307 | 0.122 | 1 | Cluster 3 | Phb2 |
| Fbl1 | 0.000381022524698757 | 0.29013498071642 | 0.303 | 0.117 | 1 | Cluster 3 | Fbl |
| Rasgef1b1 | 0.00038694753651359 | 0.406608117150607 | 0.226 | 0.093 | 1 | Cluster 3 | Rasgef1b |
| Park72 | 0.000388587385083316 | 0.427582789750773 | 0.61 | 0.34 | 1 | Cluster 3 | Park7 |
| Psma2 | 0.000402707387517823 | 0.360929234262037 | 0.61 | 0.355 | 1 | Cluster 3 | Psma2 |
| Eif3g | 0.00041804168515147 | 0.368525700493678 | 0.429 | 0.18 | 1 | Cluster 3 | Eif3g |

| | | | | | | | |
|---|---|---|---|---|---|---|---|
| Snrpc | 0.000422630368021229 | 0.278694686552917 | 0.47 | 0.29 | 1 | Cluster 3 | Snrpc |
| Ddx39 | 0.000430189532455684 | 0.282395595887899 | 0.366 | 0.141 | 1 | Cluster 3 | Ddx39 |
| Trappc6b | 0.000440433901107497 | 0.263362084415282 | 0.418 | 0.212 | 1 | Cluster 3 | Trappc6b |
| Rps6 | 0.000462509788778603 | 0.386273249863824 | 0.854 | 0.589 | 1 | Cluster 3 | Rps6 |
| Nmt1 | 0.000490716708449073 | 0.27709905871338 | 0.272 | 0.117 | 1 | Cluster 3 | Nmt1 |
| Eif6 | 0.000497987328157882 | 0.280196029254753 | 0.408 | 0.208 | 1 | Cluster 3 | Eif6 |
| Magoh | 0.000502862335431117 | 0.290152051230574 | 0.477 | 0.271 | 1 | Cluster 3 | Magoh |
| Eif4a31 | 0.00051407468839966 | 0.402888166902465 | 0.449 | 0.179 | 1 | Cluster 3 | Eif4a3 |
| mt-Nd5 | 0.000517291892682601 | 0.32230696094955 | 0.498 | 0.28 | 1 | Cluster 3 | mt-Nd5 |
| Nsrp1 | 0.000550532328135321 | 0.308459942498135 | 0.341 | 0.204 | 1 | Cluster 3 | Nsrp1 |
| Plekhj1 | 0.000560180903186379 | 0.277139941583545 | 0.491 | 0.271 | 1 | Cluster 3 | Plekhj1 |
| Rbm8a1 | 0.000570204614456753 | 0.355976849013674 | 0.62 | 0.327 | 1 | Cluster 3 | Rbm8a |
| Rbm25 | 0.00058204447077455 | 0.288886857098927 | 0.864 | 0.638 | 1 | Cluster 3 | Rbm25 |
| Gtf2h5 | 0.000603582371704442 | 0.261413695950382 | 0.502 | 0.292 | 1 | Cluster 3 | Gtf2h5 |
| Thrap31 | 0.000605943198377353 | 0.331629424970399 | 0.383 | 0.187 | 1 | Cluster 3 | Thrap3 |
| Ndufa10 | 0.000606761683123929 | 0.267735900553493 | 0.254 | 0.093 | 1 | Cluster 3 | Ndufa10 |
| Yy1 | 0.000606978856020358 | 0.274248593900066 | 0.495 | 0.299 | 1 | Cluster 3 | Yy1 |
| Txnip | 0.000626985423752463 | 0.457841012494409 | 0.746 | 0.592 | 1 | Cluster 3 | Txnip |
| Ttc14 | 0.000649284164821096 | 0.289542324859491 | 0.348 | 0.196 | 1 | Cluster 3 | Ttc14 |
| Eif3h1 | 0.000649860141037863 | 0.283440137615831 | 0.502 | 0.295 | 1 | Cluster 3 | Eif3h |
| Eci1 | 0.000651231034053549 | 0.291536913511136 | 0.209 | 0.062 | 1 | Cluster 3 | Eci1 |
| Pabpc1 | 0.000660206826610018 | 0.294757548264956 | 0.85 | 0.694 | 1 | Cluster 3 | Pabpc1 |
| Ranbp1 | 0.000675519478960171 | 0.321359344503937 | 0.366 | 0.14 | 1 | Cluster 3 | Ranbp1 |
| Ywhae1 | 0.000684223595010034 | 0.346674113575586 | 0.732 | 0.446 | 1 | Cluster 3 | Ywhae |
| Prpf40a1 | 0.000684813622901578 | 0.291798986978736 | 0.596 | 0.337 | 1 | Cluster 3 | Prpf40a |
| Cdk11b | 0.000694076205116633 | 0.331254667338636 | 0.537 | 0.311 | 1 | Cluster 3 | Cdk11b |
| Ssbp3 | 0.000715818667318092 | 0.33997379581192 | 0.293 | 0.104 | 1 | Cluster 3 | Ssbp3 |
| Pin1 | 0.000718116478539486 | 0.274954575497656 | 0.324 | 0.141 | 1 | Cluster 3 | Pin1 |
| Eif2s11 | 0.000742061531671375 | 0.294051918174229 | 0.321 | 0.123 | 1 | Cluster 3 | Eif2s1 |
| Hspa14 | 0.00076257470018606 | 0.283810420896104 | 0.293 | 0.119 | 1 | Cluster 3 | Hspa14 |
| Uqcc22 | 0.000779824432048112 | 0.287095589644451 | 0.408 | 0.196 | 1 | Cluster 3 | Uqcc2 |
| Ube2i1 | 0.000786022918940497 | 0.265564646543219 | 0.812 | 0.597 | 1 | Cluster 3 | Ube2i |
| Nipa2 | 0.0007868769261251 | 0.287229610257323 | 0.279 | 0.12 | 1 | Cluster 3 | Nipa2 |
| Elob | 0.000791402064387725 | 0.294222474321922 | 0.941 | 0.8 | 1 | Cluster 3 | Elob |
| Uqcr11 | 0.000793830048756738 | 0.325018187717592 | 0.627 | 0.411 | 1 | Cluster 3 | Uqcr11 |
| Cycs | 0.000805799841158153 | 0.280893921930344 | 0.523 | 0.294 | 1 | Cluster 3 | Cycs |
| 2410015M20Rik | 0.000818938502515201 | 0.332503380289666 | 0.439 | 0.216 | 1 | Cluster 3 | 2410015M20Rik |
| Ndufs72 | 0.000821174659658533 | 0.364640441467847 | 0.585 | 0.304 | 1 | Cluster 3 | Ndufs7 |
| Lamtor1 | 0.000822125631470494 | 0.25998178670403 | 0.568 | 0.406 | 1 | Cluster 3 | Lamtor1 |
| Llph1 | 0.000830574036501251 | 0.366632197863369 | 0.519 | 0.257 | 1 | Cluster 3 | Llph |
| Ndufb4 | 0.000839413197008437 | 0.374147736832782 | 0.47 | 0.208 | 1 | Cluster 3 | Ndufb4 |
| Grk6 | 0.000840714268052133 | 0.266459754992314 | 0.307 | 0.124 | 1 | Cluster 3 | Grk6 |
| Ndufa5 | 0.000878647700007022 | 0.297620625225905 | 0.181 | 0.038 | 1 | Cluster 3 | Ndufa5 |
| Hprt | 0.000883617474014449 | 0.302525183750108 | 0.415 | 0.196 | 1 | Cluster 3 | Hprt |
| Ncl1 | 0.00089081266369686 | 0.312433564340594 | 0.662 | 0.427 | 1 | Cluster 3 | Ncl |
| Tmed91 | 0.000899263581941281 | 0.329858147883378 | 0.537 | 0.305 | 1 | Cluster 3 | Tmed9 |
| Rbm3 | 0.000922838249467595 | 0.363189639596851 | 0.909 | 0.721 | 1 | Cluster 3 | Rbm3 |
| Eif4g2 | 0.000954368951082202 | 0.286901200178147 | 0.868 | 0.67 | 1 | Cluster 3 | Eif4g2 |
| Eif3i | 0.000962560896492951 | 0.270734020377734 | 0.286 | 0.127 | 1 | Cluster 3 | Eif3i |
| Eif3k1 | 0.000996202245249389 | 0.411414111470954 | 0.568 | 0.293 | 1 | Cluster 3 | Eif3k |
| Fkbp31 | 0.00102153255062838 | 0.334946041217679 | 0.317 | 0.122 | 1 | Cluster 3 | Fkbp3 |
| Itgax | 0.00102180397783966 | 0.255404152744837 | 0.439 | 0.262 | 1 | Cluster 3 | Itgax |
| Senp6 | 0.00105907129594938 | 0.288813436021667 | 0.425 | 0.22 | 1 | Cluster 3 | Senp6 |
| Denr | 0.00106908314788373 | 0.301008335522276 | 0.481 | 0.26 | 1 | Cluster 3 | Denr |
| Rack1 | 0.00107932366539286 | 0.314353440119826 | 0.718 | 0.504 | 1 | Cluster 3 | Rack1 |
| Snrnp48 | 0.00109840281613477 | 0.251598248234994 | 0.279 | 0.124 | 1 | Cluster 3 | Snrnp48 |
| Ube3a | 0.00110896607425587 | 0.391884370602202 | 0.314 | 0.119 | 1 | Cluster 3 | Ube3a |
| Dpm3 | 0.00115217300846972 | 0.263014204789757 | 0.387 | 0.19 | 1 | Cluster 3 | Dpm3 |
| Anapc13 | 0.00116725682066775 | 0.295353071632206 | 0.289 | 0.112 | 1 | Cluster 3 | Anapc13 |
| Smim7 | 0.00117996020101955 | 0.301677191610241 | 0.341 | 0.174 | 1 | Cluster 3 | Smim7 |
| Bax1 | 0.00119087085744015 | 0.386997432713424 | 0.557 | 0.342 | 1 | Cluster 3 | Bax |

| | | | | | | | |
|---|---|---|---|---|---|---|---|
| Sugt1 | 0.00119371487682481 | 0.295289166818606 | 0.401 | 0.214 | 1 | Cluster 3 | Sugt1 |
| Tardbp | 0.00122134208215636 | 0.296527543208349 | 0.422 | 0.226 | 1 | Cluster 3 | Tardbp |
| Calm1 | 0.00122200559694217 | 0.284613744490135 | 0.976 | 0.903 | 1 | Cluster 3 | Calm1 |
| Gpd1l | 0.00125295851293255 | 0.337037636120857 | 0.206 | 0.051 | 1 | Cluster 3 | Gpd1l |
| Sf3b6 | 0.00125708912076577 | 0.298810166121012 | 0.676 | 0.463 | 1 | Cluster 3 | Sf3b6 |
| Ubxn11 | 0.00133139090295592 | 0.253747390482934 | 0.634 | 0.445 | 1 | Cluster 3 | Ubxn1 |
| Emc7 | 0.00133283059581687 | 0.270672878549786 | 0.321 | 0.172 | 1 | Cluster 3 | Emc7 |
| Ndufb11 | 0.00134175276447953 | 0.270965614375276 | 0.373 | 0.221 | 1 | Cluster 3 | Ndufb11 |
| Zfp91 | 0.00139608840746838 | 0.280847278042426 | 0.3 | 0.143 | 1 | Cluster 3 | Zfp91 |
| Dnajc8 | 0.00139827135661407 | 0.353889238367534 | 0.519 | 0.289 | 1 | Cluster 3 | Dnajc8 |
| Rer1 | 0.00140583077094768 | 0.294059691323522 | 0.659 | 0.44 | 1 | Cluster 3 | Rer1 |
| Ssbp1 | 0.00141200550002658 | 0.281457901244815 | 0.268 | 0.087 | 1 | Cluster 3 | Ssbp1 |
| Ndufaf81 | 0.00141595312681441 | 0.33496257302204 | 0.307 | 0.114 | 1 | Cluster 3 | Ndufaf8 |
| Rpl23a | 0.00144697273421916 | 0.342839911632723 | 0.819 | 0.616 | 1 | Cluster 3 | Rpl23a |
| Timm17a1 | 0.00155880665612078 | 0.312695582208857 | 0.286 | 0.096 | 1 | Cluster 3 | Timm17a |
| Tpr1 | 0.00161877614809806 | 0.358482202595699 | 0.491 | 0.284 | 1 | Cluster 3 | Tpr |
| Ssbp4 | 0.00176299891726169 | 0.303990804699649 | 0.286 | 0.117 | 1 | Cluster 3 | Ssbp4 |
| Prrc2c | 0.00179938851550785 | 0.262673342537537 | 0.732 | 0.513 | 1 | Cluster 3 | Prrc2c |
| Pfdn22 | 0.00180012658553098 | 0.353859507913437 | 0.679 | 0.373 | 1 | Cluster 3 | Pfdn2 |
| Romo11 | 0.00182367942982818 | 0.255046071857875 | 0.505 | 0.289 | 1 | Cluster 3 | Romo1 |
| Tmem14c | 0.0018302653200697 | 0.313709827113663 | 0.659 | 0.443 | 1 | Cluster 3 | Tmem14c |
| Tex261 | 0.00185573790991089 | 0.37510494559571 | 0.338 | 0.148 | 1 | Cluster 3 | Tex261 |
| Srsf71 | 0.0018693781318037 | 0.309337201307856 | 0.557 | 0.32 | 1 | Cluster 3 | Srsf7 |
| Atp5j | 0.00187509965012717 | 0.412259561791981 | 0.791 | 0.567 | 1 | Cluster 3 | Atp5j |
| Selenos1 | 0.00188457395603535 | 0.260787248415293 | 0.439 | 0.225 | 1 | Cluster 3 | Selenos |
| Tmco1 | 0.00190703838288574 | 0.419881433347087 | 0.429 | 0.18 | 1 | Cluster 3 | Tmco1 |
| Tmem128 | 0.00195278217669055 | 0.261562795977067 | 0.436 | 0.237 | 1 | Cluster 3 | Tmem128 |
| Cebpz1 | 0.00199062052215191 | 0.288690113518477 | 0.456 | 0.225 | 1 | Cluster 3 | Cebpz |
| Pole4 | 0.00199625419080708 | 0.295241728380839 | 0.328 | 0.149 | 1 | Cluster 3 | Pole4 |
| Gpr842 | 0.00204039788520047 | 0.260795540847428 | 0.613 | 0.451 | 1 | Cluster 3 | Gpr84 |
| Hnrnpr1 | 0.00209242858854289 | 0.299011486740269 | 0.328 | 0.157 | 1 | Cluster 3 | Hnrnpr |
| Ndufb3 | 0.00209512586767115 | 0.265256700646693 | 0.474 | 0.295 | 1 | Cluster 3 | Ndufb3 |
| Ctbp1 | 0.00209675222281689 | 0.261099456303014 | 0.31 | 0.141 | 1 | Cluster 3 | Ctbp1 |
| Hnrnpm | 0.00212037216040488 | 0.295532219016003 | 0.537 | 0.348 | 1 | Cluster 3 | Hnrnpm |
| Ppp6r1 | 0.00215273694302702 | 0.280324297585448 | 0.328 | 0.131 | 1 | Cluster 3 | Ppp6r1 |
| Ppp2r5c | 0.00224391364393839 | 0.309630060819123 | 0.46 | 0.242 | 1 | Cluster 3 | Ppp2r5c |
| Anapc11 | 0.00224832800412143 | 0.353260808511233 | 0.456 | 0.225 | 1 | Cluster 3 | Anapc11 |
| Dnajc2 | 0.00231476387176451 | 0.346568108045259 | 0.303 | 0.111 | 1 | Cluster 3 | Dnajc2 |
| Ndufa81 | 0.00231509566750515 | 0.305538575302094 | 0.474 | 0.254 | 1 | Cluster 3 | Ndufa8 |
| Eef1d1 | 0.00236785502108956 | 0.355616629886308 | 0.554 | 0.297 | 1 | Cluster 3 | Eef1d |
| Xpo1 | 0.00251031464979735 | 0.303609841980634 | 0.314 | 0.135 | 1 | Cluster 3 | Xpo1 |
| Ccdc59 | 0.00258489592289528 | 0.285801770166404 | 0.31 | 0.14 | 1 | Cluster 3 | Ccdc59 |
| Bcl2a1d1 | 0.00293048160874653 | 0.417574371344324 | 0.763 | 0.535 | 1 | Cluster 3 | Bcl2a1d |
| Ndufs2 | 0.002963376748181 | 0.268079416728305 | 0.345 | 0.153 | 1 | Cluster 3 | Ndufs2 |
| Fam110a | 0.00300478209407624 | 0.277479812784872 | 0.397 | 0.235 | 1 | Cluster 3 | Fam110a |
| Akirin1 | 0.00308469874010741 | 0.261966821999451 | 0.453 | 0.245 | 1 | Cluster 3 | Akirin1 |
| Tagln21 | 0.00324995340265449 | 0.292207365935989 | 0.568 | 0.418 | 1 | Cluster 3 | Tagln2 |
| Utp3 | 0.00335223739701329 | 0.27895125492083 | 0.289 | 0.125 | 1 | Cluster 3 | Utp3 |
| Sdhb | 0.00339455611956677 | 0.340665943066035 | 0.401 | 0.18 | 1 | Cluster 3 | Sdhb |
| Crlf3 | 0.00377167184280537 | 0.259333113389028 | 0.352 | 0.207 | 1 | Cluster 3 | Crlf3 |
| Ndufa12 | 0.00393656702453943 | 0.257518696769861 | 0.394 | 0.227 | 1 | Cluster 3 | Ndufa12 |
| Tcea1 | 0.0040145857224431 | 0.277667574561382 | 0.321 | 0.143 | 1 | Cluster 3 | Tcea1 |
| Jmjd1c | 0.00403681783121004 | 0.25227376880926 | 0.683 | 0.519 | 1 | Cluster 3 | Jmjd1c |
| Wbp11 | 0.00408348106729034 | 0.276204612835417 | 0.394 | 0.192 | 1 | Cluster 3 | Wbp11 |
| Rtf1 | 0.00410007764414789 | 0.257977612479143 | 0.31 | 0.148 | 1 | Cluster 3 | Rtf1 |
| Eif2s3x | 0.00414846006914221 | 0.251378562926726 | 0.244 | 0.089 | 1 | Cluster 3 | Eif2s3x |
| Ndufb71 | 0.00423350138846555 | 0.302980337072144 | 0.662 | 0.428 | 1 | Cluster 3 | Ndufb7 |
| Hsd17b10 | 0.00474294114267578 | 0.261236465177357 | 0.223 | 0.081 | 1 | Cluster 3 | Hsd17b10 |
| Mrps21 | 0.00487813984838226 | 0.2592853166535 | 0.561 | 0.348 | 1 | Cluster 3 | Mrps21 |
| mt-Nd4l | 0.00490442850442534 | 0.284968360930076 | 0.293 | 0.137 | 1 | Cluster 3 | mt-Nd4l |
| Zc3h15 | 0.00495798504491221 | 0.321208573008642 | 0.422 | 0.217 | 1 | Cluster 3 | Zc3h15 |

| | | | | | | | |
|---|---|---|---|---|---|---|---|
| Psma4 | 0.00496396934191419 | 0.290165727364724 | 0.373 | 0.184 | 1 | Cluster 3 | Psma4 |
| Rpl10a1 | 0.00532512229763271 | 0.305812090278283 | 0.599 | 0.377 | 1 | Cluster 3 | Rpl10a |
| Cct51 | 0.00533553935816453 | 0.362873793866986 | 0.401 | 0.18 | 1 | Cluster 3 | Cct5 |
| Nop581 | 0.00554736146372821 | 0.283755554819839 | 0.324 | 0.137 | 1 | Cluster 3 | Nop58 |
| Psmd4 | 0.00585472591422462 | 0.25338351706661 | 0.484 | 0.306 | 1 | Cluster 3 | Psmd4 |
| Hmgb1 | 0.00587924225402643 | 0.275498812270512 | 0.774 | 0.577 | 1 | Cluster 3 | Hmgb1 |
| Ubl5 | 0.00604364401671024 | 0.262671703243254 | 0.902 | 0.776 | 1 | Cluster 3 | Ubl5 |
| Ebi31 | 0.00610727274637774 | 0.294550026310385 | 0.408 | 0.25 | 1 | Cluster 3 | Ebi3 |
| Aurkaip1 | 0.0068665550286499 | 0.319914072190781 | 0.456 | 0.256 | 1 | Cluster 3 | Aurkaip1 |
| Dnajc19 | 0.00714919531888128 | 0.35173435669385 | 0.279 | 0.096 | 1 | Cluster 3 | Dnajc19 |
| Ndufa4 | 0.00753135477975837 | 0.317764253419387 | 0.418 | 0.217 | 1 | Cluster 3 | Ndufa4 |
| Sfr1 | 0.00756105163895812 | 0.299514310644346 | 0.331 | 0.184 | 1 | Cluster 3 | Sfr1 |
| Trappc2l | 0.00763582502014091 | 0.301264633722051 | 0.39 | 0.198 | 1 | Cluster 3 | Trappc2l |
| Clptm1l | 0.00781077929916009 | 0.256210346057837 | 0.439 | 0.224 | 1 | Cluster 3 | Clptm1l |
| Txn2 | 0.00785057223327363 | 0.360690588833928 | 0.369 | 0.146 | 1 | Cluster 3 | Txn2 |
| Smarca5 | 0.00787802740786795 | 0.265424241215252 | 0.491 | 0.295 | 1 | Cluster 3 | Smarca5 |
| Sh3bgrl3 | 0.00815388992026126 | 0.260573310858048 | 0.979 | 0.937 | 1 | Cluster 3 | Sh3bgrl3 |
| Taf6l | 0.00821272283160308 | 0.279549871682392 | 0.467 | 0.275 | 1 | Cluster 3 | Taf6l |
| Dnajc1 | 0.00843349897007298 | 0.25023877997367 | 0.251 | 0.116 | 1 | Cluster 3 | Dnajc1 |
| Ndufb10 | 0.00952171351207069 | 0.318365519653056 | 0.373 | 0.158 | 1 | Cluster 3 | Ndufb10 |
| Glg1 | 0.00981568228859404 | 0.260001756310411 | 0.314 | 0.159 | 1 | Cluster 3 | Glg1 |
| Ndufa7 | 0.0099895339971786 | 0.271003061819025 | 0.808 | 0.608 | 1 | Cluster 3 | Ndufa7 |
| Ccl63 | 5.29529899803744e-24 | 1.60891421460333 | 0.883 | 0.488 | 1.64434919786057e-19 | Cluster 4 | Ccl6 |
| Lcn22 | 5.6669789694551e-24 | 1.87886556105003 | 0.776 | 0.276 | 1.75976697938489e-19 | Cluster 4 | Lcn2 |
| Basp11 | 6.35596854354208e-22 | 1.02702557418534 | 0.98 | 0.756 | 1.97371891182612e-17 | Cluster 4 | Basp1 |
| Cxcl32 | 1.84671901046022e-21 | 2.27617766142827 | 0.857 | 0.388 | 5.73461654318213e-17 | Cluster 4 | Cxcl3 |
| Fpr12 | 8.76268178770361e-20 | 1.2371939135731 | 0.638 | 0.205 | 2.7210755755356e-15 | Cluster 4 | Fpr1 |
| Sgms23 | 1.02152171722266e-18 | 1.02733749594597 | 0.714 | 0.293 | 3.17213138849153e-14 | Cluster 4 | Sgms2 |
| S100a63 | 1.45577727283528e-18 | 0.990379260200271 | 0.954 | 0.792 | 4.52062516533539e-14 | Cluster 4 | S100a6 |
| Wfdc212 | 1.61588326248878e-18 | 1.69151695889713 | 0.724 | 0.28 | 5.01780229500642e-14 | Cluster 4 | Wfdc21 |
| Cd141 | 2.06671341684609e-18 | 0.787617119564038 | 0.974 | 0.836 | 6.41776517333215e-14 | Cluster 4 | Cd14 |
| Pkm2 | 3.1258326131939e-18 | 0.656232920949026 | 0.974 | 0.88 | 9.70664801375101e-14 | Cluster 4 | Pkm |
| Cd91 | 2.26567994040305e-17 | 0.587222280276073 | 0.995 | 0.923 | 7.03561591893359e-13 | Cluster 4 | Cd9 |
| Retnlg3 | 2.9740894510182e-17 | 2.30185386542256 | 0.612 | 0.167 | 9.23543997224683e-13 | Cluster 4 | Retnlg |
| Upp11 | 5.14157223589365e-17 | 1.21153480973795 | 0.709 | 0.235 | 1.59661242641206e-12 | Cluster 4 | Upp1 |
| Grina3 | 5.22903113131916e-17 | 0.790382464808845 | 0.959 | 0.754 | 1.62377103720854e-12 | Cluster 4 | Grina |
| Cd1772 | 1.06105011710868e-16 | 1.24074869212063 | 0.469 | 0.046 | 3.29487892865757e-12 | Cluster 4 | Cd177 |
| G0s21 | 2.17315263566196e-16 | 1.33221976460949 | 0.867 | 0.553 | 6.74829087952108e-12 | Cluster 4 | G0s2 |
| Ltb4r11 | 4.45066605944557e-16 | 0.887908627577509 | 0.704 | 0.279 | 1.38206533143963e-11 | Cluster 4 | Ltb4r1 |
| Srgn1 | 5.35441832615473e-16 | 0.502022456825752 | 1 | 0.998 | 1.66270752282083e-11 | Cluster 4 | Srgn |
| Hp3 | 1.4625419901528e-15 | 0.953128559592971 | 0.791 | 0.413 | 4.5416316420215e-11 | Cluster 4 | Hp |
| Smox3 | 1.81205070198518e-15 | 0.851988599818517 | 0.847 | 0.478 | 5.62696104487459e-11 | Cluster 4 | Smox |
| Slc16a32 | 1.08148769483059e-14 | 0.805595183621568 | 0.923 | 0.679 | 3.35834373875743e-10 | Cluster 4 | Slc16a3 |
| Itgam1 | 2.56754100854869e-14 | 0.672697468988659 | 0.918 | 0.661 | 7.97298509384624e-10 | Cluster 4 | Itgam |
| Rab442 | 7.02599320431276e-14 | 1.00521694965785 | 0.622 | 0.151 | 2.18178166973524e-09 | Cluster 4 | Rab44 |
| Tes | 1.91500632740108e-13 | 0.705957916129433 | 0.668 | 0.308 | 5.94666914847858e-09 | Cluster 4 | Tes |
| Pfkp | 2.93225947749521e-13 | 0.90174334172399 | 0.561 | 0.19 | 9.10554535546588e-09 | Cluster 4 | Pfkp |
| AC110211.13 | 2.93628307337954e-13 | 0.953902609532 | 0.673 | 0.222 | 9.11803982776549e-09 | Cluster 4 | AC110211.1 |
| Sell3 | 3.8068224912129e-13 | 0.849767557568685 | 0.638 | 0.256 | 1.18213258819634e-08 | Cluster 4 | Sell |
| Prdx52 | 5.35591100523689e-13 | 0.761132223920524 | 0.949 | 0.889 | 1.66317104445621e-08 | Cluster 4 | Prdx5 |
| Osm1 | 5.82668522675175e-13 | 1.55367715162299 | 0.653 | 0.293 | 1.80936056346322e-08 | Cluster 4 | Osm |
| Actg13 | 8.62971090366926e-13 | 0.602180146708077 | 0.995 | 0.985 | 2.67978412691641e-08 | Cluster 4 | Actg1 |
| Cd1641 | 1.00721016952768e-12 | 0.633629083281278 | 0.648 | 0.314 | 3.1276897394343e-08 | Cluster 4 | Cd164 |
| Plaur | 1.20044755153924e-12 | 0.932983112114823 | 0.954 | 0.719 | 3.72774978179479e-08 | Cluster 4 | Plaur |
| Pgk1 | 1.52163569170785e-12 | 1.07621322921411 | 0.719 | 0.364 | 4.7251353134604e-08 | Cluster 4 | Pgk1 |
| Slfn42 | 2.11316374047606e-12 | 1.0451828185274 | 0.796 | 0.398 | 6.5620073633003e-08 | Cluster 4 | Slfn4 |
| Glipr21 | 3.32001269576527e-12 | 0.795222385911763 | 0.684 | 0.267 | 1.03096354241599e-07 | Cluster 4 | Glipr2 |
| Mgst11 | 4.67488155218574e-12 | 1.0808980977829 | 0.434 | 0.085 | 1.45169096840024e-07 | Cluster 4 | Mgst1 |
| Lgals31 | 2.04256731338409e-11 | 0.7360054446247 | 0.913 | 0.757 | 6.3427842782516e-07 | Cluster 4 | Lgals3 |
| Plin2 | 2.1437065403656e-11 | 1.04673358796039 | 0.893 | 0.653 | 6.65685191979729e-07 | Cluster 4 | Plin2 |
| Thbs11 | 2.27178665348175e-11 | 1.32979212184444 | 0.837 | 0.595 | 7.05457909505689e-07 | Cluster 4 | Thbs1 |

| | | | | | | | |
|---|---|---|---|---|---|---|---|
| Anxa2 | 5.77300215715601e-11 | 0.653737251809621 | 0.898 | 0.767 | 1.79269035986166e-06 | Cluster 4 | Anxa2 |
| S100a92 | 7.11062017209111e-11 | 0.812966756750495 | 0.98 | 0.926 | 2.20806088203945e-06 | Cluster 4 | S100a9 |
| Crispld21 | 1.44041879247479e-10 | 0.725601295645366 | 0.347 | 0.065 | 4.47293247627197e-06 | Cluster 4 | Crispld2 |
| Fmnl12 | 1.95315234649079e-10 | 0.472529845034946 | 0.76 | 0.452 | 6.06512398155784e-06 | Cluster 4 | Fmnl1 |
| Flot12 | 2.06209489871587e-10 | 0.701633876520877 | 0.49 | 0.137 | 6.40342328898238e-06 | Cluster 4 | Flot1 |
| Gabbr1 | 2.16502699161306e-10 | 0.467298915866544 | 0.255 | 0.028 | 6.72305831705602e-06 | Cluster 4 | Gabbr1 |
| Rdh122 | 3.97083140449581e-10 | 0.721441174975913 | 0.469 | 0.126 | 1.23306227603808e-05 | Cluster 4 | Rdh12 |
| Stx11 | 6.70711176267469e-10 | 0.797129754117505 | 0.658 | 0.327 | 2.08275941566337e-05 | Cluster 4 | Stx11 |
| Anxa3 | 7.27762836584623e-10 | 0.768093814476965 | 0.602 | 0.204 | 2.25992193644623e-05 | Cluster 4 | Anxa3 |
| Adam82 | 7.45828638540291e-10 | 0.784737124221562 | 0.694 | 0.327 | 2.31602167125917e-05 | Cluster 4 | Adam8 |
| Lipg | 8.2712326680742e-10 | 0.734878780498353 | 0.235 | 0.026 | 2.56846588041708e-05 | Cluster 4 | Lipg |
| Tpi1 | 1.03047978718059e-09 | 0.780806686543066 | 0.556 | 0.325 | 3.19994888313189e-05 | Cluster 4 | Tpi1 |
| Gm145481 | 1.17361578640778e-09 | 0.532868014935531 | 0.454 | 0.144 | 3.64442910153209e-05 | Cluster 4 | Gm14548 |
| Lrg13 | 1.39019467508266e-09 | 0.790244693909702 | 0.602 | 0.297 | 4.31697152453417e-05 | Cluster 4 | Lrg1 |
| Fth11 | 1.55446133552778e-09 | 0.513612319872779 | 1 | 1 | 4.8270687852144e-05 | Cluster 4 | Fth1 |
| Gadd45a | 1.58352543472825e-09 | 0.857737461299652 | 0.607 | 0.269 | 4.91732153246162e-05 | Cluster 4 | Gadd45a |
| Cish | 1.70523015525008e-09 | 0.532766684768459 | 0.255 | 0.038 | 5.29525120109806e-05 | Cluster 4 | Cish |
| Tlr42 | 1.98807910554731e-09 | 0.804896735219176 | 0.617 | 0.197 | 6.17358204645608e-05 | Cluster 4 | Tlr4 |
| Diaph11 | 2.90625907798255e-09 | 0.580100899472136 | 0.796 | 0.524 | 9.02480631485922e-05 | Cluster 4 | Diaph1 |
| Cd300lf1 | 3.98359013224486e-09 | 0.547243844694136 | 0.929 | 0.756 | 0.0001237024243766 | Cluster 4 | Cd300lf |
| Dusp161 | 5.95543305621012e-09 | 0.582561840349709 | 0.684 | 0.392 | 0.000184934062694493 | Cluster 4 | Dusp16 |
| Gapdh1 | 7.83695389892214e-09 | 0.832306367878643 | 0.934 | 0.88 | 0.000243360929423229 | Cluster 4 | Gapdh |
| Bnip3 | 1.46738290189732e-08 | 0.91544935749878 | 0.454 | 0.186 | 0.000455666412526176 | Cluster 4 | Bnip3 |
| Bnip3l | 1.75757710610856e-08 | 0.521165819989462 | 0.663 | 0.459 | 0.000545780418759892 | Cluster 4 | Bnip3l |
| Trem11 | 2.16699483957418e-08 | 0.645441249187602 | 0.964 | 0.883 | 0.00067291690753297 | Cluster 4 | Trem1 |
| Fmnl2 | 2.9339752367375e-08 | 0.545189161512824 | 0.311 | 0.037 | 0.000911087330264095 | Cluster 4 | Fmnl2 |
| Gda2 | 3.90937706853759e-08 | 0.66050457059952 | 0.602 | 0.28 | 0.00121397886109298 | Cluster 4 | Gda |
| Tarm1 | 3.92984276958272e-08 | 0.662859101957991 | 0.429 | 0.137 | 0.00122033407523852 | Cluster 4 | Tarm1 |
| Ier3 | 4.30436097206798e-08 | 0.435300831957887 | 0.974 | 0.881 | 0.00133663321265627 | Cluster 4 | Ier3 |
| Slpi3 | 4.82171846930933e-08 | 0.563621645880333 | 0.811 | 0.554 | 0.00149728823627463 | Cluster 4 | Slpi |
| S100a82 | 5.51374977873411e-08 | 0.661618017752419 | 0.969 | 0.917 | 0.0017121847187903 | Cluster 4 | S100a8 |
| Slc7a112 | 5.71195172852579e-08 | 0.412108979204086 | 0.985 | 0.835 | 0.00177373237025911 | Cluster 4 | Slc7a11 |
| Fcgr2b | 7.38035139875364e-08 | 0.515936359135467 | 0.393 | 0.145 | 0.00229182051985497 | Cluster 4 | Fcgr2b |
| Eno1 | 7.45460352056624e-08 | 0.556637032884498 | 0.776 | 0.519 | 0.00231487803124143 | Cluster 4 | Eno1 |
| Gpat31 | 9.21652327844252e-08 | 0.547735857579006 | 0.434 | 0.143 | 0.00286200697365476 | Cluster 4 | Gpat3 |
| Spp1 | 1.00375048904384e-07 | 1.09732051658077 | 0.929 | 0.902 | 0.00311694639362783 | Cluster 4 | Spp1 |
| Lilra62 | 1.21185071905416e-07 | 0.458041053795725 | 0.49 | 0.256 | 0.00376316003787887 | Cluster 4 | Lilra6 |
| Tgm1 | 1.29575631574881e-07 | 0.508503240175457 | 0.209 | 0.014 | 0.00402371208729477 | Cluster 4 | Tgm1 |
| Mcemp13 | 1.33821890625258e-07 | 0.629673343808985 | 0.684 | 0.418 | 0.00415557116958613 | Cluster 4 | Mcemp1 |
| Adam19 | 1.55058605805025e-07 | 0.492595879335621 | 0.245 | 0.028 | 0.00481503488606345 | Cluster 4 | Adam19 |
| Slfn12 | 1.92807327211718e-07 | 0.429833417838394 | 0.76 | 0.524 | 0.00598724593190549 | Cluster 4 | Slfn1 |
| Xbp1 | 2.09111489551457e-07 | 0.711132849138765 | 0.602 | 0.309 | 0.0064935390850414 | Cluster 4 | Xbp1 |
| Rnasel | 2.39483249027902e-07 | 0.601649687140658 | 0.311 | 0.077 | 0.00743667333206344 | Cluster 4 | Rnasel |
| Rhoh | 3.15206716784155e-07 | 0.483022440318871 | 0.286 | 0.079 | 0.00978811417629838 | Cluster 4 | Rhoh |
| Zbp12 | 3.17351580792476e-07 | 0.601337570367929 | 0.332 | 0.085 | 0.00985471863834876 | Cluster 4 | Zbp1 |
| Fndc3a1 | 3.26890914739539e-07 | 0.701916286884272 | 0.485 | 0.172 | 0.0101509435754069 | Cluster 4 | Fndc3a |
| Hilpda1 | 3.93008482733958e-07 | 0.667563084904665 | 0.796 | 0.627 | 0.0122040924143376 | Cluster 4 | Hilpda |
| App | 3.96564161931017e-07 | 0.468509737044323 | 0.74 | 0.481 | 0.0123145069204439 | Cluster 4 | App |
| Cxcl23 | 4.12921745899329e-07 | 0.365419443739861 | 1 | 0.996 | 0.0128224589754119 | Cluster 4 | Cxcl2 |
| Gm54831 | 4.60009701035658e-07 | 1.59107144068432 | 0.398 | 0.218 | 0.0142846812462603 | Cluster 4 | Gm5483 |
| Clec4d3 | 5.2421012813473e-07 | 0.358467117185191 | 0.985 | 0.932 | 0.0162782971089678 | Cluster 4 | Clec4d |
| Ccl4 | 5.35864797930487e-07 | 0.945137435540489 | 0.776 | 0.598 | 0.0166402095701354 | Cluster 4 | Ccl4 |
| Fpr2 | 5.61417369069894e-07 | 0.503727671902422 | 0.648 | 0.393 | 0.0174336935617274 | Cluster 4 | Fpr2 |
| Sp1403 | 5.88638971978658e-07 | 0.536452793601177 | 0.587 | 0.303 | 0.0182790059968533 | Cluster 4 | Sp140 |
| Zfp36l1 | 6.22989804149297e-07 | 0.48063388809555 | 0.796 | 0.566 | 0.0193457023882481 | Cluster 4 | Zfp36l1 |
| Tmem189 | 6.90202691976299e-07 | 0.454564623189105 | 0.587 | 0.371 | 0.02143286419394 | Cluster 4 | Tmem189 |
| Tlr23 | 7.51874016064522e-07 | 0.473291220473985 | 0.648 | 0.376 | 0.0233479438208516 | Cluster 4 | Tlr2 |
| Pglyrp12 | 9.43143973261034e-07 | 0.402210965428645 | 0.663 | 0.493 | 0.0292874498016749 | Cluster 4 | Pglyrp1 |
| Slc15a33 | 9.79687666005623e-07 | 0.431628603370376 | 0.832 | 0.615 | 0.0304222410924726 | Cluster 4 | Slc15a3 |
| Ceacam10 | 1.08928734430255e-06 | 0.542424655619536 | 0.23 | 0.036 | 0.0338256399026271 | Cluster 4 | Ceacam10 |
| Mxd13 | 1.17036271242263e-06 | 0.410836138736479 | 0.969 | 0.861 | 0.0363432733088598 | Cluster 4 | Mxd1 |

| | | | | | | | |
|---|---|---|---|---|---|---|---|
| Ero1l | 1.24693546585974e-06 | 1.02469408808507 | 0.474 | 0.23 | 0.0387210870213425 | Cluster 4 | Ero1l |
| A530064D06Rik2 | 1.38261434969533e-06 | 0.464052660850261 | 0.291 | 0.083 | 0.0429343234010892 | Cluster 4 | A530064D06Rik |
| Ccr12 | 1.46158705071231e-06 | 0.342691276284135 | 0.985 | 0.897 | 0.0453866626857693 | Cluster 4 | Ccr1 |
| Wfdc173 | 1.50422955926903e-06 | 1.34575732494877 | 0.704 | 0.549 | 0.0467108405039812 | Cluster 4 | Wfdc17 |
| Arg2 | 1.75153727374628e-06 | 0.419457336661452 | 0.77 | 0.572 | 0.0543904869616434 | Cluster 4 | Arg2 |
| Egr1 | 1.82264842129708e-06 | 0.456899149205534 | 0.929 | 0.805 | 0.0565987014265381 | Cluster 4 | Egr1 |
| Acvrl1 | 3.25981004680397e-06 | 0.530485620455358 | 0.184 | 0.021 | 0.101226881383404 | Cluster 4 | Acvrl1 |
| Slc11a1 | 3.38111770585037e-06 | 0.550997191959005 | 0.663 | 0.385 | 0.104993848119771 | Cluster 4 | Slc11a1 |
| Il1rap | 3.71768515967225e-06 | 0.498954562593646 | 0.653 | 0.388 | 0.115445277263302 | Cluster 4 | Il1rap |
| Rtp41 | 3.95904868670496e-06 | 0.36305703160611 | 0.449 | 0.209 | 0.122940338868249 | Cluster 4 | Rtp4 |
| Snx181 | 4.69931518706678e-06 | 0.473846769226131 | 0.745 | 0.509 | 0.145927834503985 | Cluster 4 | Snx18 |
| Mmp19 | 5.386233347186e-06 | 0.56079056595011 | 0.235 | 0.074 | 0.167258704130167 | Cluster 4 | Mmp19 |
| Egln1 | 5.61048345878379e-06 | 0.4790712588062 | 0.25 | 0.12 | 0.174222342845613 | Cluster 4 | Egln1 |
| Pnrc12 | 6.00241963155864e-06 | 0.403927336487389 | 0.964 | 0.891 | 0.18639313681879 | Cluster 4 | Pnrc1 |
| Eif4ebp1 | 6.50538742600326e-06 | 0.644431463018585 | 0.541 | 0.309 | 0.202011795739679 | Cluster 4 | Eif4ebp1 |
| Emilin21 | 7.89271632682937e-06 | 0.489216164806493 | 0.883 | 0.698 | 0.245092520097033 | Cluster 4 | Emilin2 |
| Acta21 | 8.28943245882668e-06 | 0.495061985430177 | 0.265 | 0.093 | 0.257411746143945 | Cluster 4 | Acta2 |
| Dmxl21 | 8.75929307579517e-06 | 0.572704861278063 | 0.454 | 0.227 | 0.272002327882667 | Cluster 4 | Dmxl2 |
| Fam162a | 8.88210113893264e-06 | 0.738829992050895 | 0.378 | 0.16 | 0.275815886667275 | Cluster 4 | Fam162a |
| Esd | 1.04766717596083e-05 | 0.497611542522725 | 0.612 | 0.366 | 0.325332088151116 | Cluster 4 | Esd |
| Pygl | 1.05738115468295e-05 | 0.34250080076983 | 0.628 | 0.439 | 0.328348569963696 | Cluster 4 | Pygl |
| Crem | 1.06636845938766e-05 | 0.598900195262865 | 0.469 | 0.229 | 0.33113939769365 | Cluster 4 | Crem |
| Lpp | 1.07056322489842e-05 | 0.282149544953651 | 0.383 | 0.203 | 0.332441998227706 | Cluster 4 | Lpp |
| Il13ra1 | 1.12305597242294e-05 | 0.408810102438725 | 0.571 | 0.351 | 0.348742571116496 | Cluster 4 | Il13ra1 |
| Ripor22 | 1.31327320679997e-05 | 0.33030131934843 | 0.352 | 0.17 | 0.407810728907595 | Cluster 4 | Ripor2 |
| Nedd92 | 1.41620539707971e-05 | 0.471918492206844 | 0.408 | 0.142 | 0.439774261955162 | Cluster 4 | Nedd9 |
| Ifitm62 | 1.50318619131695e-05 | 0.488920713915141 | 0.332 | 0.144 | 0.466784407989652 | Cluster 4 | Ifitm6 |
| Samd9l1 | 1.59511105677578e-05 | 0.325036093334903 | 0.459 | 0.263 | 0.495329836460583 | Cluster 4 | Samd9l |
| Tnfaip6 | 1.65202145898805e-05 | 0.368678756425625 | 0.194 | 0.05 | 0.513002223659559 | Cluster 4 | Tnfaip6 |
| Isg201 | 1.74090749844847e-05 | 0.59682728971679 | 0.5 | 0.226 | 0.540604005493202 | Cluster 4 | Isg20 |
| Capg1 | 1.76033360472069e-05 | 0.396521847483643 | 0.714 | 0.529 | 0.546636394273914 | Cluster 4 | Capg |
| Selplg3 | 1.83774454053885e-05 | 0.400374799152469 | 0.842 | 0.691 | 0.570674812173529 | Cluster 4 | Selplg |
| Inhba | 1.86897307559532e-05 | 0.441610211294092 | 0.27 | 0.114 | 0.580372209164615 | Cluster 4 | Inhba |
| Impa2 | 1.89372053207443e-05 | 0.58256010109572 | 0.23 | 0.041 | 0.588057036825074 | Cluster 4 | Impa2 |
| Isg151 | 1.91850563319872e-05 | 0.542703429593964 | 0.582 | 0.366 | 0.595753554277199 | Cluster 4 | Isg15 |
| Samsn11 | 1.99240955411144e-05 | 0.310358885714781 | 0.949 | 0.816 | 0.618702938838225 | Cluster 4 | Samsn1 |
| Btg11 | 2.11622008763473e-05 | 0.326500516233327 | 0.99 | 0.979 | 0.657149823813213 | Cluster 4 | Btg1 |
| Gm14005 | 2.30990114561445e-05 | 0.377472694773327 | 0.464 | 0.27 | 0.717293602747655 | Cluster 4 | Gm14005 |
| Chil31 | 2.34011206633576e-05 | 1.30882778114944 | 0.546 | 0.332 | 0.726674999959243 | Cluster 4 | Chil3 |
| Ly6c22 | 2.76490734282085e-05 | 0.645999584207924 | 0.281 | 0.109 | 0.858586677166159 | Cluster 4 | Ly6c2 |
| C31 | 2.78217190611056e-05 | 0.405153030321904 | 0.694 | 0.501 | 0.863947842004512 | Cluster 4 | C3 |
| Rhoq | 3.01761544892835e-05 | 0.416088214725483 | 0.235 | 0.059 | 0.937060125355721 | Cluster 4 | Rhoq |
| Lilr4b1 | 3.23690590229384e-05 | 0.41503578626097 | 0.929 | 0.813 | 1 | Cluster 4 | Lilr4b |
| Dach1 | 3.38085723046232e-05 | 0.292023511748156 | 0.163 | 0.017 | 1 | Cluster 4 | Dach1 |
| Mmp81 | 3.8940882419992e-05 | 0.677607168133381 | 0.296 | 0.084 | 1 | Cluster 4 | Mmp8 |
| Ifi2042 | 4.19959136576112e-05 | 0.511806006675622 | 0.429 | 0.222 | 1 | Cluster 4 | Ifi204 |
| Rgcc1 | 4.44820365669966e-05 | 0.517755838989842 | 0.694 | 0.505 | 1 | Cluster 4 | Rgcc |
| Carhsp1 | 4.55529915032924e-05 | 0.423871130985081 | 0.418 | 0.211 | 1 | Cluster 4 | Carhsp1 |
| Egln3 | 4.60895316539297e-05 | 0.48759650307031 | 0.311 | 0.103 | 1 | Cluster 4 | Egln3 |
| Jaml1 | 5.39447658224731e-05 | 0.515206663522879 | 0.383 | 0.154 | 1 | Cluster 4 | Jaml |
| Phf21a | 5.5531754441314e-05 | 0.342905558096609 | 0.347 | 0.152 | 1 | Cluster 4 | Phf21a |
| Timp21 | 6.73790087101472e-05 | 0.304544095323222 | 0.383 | 0.206 | 1 | Cluster 4 | Timp2 |
| Snai1 | 6.90116395675372e-05 | 0.430671289301733 | 0.189 | 0.022 | 1 | Cluster 4 | Snai1 |
| Tnfrsf26 | 9.02156208068877e-05 | 0.399525291462688 | 0.296 | 0.145 | 1 | Cluster 4 | Tnfrsf26 |
| Fbxl51 | 9.18238861690829e-05 | 0.515047465529868 | 0.546 | 0.307 | 1 | Cluster 4 | Fbxl5 |
| Glrx1 | 9.24938154079241e-05 | 0.698256263868471 | 0.577 | 0.295 | 1 | Cluster 4 | Glrx |
| Rap2a | 9.74475087742835e-05 | 0.371542752523485 | 0.189 | 0.048 | 1 | Cluster 4 | Rap2a |
| Oasl21 | 0.000106985754504259 | 0.380426600177891 | 0.597 | 0.357 | 1 | Cluster 4 | Oasl2 |
| Rps6ka2 | 0.000110366513763574 | 0.332284875754783 | 0.199 | 0.055 | 1 | Cluster 4 | Rps6ka2 |
| Fosl1 | 0.000115167409146954 | 0.304635867969607 | 0.194 | 0.06 | 1 | Cluster 4 | Fosl1 |
| Gpr351 | 0.000116626763833126 | 0.339116455335153 | 0.612 | 0.413 | 1 | Cluster 4 | Gpr35 |

| | | | | | | | | |
|---|---|---|---|---|---|---|---|---|
| Ccl32 | 0.000117828297380103 | 0.287002711043538 | 0.883 | 0.832 | 1 | | Cluster 4 | Ccl3 |
| Qsox1 | 0.000119902918978571 | 0.312930408032692 | 0.214 | 0.084 | 1 | | Cluster 4 | Qsox1 |
| Cdk6 | 0.000131350891851061 | 0.545912768603214 | 0.24 | 0.093 | 1 | | Cluster 4 | Cdk6 |
| Homer1 | 0.000131904340251041 | 0.374925342640914 | 0.418 | 0.226 | 1 | | Cluster 4 | Homer1 |
| Atp11b | 0.000133659711002319 | 0.287306382292953 | 0.413 | 0.227 | 1 | | Cluster 4 | Atp11b |
| Enah | 0.000135064984943156 | 0.271662787773202 | 0.138 | 0.024 | 1 | | Cluster 4 | Enah |
| Gpi12 | 0.000146810575331095 | 0.629777320987477 | 0.628 | 0.473 | 1 | | Cluster 4 | Gpi1 |
| Cd53 | 0.000174103218603081 | 0.293731297724257 | 0.969 | 0.915 | 1 | | Cluster 4 | Cd53 |
| Prr131 | 0.000181010780442587 | 0.489028677485772 | 0.648 | 0.46 | 1 | | Cluster 4 | Prr13 |
| Mafg | 0.000193677529685039 | 0.268642005894883 | 0.5 | 0.347 | 1 | | Cluster 4 | Mafg |
| Trpv2 | 0.000220020325044473 | 0.401196728576889 | 0.413 | 0.184 | 1 | | Cluster 4 | Trpv2 |
| Slc27a41 | 0.000221273135743849 | 0.431762888779468 | 0.327 | 0.105 | 1 | | Cluster 4 | Slc27a4 |
| Aldoa | 0.0002245297821646 | 0.447637976451455 | 0.867 | 0.787 | 1 | | Cluster 4 | Aldoa |
| Pira21 | 0.000228715283519728 | 0.277782750695248 | 0.464 | 0.299 | 1 | | Cluster 4 | Pira2 |
| Phlda11 | 0.000229019280863176 | 0.51252814114825 | 0.679 | 0.53 | 1 | | Cluster 4 | Phlda1 |
| Il1f91 | 0.000240984791220445 | 0.81137142916885 | 0.643 | 0.365 | 1 | | Cluster 4 | Il1f9 |
| Cd47 | 0.000243416171321381 | 0.580248443410661 | 0.781 | 0.629 | 1 | | Cluster 4 | Cd47 |
| Chil13 | 0.000263481880995316 | 0.353759828896943 | 0.388 | 0.194 | 1 | | Cluster 4 | Chil1 |
| F10 | 0.000279729770181087 | 0.38427135948737 | 0.327 | 0.168 | 1 | | Cluster 4 | F10 |
| Tec | 0.000282675397241934 | 0.327022707922567 | 0.276 | 0.127 | 1 | | Cluster 4 | Tec |
| Bcl2l1 | 0.000284670302161661 | 0.323508018354023 | 0.566 | 0.358 | 1 | | Cluster 4 | Bcl2l1 |
| Slc2a31 | 0.000312773130443945 | 0.3792394197555 | 0.24 | 0.117 | 1 | | Cluster 4 | Slc2a3 |
| Ywhaz | 0.000397632635440204 | 0.328056615602799 | 0.801 | 0.625 | 1 | | Cluster 4 | Ywhaz |
| Gm19951 | 0.000413592227638439 | 0.585860056428589 | 0.327 | 0.17 | 1 | | Cluster 4 | Gm19951 |
| Rnf1492 | 0.000414472125471027 | 0.37168490639062 | 0.939 | 0.869 | 1 | | Cluster 4 | Rnf149 |
| Padi4 | 0.000562071457492247 | 0.257853339973903 | 0.184 | 0.065 | 1 | | Cluster 4 | Padi4 |
| Dgat21 | 0.000602294732956705 | 0.624183757371575 | 0.367 | 0.122 | 1 | | Cluster 4 | Dgat2 |
| Cd33 | 0.00060554109422765 | 0.396098608146022 | 0.827 | 0.662 | 1 | | Cluster 4 | Cd33 |
| Fkbp1a | 0.000609641532363941 | 0.426813117309189 | 0.306 | 0.137 | 1 | | Cluster 4 | Fkbp1a |
| Pgm2 | 0.000690390853667533 | 0.255114486390127 | 0.153 | 0.035 | 1 | | Cluster 4 | Pgm2 |
| Cxcr22 | 0.000692295880247264 | 0.3515116815489 | 0.765 | 0.632 | 1 | | Cluster 4 | Cxcr2 |
| Csf2rb | 0.000716667564956158 | 0.328808527033156 | 0.821 | 0.624 | 1 | | Cluster 4 | Csf2rb |
| Ffar2 | 0.000717923654542841 | 0.304353482920148 | 0.168 | 0.042 | 1 | | Cluster 4 | Ffar2 |
| Klhdc4 | 0.000761743962580267 | 0.331923327676271 | 0.291 | 0.149 | 1 | | Cluster 4 | Klhdc4 |
| Entpd12 | 0.000774376854067487 | 0.399271596919034 | 0.735 | 0.517 | 1 | | Cluster 4 | Entpd1 |
| Asnsd1 | 0.000778096457981952 | 0.257767716596456 | 0.291 | 0.171 | 1 | | Cluster 4 | Asnsd1 |
| Chd7 | 0.000833587698516079 | 0.370240476041592 | 0.724 | 0.578 | 1 | | Cluster 4 | Chd7 |
| Npepps | 0.000898230632200046 | 0.500394281363642 | 0.449 | 0.234 | 1 | | Cluster 4 | Npepps |
| Alas1 | 0.000937674051511349 | 0.357198246417112 | 0.582 | 0.387 | 1 | | Cluster 4 | Alas1 |
| Samhd12 | 0.00114131050150504 | 0.267505607829866 | 0.74 | 0.599 | 1 | | Cluster 4 | Samhd1 |
| Hif1a2 | 0.00114931839530468 | 0.402537754740657 | 0.592 | 0.362 | 1 | | Cluster 4 | Hif1a |
| Uck2 | 0.00116168038019859 | 0.348527443023138 | 0.204 | 0.107 | 1 | | Cluster 4 | Uck2 |
| Ly6g | 0.0012493917452647 | 0.299708392737416 | 0.148 | 0.025 | 1 | | Cluster 4 | Ly6g |
| Gys1 | 0.00127519519882892 | 0.331499778378971 | 0.163 | 0.04 | 1 | | Cluster 4 | Gys1 |
| Rab20 | 0.0013012282429631 | 0.31114395339507 | 0.867 | 0.722 | 1 | | Cluster 4 | Rab20 |
| Ppp1r3b | 0.00144595920715016 | 0.351588266170377 | 0.408 | 0.206 | 1 | | Cluster 4 | Ppp1r3b |
| Ifitm32 | 0.00146119158114382 | 0.625612974999041 | 0.796 | 0.679 | 1 | | Cluster 4 | Ifitm3 |
| Nt5e | 0.00154999786727481 | 0.690588552924191 | 0.429 | 0.218 | 1 | | Cluster 4 | Nt5e |
| Cpd | 0.00155383852907255 | 0.335199820869231 | 0.286 | 0.17 | 1 | | Cluster 4 | Cpd |
| Iqsec1 | 0.00173797909467485 | 0.30007210130341 | 0.495 | 0.323 | 1 | | Cluster 4 | Iqsec1 |
| Plk3 | 0.00174728796915258 | 0.56703777969288 | 0.668 | 0.447 | 1 | | Cluster 4 | Plk3 |
| Trim30b2 | 0.00185775006356373 | 0.337763888230191 | 0.571 | 0.336 | 1 | | Cluster 4 | Trim30b |
| Manf | 0.0019380672376086 | 0.491826004888787 | 0.281 | 0.171 | 1 | | Cluster 4 | Manf |
| Ell | 0.00198128474140116 | 0.357185448017401 | 0.255 | 0.083 | 1 | | Cluster 4 | Ell |
| Rbpj | 0.00199637123636364 | 0.396606644553765 | 0.679 | 0.492 | 1 | | Cluster 4 | Rbpj |
| Cyth1 | 0.00205630780795335 | 0.272187628406532 | 0.347 | 0.251 | 1 | | Cluster 4 | Cyth1 |
| Il1r2 | 0.00211004880794447 | 0.344100241163466 | 0.974 | 0.921 | 1 | | Cluster 4 | Il1r2 |
| Asprv13 | 0.00237928926083618 | 0.377286938376475 | 0.327 | 0.214 | 1 | | Cluster 4 | Asprv1 |
| Cast | 0.00259254639276316 | 0.320038142514252 | 0.459 | 0.324 | 1 | | Cluster 4 | Cast |
| Fas | 0.00281430733546809 | 0.394529997651146 | 0.464 | 0.286 | 1 | | Cluster 4 | Fas |
| St3gal6 | 0.0028784917446993 | 0.384272533697236 | 0.495 | 0.28 | 1 | | Cluster 4 | St3gal6 |

| | | | | | | | |
|---|---|---|---|---|---|---|---|
| Adora2b | 0.00295538053881754 | 0.407191523211793 | 0.235 | 0.077 | 1 | Cluster 4 | Adora2b |
| Nsd3 | 0.00295726457735691 | 0.265951831384363 | 0.566 | 0.396 | 1 | Cluster 4 | Nsd3 |
| P4ha1 | 0.00305239970503301 | 0.499237709677097 | 0.306 | 0.155 | 1 | Cluster 4 | P4ha1 |
| Gm20234 | 0.00311239748614287 | 0.497606078466671 | 0.158 | 0.065 | 1 | Cluster 4 | Gm20234 |
| P4hb | 0.00312447584529368 | 0.55587740247012 | 0.546 | 0.355 | 1 | Cluster 4 | P4hb |
| Neat11 | 0.00314589655241849 | 0.259097637444739 | 0.969 | 0.956 | 1 | Cluster 4 | Neat1 |
| Ptpn1 | 0.00314915499994239 | 0.405037474072541 | 0.602 | 0.409 | 1 | Cluster 4 | Ptpn1 |
| Fcgr3 | 0.00323145077005792 | 0.519011344101314 | 0.745 | 0.6 | 1 | Cluster 4 | Fcgr3 |
| Treml21 | 0.00346589694617041 | 0.263452306677726 | 0.27 | 0.131 | 1 | Cluster 4 | Treml2 |
| Id1 | 0.00350821593422707 | 0.449882004865734 | 0.235 | 0.16 | 1 | Cluster 4 | Id1 |
| Tnfrsf231 | 0.00369347430838538 | 0.293331191884859 | 0.612 | 0.428 | 1 | Cluster 4 | Tnfrsf23 |
| Mbd2 | 0.00395827364795542 | 0.387270321711476 | 0.658 | 0.431 | 1 | Cluster 4 | Mbd2 |
| Lamp21 | 0.00399483148755375 | 0.30543734932384 | 0.74 | 0.617 | 1 | Cluster 4 | Lamp2 |
| Stfa2l11 | 0.00490263974760967 | 0.831219321281151 | 0.332 | 0.237 | 1 | Cluster 4 | Stfa2l1 |
| Mif1 | 0.00524576406148079 | 0.649685158788653 | 0.714 | 0.62 | 1 | Cluster 4 | Mif |
| Tiparp | 0.00552851431004829 | 0.354580025457075 | 0.541 | 0.4 | 1 | Cluster 4 | Tiparp |
| Osbpl9 | 0.00580861092992597 | 0.341259693742882 | 0.536 | 0.419 | 1 | Cluster 4 | Osbpl9 |
| Antxr2 | 0.00676224310098542 | 0.308076897353603 | 0.663 | 0.538 | 1 | Cluster 4 | Antxr2 |
| Noct | 0.00676832319449118 | 0.388737709033906 | 0.546 | 0.356 | 1 | Cluster 4 | Noct |
| Xaf1 | 0.00720679153903199 | 0.260625990188685 | 0.173 | 0.076 | 1 | Cluster 4 | Xaf1 |
| Nfil3 | 0.00728677759409955 | 0.313862453901212 | 0.342 | 0.205 | 1 | Cluster 4 | Nfil3 |
| Btg2 | 0.00729887860838632 | 0.373875712927693 | 0.98 | 0.959 | 1 | Cluster 4 | Btg2 |
| Atxn1 | 0.00731270411800088 | 0.304942821478521 | 0.224 | 0.086 | 1 | Cluster 4 | Atxn1 |
| Il1a1 | 0.00810966177253062 | 0.307040476057757 | 0.638 | 0.504 | 1 | Cluster 4 | Il1a |
| Dstn | 0.00835280655732986 | 0.297338324029291 | 0.398 | 0.314 | 1 | Cluster 4 | Dstn |
| Map2k1 | 0.00954572386439037 | 0.251618916272548 | 0.301 | 0.179 | 1 | Cluster 4 | Map2k1 |
| Pts | 0.00974764742753209 | 0.365774488894539 | 0.495 | 0.352 | 1 | Cluster 4 | Pts |
| Slfn5 | 1.79359735356676e-36 | 2.0111329030687 | 0.978 | 0.232 | 5.56965786203087e-32 | Cluster 5 | Slfn5 |
| Ifit11 | 1.63447449187149e-35 | 2.83323396998262 | 0.856 | 0.082 | 5.07553363960852e-31 | Cluster 5 | Ifit1 |
| Isg152 | 1.85591278423216e-35 | 2.27919470782522 | 0.989 | 0.356 | 5.76316596887612e-31 | Cluster 5 | Isg15 |
| Rsad21 | 2.19244214524479e-35 | 3.21502964112664 | 0.9 | 0.138 | 6.80819059362866e-31 | Cluster 5 | Rsad2 |
| Oasl22 | 3.01856500176494e-33 | 1.74032361779988 | 0.978 | 0.351 | 9.37354989998068e-29 | Cluster 5 | Oasl2 |
| Irf71 | 4.53185713629577e-32 | 1.8728860685054 | 0.911 | 0.208 | 1.40727759653393e-27 | Cluster 5 | Irf7 |
| Parp14 | 7.61289429154451e-32 | 1.68943120332197 | 0.811 | 0.145 | 2.36403206435332e-27 | Cluster 5 | Parp14 |
| Ifit3b1 | 2.15947016335147e-31 | 2.23743687063487 | 0.733 | 0.03 | 6.70580269825531e-27 | Cluster 5 | Ifit3b |
| Ifit32 | 7.64476435792852e-31 | 2.57788943604461 | 0.744 | 0.049 | 2.37392867606754e-26 | Cluster 5 | Ifit3 |
| Isg202 | 1.1034086431832e-29 | 1.83466289342022 | 0.9 | 0.221 | 3.42641485967678e-25 | Cluster 5 | Isg20 |
| Oasl11 | 4.23089950698707e-28 | 1.8740602088119 | 0.733 | 0.078 | 1.3138212239047e-23 | Cluster 5 | Oasl1 |
| Usp181 | 8.61097910505086e-28 | 1.68184791607442 | 0.733 | 0.052 | 2.67396734149144e-23 | Cluster 5 | Usp18 |
| Slfn43 | 2.92857397943559e-27 | 1.5729754237496 | 0.967 | 0.418 | 9.09410077834133e-23 | Cluster 5 | Slfn4 |
| Herc6 | 2.19263859628362e-26 | 1.42952893178255 | 0.633 | 0.04 | 6.80880063303953e-22 | Cluster 5 | Herc6 |
| Samd9l2 | 5.46233301934624e-26 | 1.27376917470771 | 0.844 | 0.252 | 1.69621827249759e-21 | Cluster 5 | Samd9l |
| Ifi2043 | 1.48993535817956e-25 | 1.91800427803386 | 0.8 | 0.213 | 4.62669626775498e-21 | Cluster 5 | Ifi204 |
| Irgm11 | 9.2793902512437e-24 | 1.25136557340849 | 0.644 | 0.062 | 2.8815290547187e-19 | Cluster 5 | Irgm1 |
| Ifitm33 | 1.39412276718345e-23 | 1.64182874466306 | 0.967 | 0.677 | 4.32916942893477e-19 | Cluster 5 | Ifitm3 |
| Rtp42 | 1.64494859449374e-23 | 1.56254319178379 | 0.878 | 0.199 | 5.10805887048143e-19 | Cluster 5 | Rtp4 |
| Gm138221 | 2.63800832020697e-23 | 1.55609930921289 | 0.733 | 0.129 | 8.19180723673871e-19 | Cluster 5 | Gm13822 |
| Trim12a1 | 3.22071484362873e-23 | 1.31219707779929 | 0.811 | 0.218 | 1.00012858039203e-18 | Cluster 5 | Trim12a |
| Oas31 | 1.7474706559467e-22 | 1.379167139003 | 0.8 | 0.141 | 5.42642062791129e-18 | Cluster 5 | Oas3 |
| Slfn8 | 2.19235131494191e-22 | 1.31790033010278 | 0.756 | 0.123 | 6.8079085382891e-18 | Cluster 5 | Slfn8 |
| B2m | 6.83847988825488e-22 | 0.822506278694474 | 1 | 0.95 | 2.12355315969979e-17 | Cluster 5 | B2m |
| Plac8 | 9.94499750575741e-22 | 1.73882149438456 | 0.789 | 0.259 | 3.08822007546285e-17 | Cluster 5 | Plac8 |
| Sp1001 | 2.24379873792744e-21 | 1.28045592379436 | 0.844 | 0.298 | 6.96766822088608e-17 | Cluster 5 | Sp100 |
| Igtp | 4.62282071436337e-21 | 1.10339695347657 | 0.656 | 0.07 | 1.43552451643126e-16 | Cluster 5 | Igtp |
| Ddx581 | 1.22511036361033e-20 | 1.19962325739428 | 0.7 | 0.13 | 3.80433521211915e-16 | Cluster 5 | Ddx58 |
| Trim30a1 | 3.44772513503553e-20 | 1.37035721244856 | 0.889 | 0.398 | 1.07062208618258e-15 | Cluster 5 | Trim30a |
| Ifi47 | 4.48155471496405e-20 | 1.56203649106796 | 0.522 | 0.033 | 1.39165718563779e-15 | Cluster 5 | Ifi47 |
| Trafd11 | 1.41498934811017e-19 | 1.22638770269773 | 0.678 | 0.13 | 4.3939664226865e-15 | Cluster 5 | Trafd1 |
| Dtx3l | 1.62745868636307e-19 | 1.0839731981284 | 0.6 | 0.084 | 5.05374745876324e-15 | Cluster 5 | Dtx3l |
| Ddx601 | 4.2849037314609e-19 | 1.22680755791386 | 0.578 | 0.078 | 1.33059115573055e-14 | Cluster 5 | Ddx60 |
| Cmpk2 | 9.78710854366013e-19 | 1.82774057602118 | 0.511 | 0.028 | 3.03919081606278e-14 | Cluster 5 | Cmpk2 |

| | | | | | | | |
|---|---|---|---|---|---|---|---|
| H2-T231 | 1.28459542906557e-18 | 0.965401017522477 | 0.844 | 0.337 | 3.98905418587733e-14 | Cluster 5 | H2-T23 |
| Bst21 | 3.88587324834705e-18 | 1.28100336208947 | 0.711 | 0.216 | 1.20668021980921e-13 | Cluster 5 | Bst2 |
| H2-T22 | 7.91917037239213e-18 | 0.876306552717905 | 0.544 | 0.082 | 2.45913997573893e-13 | Cluster 5 | H2-T22 |
| Epsti11 | 1.53931734156118e-17 | 1.20521893040356 | 0.733 | 0.246 | 4.78004214074992e-13 | Cluster 5 | Epsti1 |
| Gbp3 | 3.72728233243236e-17 | 0.964404237875281 | 0.422 | 0.008 | 1.15743298269022e-12 | Cluster 5 | Gbp3 |
| Trim12c1 | 5.55067307810404e-17 | 1.08525587956223 | 0.778 | 0.225 | 1.72365051094365e-12 | Cluster 5 | Trim12c |
| Ifi209 | 5.78725314160025e-17 | 1.09250841387819 | 0.433 | 0.044 | 1.79711571806113e-12 | Cluster 5 | Ifi209 |
| Rnf213 | 8.58393814514242e-17 | 1.06894564445328 | 0.678 | 0.17 | 2.66557031221108e-12 | Cluster 5 | Rnf213 |
| Ifi207 | 9.69519066526521e-17 | 1.20692897824059 | 0.711 | 0.234 | 3.01064755728481e-12 | Cluster 5 | Ifi207 |
| Gbp2 | 1.08443718290887e-16 | 1.79019892073175 | 0.489 | 0.049 | 3.36750278408691e-12 | Cluster 5 | Gbp2 |
| Gbp7 | 1.63207789762807e-16 | 1.08551283096546 | 0.556 | 0.097 | 5.06809149550446e-12 | Cluster 5 | Gbp7 |
| Ly6e | 3.31629934273347e-16 | 0.977378338719456 | 0.933 | 0.731 | 1.02981043489902e-11 | Cluster 5 | Ly6e |
| Ifit2 | 3.8431937075534e-16 | 1.5865892798007 | 0.4 | 0.011 | 1.19342694200656e-11 | Cluster 5 | Ifit2 |
| H2-K1 | 6.47493302178393e-16 | 0.585149937096015 | 1 | 0.895 | 2.01066095125457e-11 | Cluster 5 | H2-K1 |
| Stat1 | 2.74098524142761e-15 | 1.27102697063533 | 0.7 | 0.179 | 8.51158147020515e-11 | Cluster 5 | Stat1 |
| Ifi211 | 3.81846416585581e-15 | 1.11497456039161 | 0.378 | 0.037 | 1.18574767742321e-10 | Cluster 5 | Ifi211 |
| Irf9 | 6.301351850465e-15 | 0.687039900025881 | 0.644 | 0.145 | 1.9567587901249e-10 | Cluster 5 | Irf9 |
| Slfn13 | 2.09347366903762e-14 | 1.02439018121312 | 0.978 | 0.528 | 6.50086378446253e-10 | Cluster 5 | Slfn1 |
| 9930111J21Rik2 | 3.77744048480832e-14 | 1.12850383464076 | 0.556 | 0.15 | 1.17300859374753e-09 | Cluster 5 | 9930111J21Rik2 |
| Tap1 | 7.12329318061827e-14 | 0.978695852529447 | 0.6 | 0.116 | 2.21199623137739e-09 | Cluster 5 | Tap1 |
| Psmb8 | 7.62935114493406e-14 | 0.931057388460257 | 0.822 | 0.426 | 2.36914241103637e-09 | Cluster 5 | Psmb8 |
| Nmi | 8.16393934378743e-14 | 0.692507080358969 | 0.567 | 0.171 | 2.53514808442631e-09 | Cluster 5 | Nmi |
| Ogfr | 3.65357327184302e-13 | 0.889980856620657 | 0.622 | 0.16 | 1.13454410810541e-08 | Cluster 5 | Ogfr |
| Usp25 | 4.27544242326331e-13 | 0.810058576110983 | 0.689 | 0.27 | 1.32765313569596e-08 | Cluster 5 | Usp25 |
| Trim30d | 1.03744000871072e-12 | 0.860086765556832 | 0.833 | 0.369 | 3.2215624590494e-08 | Cluster 5 | Trim30d |
| Parp12 | 1.09421527405796e-12 | 0.74721337848218 | 0.411 | 0.059 | 3.39786669053219e-08 | Cluster 5 | Parp12 |
| Chmp4b | 1.11646632358334e-12 | 0.625325838334011 | 0.956 | 0.769 | 3.46696287462334e-08 | Cluster 5 | Chmp4b |
| Lgals9 | 1.18240755747512e-12 | 0.832924558657635 | 0.644 | 0.197 | 3.67173018822749e-08 | Cluster 5 | Lgals9 |
| Cxcl101 | 1.33990678420489e-12 | 2.38954283137602 | 0.478 | 0.099 | 4.16081253699144e-08 | Cluster 5 | Cxcl10 |
| Nampt | 1.43497420267319e-12 | 0.945321690060272 | 0.522 | 0.17 | 4.45602539156107e-08 | Cluster 5 | Nampt |
| Tor1aip1 | 1.5396386794497e-12 | 0.821721461527161 | 0.756 | 0.315 | 4.78103999129517e-08 | Cluster 5 | Tor1aip1 |
| Trim30b3 | 2.05321825253628e-12 | 0.928367196740767 | 0.789 | 0.34 | 6.3758586396009e-08 | Cluster 5 | Trim30b |
| Tor3a | 5.21717872120934e-12 | 0.796961691996808 | 0.456 | 0.062 | 1.62009050829714e-07 | Cluster 5 | Tor3a |
| Sp110 | 5.48979034462182e-12 | 0.762821295675204 | 0.489 | 0.135 | 1.70474459571541e-07 | Cluster 5 | Sp110 |
| Psmb9 | 2.13411022437368e-11 | 0.749775883304829 | 0.633 | 0.202 | 6.6270524797476e-07 | Cluster 5 | Psmb9 |
| Zbp13 | 2.15174968453249e-11 | 1.10586987986766 | 0.478 | 0.095 | 6.68182829537874e-07 | Cluster 5 | Zbp1 |
| 1600014C10Rik | 2.66984184689981e-11 | 0.885626972037547 | 0.689 | 0.262 | 8.29065988717798e-07 | Cluster 5 | 1600014C10Rik |
| Ifih1 | 2.7986192112144e-11 | 0.938626263237653 | 0.489 | 0.072 | 8.69055223658409e-07 | Cluster 5 | Ifih1 |
| Oas2 | 4.41953522406849e-11 | 0.823539894304872 | 0.489 | 0.084 | 1.37239827312999e-06 | Cluster 5 | Oas2 |
| Ifit1bl2 | 6.0776718611794e-11 | 0.694194009542209 | 0.378 | 0.05 | 1.88729944305204e-06 | Cluster 5 | Ifit1bl2 |
| Dhx58 | 8.88261911863487e-11 | 0.44832885964121 | 0.256 | 0.034 | 2.75831971490969e-06 | Cluster 5 | Dhx58 |
| Helz21 | 1.18361533963424e-10 | 0.771845560037202 | 0.456 | 0.09 | 3.67548071416621e-06 | Cluster 5 | Helz2 |
| Ctss | 1.39591087390848e-10 | 0.87511709612776 | 0.856 | 0.551 | 4.334722036748e-06 | Cluster 5 | Ctss |
| Eif2ak2 | 4.2486694186598e-10 | 0.802815213158363 | 0.411 | 0.071 | 1.31933931457643e-05 | Cluster 5 | Eif2ak2 |
| Samhd13 | 5.27665570650379e-10 | 0.775507690607678 | 0.911 | 0.598 | 1.63855989654062e-05 | Cluster 5 | Samhd1 |
| Xaf11 | 5.31681472708856e-10 | 0.78169405461014 | 0.411 | 0.068 | 1.65103047720281e-05 | Cluster 5 | Xaf1 |
| Mitd1 | 6.07757896779389e-10 | 0.673978669846789 | 0.4 | 0.078 | 1.88727059686904e-05 | Cluster 5 | Mitd1 |
| Gbp5 | 7.24622731633112e-10 | 0.956939788350155 | 0.322 | 0.027 | 2.2501709685403e-05 | Cluster 5 | Gbp5 |
| Parp9 | 7.60911117802364e-10 | 0.60570684385951 | 0.356 | 0.046 | 2.36285729411168e-05 | Cluster 5 | Parp9 |
| Zufsp | 1.58624852037159e-09 | 0.719306123153165 | 0.289 | 0.039 | 4.9257775303099e-05 | Cluster 5 | Zufsp |
| Trim30c | 1.64138172312748e-09 | 0.501310797468332 | 0.244 | 0.01 | 5.09698266482776e-05 | Cluster 5 | Trim30c |
| Parp10 | 1.76530447721384e-09 | 0.70578114276172 | 0.456 | 0.097 | 5.48179999309214e-05 | Cluster 5 | Parp10 |
| Ifitm63 | 2.67057881058658e-09 | 0.979586531534028 | 0.511 | 0.147 | 8.29294838051451e-05 | Cluster 5 | Ifitm6 |
| Rnf114 | 5.23849581233647e-09 | 0.59571388097707 | 0.467 | 0.184 | 0.000162671010460484 | Cluster 5 | Rnf114 |
| Trim26 | 7.20789189473704e-09 | 0.509424776101486 | 0.356 | 0.089 | 0.000223826667007269 | Cluster 5 | Trim26 |
| Trim25 | 7.74898809872359e-09 | 0.543746798079664 | 0.611 | 0.316 | 0.000240629327429664 | Cluster 5 | Trim25 |
| Oas1a | 8.46318580352016e-09 | 0.504113272994745 | 0.289 | 0.033 | 0.000262807308756712 | Cluster 5 | Oas1a |
| Trim34a | 9.06962494865266e-09 | 0.535140457632143 | 0.311 | 0.034 | 0.000281639063530511 | Cluster 5 | Trim34a |
| Clec2d1 | 1.48940984517971e-08 | 0.76366515149627 | 0.878 | 0.559 | 0.000462506439223657 | Cluster 5 | Clec2d |
| Znfx1 | 1.73270683742162e-08 | 0.657487857088452 | 0.344 | 0.077 | 0.000538057454224537 | Cluster 5 | Znfx1 |
| H2-Q4 | 2.01446735782422e-08 | 0.640271027824741 | 0.489 | 0.124 | 0.000625552548625154 | Cluster 5 | H2-Q4 |

| | | | | | | | |
|---|---|---|---|---|---|---|---|
| Pnp1 | 2.25825780604212e-08 | 1.00301967721991 | 0.756 | 0.489 | 0.000701256796510259 | Cluster 5 | Pnp |
| Ube2l6 | 2.52176030135393e-08 | 0.601930507957879 | 0.333 | 0.073 | 0.000783082226379436 | Cluster 5 | Ube2l6 |
| Ifi206 | 3.00235490969481e-08 | 0.69871958083585 | 0.211 | 0.012 | 0.000932321270107529 | Cluster 5 | Ifi206 |
| H2-D13 | 3.19937204286246e-08 | 0.488119406422406 | 1 | 0.957 | 0.000993501000470081 | Cluster 5 | H2-D1 |
| Adar | 6.16401269490899e-08 | 0.54021400386234 | 0.389 | 0.103 | 0.00191411086215009 | Cluster 5 | Adar |
| Fcgr1 | 7.4879512830445e-08 | 1.01204425765205 | 0.344 | 0.078 | 0.00232523351192381 | Cluster 5 | Fcgr1 |
| Ifi208 | 8.16703991325882e-08 | 0.593202308543681 | 0.2 | 0.006 | 0.00253611090426426 | Cluster 5 | Ifi208 |
| Slfn9 | 9.0117289431554e-08 | 0.535242188276114 | 0.233 | 0.013 | 0.00279841218871805 | Cluster 5 | Slfn9 |
| Rap2c | 1.17112378395131e-07 | 0.455034876827611 | 0.544 | 0.316 | 0.00363669068630401 | Cluster 5 | Rap2c |
| Psme2b | 1.48364586701781e-07 | 0.502135968359363 | 0.3 | 0.04 | 0.0046071655108504 | Cluster 5 | Psme2b |
| Pttg1 | 2.11863389580219e-07 | 0.64525229591882 | 0.6 | 0.251 | 0.00657899383663454 | Cluster 5 | Pttg1 |
| Ogfrl11 | 2.23876353569508e-07 | 0.534757083975689 | 0.778 | 0.492 | 0.00695203240739395 | Cluster 5 | Ogfrl1 |
| Shisa5 | 2.71110422993202e-07 | 0.57237953653269 | 0.778 | 0.522 | 0.00841879196520791 | Cluster 5 | Shisa5 |
| Tspo1 | 3.11334000408881e-07 | 0.596703561256993 | 0.811 | 0.588 | 0.00966785471469697 | Cluster 5 | Tspo |
| Irgm2 | 3.42253692008319e-07 | 0.399840824753356 | 0.2 | 0.021 | 0.0106280038979343 | Cluster 5 | Irgm2 |
| Treml22 | 4.02931036685338e-07 | 0.575413534047162 | 0.422 | 0.132 | 0.0125122174821898 | Cluster 5 | Treml2 |
| Sppl2a | 8.47950018949063e-07 | 0.536229122448732 | 0.511 | 0.258 | 0.0263313919384252 | Cluster 5 | Sppl2a |
| Trim56 | 1.00431988952467e-06 | 0.498943214947767 | 0.367 | 0.1 | 0.0311871455294097 | Cluster 5 | Trim56 |
| Irf11 | 1.30195535723902e-06 | 0.703173181343957 | 0.678 | 0.35 | 0.0404296197083434 | Cluster 5 | Irf1 |
| Sgk3 | 1.36571255903757e-06 | 0.415806213143249 | 0.411 | 0.177 | 0.0424094720957936 | Cluster 5 | Sgk3 |
| Uba7 | 1.36730544391425e-06 | 0.409625065775086 | 0.278 | 0.04 | 0.0424589359498693 | Cluster 5 | Uba7 |
| Smchd1 | 1.41643973540525e-06 | 0.744740247949202 | 0.556 | 0.286 | 0.0439847031035391 | Cluster 5 | Smchd1 |
| Fcgr41 | 1.51992575247123e-06 | 0.704819667492756 | 0.589 | 0.281 | 0.047198254391489 | Cluster 5 | Fcgr4 |
| Stat2 | 2.27718224817492e-06 | 0.56383921595608 | 0.289 | 0.042 | 0.0707133403525758 | Cluster 5 | Stat2 |
| Cd274 | 3.04628134452564e-06 | 0.623708761733194 | 0.622 | 0.347 | 0.0945961745915547 | Cluster 5 | Cd274 |
| Sap30 | 3.55351027722468e-06 | 0.443798233947117 | 0.289 | 0.067 | 0.110347154638658 | Cluster 5 | Sap30 |
| Mov10 | 3.70661043022681e-06 | 0.301481297373547 | 0.211 | 0.024 | 0.115101373689833 | Cluster 5 | Mov10 |
| Slc15a34 | 3.74034290593901e-06 | 0.544620174409409 | 0.844 | 0.632 | 0.116148868258124 | Cluster 5 | Slc15a3 |
| Cxcr23 | 4.5904816687319e-06 | 0.346966971803212 | 0.833 | 0.638 | 0.142548227259132 | Cluster 5 | Cxcr2 |
| Mndal | 6.25185958636585e-06 | 0.604935837603114 | 0.2 | 0.03 | 0.194138995735419 | Cluster 5 | Mndal |
| Ifi27l2a | 6.73600567147963e-06 | 1.25681693175781 | 0.589 | 0.358 | 0.209173184116457 | Cluster 5 | Ifi27l2a |
| Psme21 | 7.35342439033015e-06 | 0.488932181717949 | 0.633 | 0.344 | 0.228345887592922 | Cluster 5 | Psme2 |
| Sdcbp | 7.38918073341849e-06 | 0.401208308343165 | 0.978 | 0.901 | 0.229456229314844 | Cluster 5 | Sdcbp |
| Gdap10 | 8.82788411344117e-06 | 0.551779513622572 | 0.333 | 0.104 | 0.274132285374689 | Cluster 5 | Gdap10 |
| Tbrg1 | 8.97954806908993e-06 | 0.56960707514645 | 0.411 | 0.155 | 0.27884190618945 | Cluster 5 | Tbrg1 |
| Psme1 | 9.29110940761741e-06 | 0.531112181461807 | 0.578 | 0.308 | 0.288516820434744 | Cluster 5 | Psme1 |
| Casp8 | 9.4702620772194e-06 | 0.511079518235497 | 0.344 | 0.141 | 0.294080048283894 | Cluster 5 | Casp8 |
| Hmox2 | 1.03331772038604e-05 | 0.536619314536007 | 0.4 | 0.154 | 0.320876151711478 | Cluster 5 | Hmox2 |
| Ufsp2 | 1.0921824704853e-05 | 0.259294402205066 | 0.222 | 0.094 | 0.339155422559801 | Cluster 5 | Ufsp2 |
| Ankfy1 | 1.30182258966777e-05 | 0.517392908140726 | 0.411 | 0.135 | 0.404254968769531 | Cluster 5 | Ankfy1 |
| D1Ertd622e | 1.30996738427887e-05 | 0.390359794188221 | 0.467 | 0.205 | 0.406784171840116 | Cluster 5 | D1Ertd622e |
| Ifi203 | 1.47790682084553e-05 | 0.643932554358421 | 0.189 | 0.027 | 0.458934405077163 | Cluster 5 | Ifi203 |
| Lipg1 | 1.99597678558116e-05 | 0.285290610522123 | 0.189 | 0.046 | 0.619810671226519 | Cluster 5 | Lipg |
| Fam46a | 2.01359299847018e-05 | 0.467930802481739 | 0.278 | 0.082 | 0.625281033814945 | Cluster 5 | Fam46a |
| Tmem184b | 2.05715859439831e-05 | 0.425439422996011 | 0.433 | 0.205 | 0.638809458318506 | Cluster 5 | Tmem184b |
| Apobec3 | 2.77868353089145e-05 | 0.417946942742748 | 0.311 | 0.075 | 0.862864596847723 | Cluster 5 | Apobec3 |
| Trim21 | 2.9297309115316e-05 | 0.276758767056878 | 0.133 | 0.006 | 0.909769339957908 | Cluster 5 | Trim21 |
| Taf7 | 3.12259414980483e-05 | 0.462371574898478 | 0.522 | 0.273 | 0.969659161338893 | Cluster 5 | Taf7 |
| Ly6i | 3.16932567936419e-05 | 0.470920597616398 | 0.233 | 0.055 | 0.984170703212962 | Cluster 5 | Ly6i |
| Ifit1bl1 | 3.21644891059616e-05 | 0.431277739932343 | 0.111 | 0.005 | 0.998803880207425 | Cluster 5 | Ifit1bl1 |
| Psmb101 | 3.35163455225558e-05 | 0.522992844596871 | 0.433 | 0.173 | 1 | Cluster 5 | Psmb10 |
| Il18bp | 3.90817880864454e-05 | 0.519794078580071 | 0.244 | 0.021 | 1 | Cluster 5 | Il18bp |
| Zc3hav1 | 4.29839805835613e-05 | 0.403380890870966 | 0.667 | 0.424 | 1 | Cluster 5 | Zc3hav1 |
| Daxx | 5.84566270034693e-05 | 0.433312615133083 | 0.211 | 0.043 | 1 | Cluster 5 | Daxx |
| Nsd31 | 6.04688543835156e-05 | 0.491518964471483 | 0.622 | 0.406 | 1 | Cluster 5 | Nsd3 |
| Dcp2 | 6.5748457450512e-05 | 0.35505389571392 | 0.222 | 0.069 | 1 | Cluster 5 | Dcp2 |
| Nt5c3 | 7.70844611516643e-05 | 0.512149705418093 | 0.278 | 0.097 | 1 | Cluster 5 | Nt5c3 |
| Casp4 | 7.71197155844605e-05 | 0.443434637605381 | 0.644 | 0.413 | 1 | Cluster 5 | Casp4 |
| B430306N03Rik | 8.03106026777525e-05 | 0.322030952711581 | 0.444 | 0.235 | 1 | Cluster 5 | B430306N03Rik |
| Nlrc5 | 9.35880999923256e-05 | 0.433571918607024 | 0.3 | 0.06 | 1 | Cluster 5 | Nlrc5 |
| Peli1 | 9.36977539180105e-05 | 0.505247601351841 | 0.533 | 0.323 | 1 | Cluster 5 | Peli1 |

| | | | | | | | |
|---|---|---|---|---|---|---|---|
| Papd7 | 0.000118130268487269 | 0.273045797129491 | 0.178 | 0.028 | 1 | Cluster 5 | Papd7 |
| H2-T24 | 0.00012196767118074 | 0.345352831668245 | 0.178 | 0.036 | 1 | Cluster 5 | H2-T24 |
| Prdx53 | 0.0001266518823278 | 0.262131112905177 | 0.944 | 0.894 | 1 | Cluster 5 | Prdx5 |
| Il151 | 0.000128535279845822 | 0.278271614401859 | 0.222 | 0.084 | 1 | Cluster 5 | Il15 |
| Stard3 | 0.000131911149580295 | 0.400466599730386 | 0.356 | 0.116 | 1 | Cluster 5 | Stard3 |
| Tdrd7 | 0.000148033055173246 | 0.403752281247234 | 0.267 | 0.066 | 1 | Cluster 5 | Tdrd7 |
| Irf2 | 0.000158764713096111 | 0.519448551126443 | 0.611 | 0.373 | 1 | Cluster 5 | Irf2 |
| Acod12 | 0.000198007879999764 | 0.449563929404477 | 0.933 | 0.82 | 1 | Cluster 5 | Acod1 |
| Cd471 | 0.000199480088392605 | 0.398767016441767 | 0.8 | 0.64 | 1 | Cluster 5 | Cd47 |
| Fgl2 | 0.000222992090390649 | 0.731339392527034 | 0.733 | 0.514 | 1 | Cluster 5 | Fgl2 |
| Ifi35 | 0.000234257961495027 | 0.510073327616075 | 0.478 | 0.221 | 1 | Cluster 5 | Ifi35 |
| Ccrl22 | 0.000234308987520374 | 0.483223889550925 | 0.922 | 0.897 | 1 | Cluster 5 | Ccrl2 |
| Mxd14 | 0.000240040876105304 | 0.401745407766144 | 0.956 | 0.871 | 1 | Cluster 5 | Mxd1 |
| Coq2 | 0.000277709145441589 | 0.310351517081336 | 0.178 | 0.046 | 1 | Cluster 5 | Coq2 |
| Gm12216 | 0.000301429380808847 | 0.261659968518667 | 0.167 | 0.041 | 1 | Cluster 5 | Gm12216 |
| Tapbp | 0.000341142140200892 | 0.367836383895168 | 0.767 | 0.564 | 1 | Cluster 5 | Tapbp |
| Map2k11 | 0.00038784769673203 | 0.349880575896259 | 0.4 | 0.183 | 1 | Cluster 5 | Map2k1 |
| Ascc3 | 0.000407464511857355 | 0.456203047473204 | 0.511 | 0.287 | 1 | Cluster 5 | Ascc3 |
| AW011738 | 0.000410794479602164 | 0.30888552386055 | 0.2 | 0.063 | 1 | Cluster 5 | AW011738 |
| Gbp9 | 0.000420546592118498 | 0.367994376188409 | 0.189 | 0.036 | 1 | Cluster 5 | Gbp9 |
| Hcar21 | 0.000646871500451041 | 0.31933601252345 | 0.878 | 0.741 | 1 | Cluster 5 | Hcar2 |
| Hck | 0.000711982935187705 | 0.349792078250242 | 0.533 | 0.365 | 1 | Cluster 5 | Hck |
| Csrnp1 | 0.000725182045059121 | 0.284429485046435 | 0.9 | 0.795 | 1 | Cluster 5 | Csrnp1 |
| Arl6ip1 | 0.000751481886002753 | 0.418885597628686 | 0.711 | 0.493 | 1 | Cluster 5 | Arl6ip1 |
| Clic4 | 0.000890900862150905 | 0.537803255798523 | 0.356 | 0.175 | 1 | Cluster 5 | Clic4 |
| Atp8b4 | 0.000894963578862801 | 0.346464385142683 | 0.367 | 0.208 | 1 | Cluster 5 | Atp8b4 |
| Socs1 | 0.000910001484880336 | 0.29186011730567 | 0.2 | 0.071 | 1 | Cluster 5 | Socs1 |
| Elf1 | 0.000914675583264628 | 0.276387110480516 | 0.378 | 0.23 | 1 | Cluster 5 | Elf1 |
| Rnf34 | 0.000946846222093846 | 0.40520152211193 | 0.333 | 0.126 | 1 | Cluster 5 | Rnf34 |
| Dusp1 | 0.000948925132454983 | 0.3593705778606 | 0.989 | 0.939 | 1 | Cluster 5 | Dusp1 |
| Morc3 | 0.0011166597439266 | 0.434109450150049 | 0.356 | 0.172 | 1 | Cluster 5 | Morc3 |
| Ccnyl1 | 0.00118099833336017 | 0.347539471209769 | 0.233 | 0.098 | 1 | Cluster 5 | Ccnyl1 |
| Sp1404 | 0.00119866090314555 | 0.416581213501463 | 0.533 | 0.329 | 1 | Cluster 5 | Sp140 |
| Tmbim6 | 0.00123132019714781 | 0.252714551747111 | 0.933 | 0.882 | 1 | Cluster 5 | Tmbim6 |
| Itm2b1 | 0.00125349896351768 | 0.275942846279891 | 0.978 | 0.9 | 1 | Cluster 5 | Itm2b |
| Gm21188 | 0.0013600298204787 | 0.480588951449215 | 0.467 | 0.204 | 1 | Cluster 5 | Gm21188 |
| Npc2 | 0.00157528117287477 | 0.47469756124468 | 0.678 | 0.465 | 1 | Cluster 5 | Npc2 |
| Tap2 | 0.00185563319627513 | 0.325774547235577 | 0.467 | 0.237 | 1 | Cluster 5 | Tap2 |
| Prr5l | 0.00215827490303143 | 0.291219393488148 | 0.178 | 0.046 | 1 | Cluster 5 | Prr5l |
| Zfp361 | 0.00219035456575302 | 0.321745049001997 | 0.944 | 0.897 | 1 | Cluster 5 | Zfp36 |
| Fcgr31 | 0.00219227850873397 | 0.26629744383853 | 0.667 | 0.617 | 1 | Cluster 5 | Fcgr3 |
| Hsh2d | 0.00236329781803801 | 0.289779186876156 | 0.2 | 0.03 | 1 | Cluster 5 | Hsh2d |
| Slfn21 | 0.00243951911496488 | 0.36588034216324 | 0.978 | 0.916 | 1 | Cluster 5 | Slfn2 |
| Pxk | 0.00325552899192791 | 0.3745059055344 | 0.356 | 0.165 | 1 | Cluster 5 | Pxk |
| Il18 | 0.00333544164496064 | 0.322160794199901 | 0.167 | 0.037 | 1 | Cluster 5 | Il18 |
| Myd88 | 0.00347365276077682 | 0.378704273457425 | 0.456 | 0.338 | 1 | Cluster 5 | Myd88 |
| A530064D06Rik3 | 0.00356363389379648 | 0.308843009590521 | 0.278 | 0.1 | 1 | Cluster 5 | A530064D06Rik |
| Ly6c23 | 0.00359290466023198 | 1.48713394469599 | 0.256 | 0.125 | 1 | Cluster 5 | Ly6c2 |
| Mb21d11 | 0.00361528658982913 | 0.330061595606528 | 0.311 | 0.155 | 1 | Cluster 5 | Mb21d1 |
| Tlr24 | 0.00547383153623517 | 0.363716576598603 | 0.622 | 0.4 | 1 | Cluster 5 | Tlr2 |
| Ube2q1 | 0.00616604932984509 | 0.2591244172405 | 0.389 | 0.245 | 1 | Cluster 5 | Ube2q1 |
| Zfp281 | 0.00625588494632567 | 0.256947837819844 | 0.289 | 0.157 | 1 | Cluster 5 | Zfp281 |
| Unc93b11 | 0.00694144958484628 | 0.265089712552024 | 0.667 | 0.529 | 1 | Cluster 5 | Unc93b1 |
| H2-Q72 | 0.00709321033438526 | 0.386246847129392 | 0.278 | 0.133 | 1 | Cluster 5 | H2-Q7 |
| Whamm | 0.00790268488927918 | 0.274292737625518 | 0.167 | 0.042 | 1 | Cluster 5 | Whamm |
| Atf3 | 0.00797651806653567 | 0.337246993506524 | 0.656 | 0.478 | 1 | Cluster 5 | Atf3 |
| Jaml2 | 0.00950455071403311 | 0.491478077620526 | 0.289 | 0.179 | 1 | Cluster 5 | Jaml |
| Pdcd1lg2 | 1.07997420895189e-23 | 1.75656268011855 | 0.704 | 0.028 | 3.35364391105829e-19 | Cluster 6 | Pdcd1lg2 |
| Nceh1 | 8.08559865145651e-21 | 1.54106437125572 | 0.741 | 0.058 | 2.51082094923679e-16 | Cluster 6 | Nceh1 |
| Plgrkt | 1.35730793481241e-19 | 1.52050215021237 | 0.889 | 0.166 | 4.21484832997297e-15 | Cluster 6 | Plgrkt |
| Mreg | 1.65439291145226e-19 | 1.09721533922197 | 0.63 | 0.049 | 5.13738630793271e-15 | Cluster 6 | Mreg |

| | | | | | | | |
|---|---|---|---|---|---|---|---|
| Fnip2 | 4.64261883251868e-18 | 1.49126404986286 | 0.852 | 0.118 | 1.44167242606203e-13 | Cluster 6 | Fnip2 |
| Slc26a11 | 6.43673100347331e-18 | 1.97481335631573 | 0.63 | 0.065 | 1.99879807850857e-13 | Cluster 6 | Slc26a11 |
| Itpr2 | 2.40162074911293e-17 | 2.09968117460833 | 0.815 | 0.226 | 7.45775291222037e-13 | Cluster 6 | Itpr2 |
| Naglu | 3.29047092771385e-17 | 1.14363477699201 | 0.556 | 0.031 | 1.02178993718298e-12 | Cluster 6 | Naglu |
| Gstm1 | 5.12277843577126e-17 | 1.6519514576314 | 0.593 | 0.072 | 1.59077638766005e-12 | Cluster 6 | Gstm1 |
| Aph1c | 5.86797400224253e-17 | 1.55995619521018 | 0.63 | 0.081 | 1.82218196691637e-12 | Cluster 6 | Aph1c |
| Ctsb1 | 1.62536393342969e-15 | 1.22406311544356 | 1 | 0.932 | 5.04724262247921e-11 | Cluster 6 | Ctsb |
| Atp6v1c11 | 6.53949188825064e-15 | 1.44564298672533 | 0.926 | 0.29 | 2.03070841605847e-10 | Cluster 6 | Atp6v1c1 |
| Smad3 | 1.24042602818278e-14 | 1.16626752479356 | 0.556 | 0.067 | 3.85189494531598e-10 | Cluster 6 | Smad3 |
| Atp6v1a1 | 1.27542181963423e-14 | 1.68147469433439 | 1 | 0.38 | 3.96056737651018e-10 | Cluster 6 | Atp6v1a |
| Cstb | 3.27929840496027e-14 | 1.7526552242566 | 1 | 0.874 | 1.01832053369231e-09 | Cluster 6 | Cstb |
| Gpr137b | 4.23976981356826e-14 | 1.31814716048643 | 0.741 | 0.155 | 1.31657572020735e-09 | Cluster 6 | Gpr137b |
| Slc31a2 | 5.46457524774375e-14 | 1.29184595798872 | 0.815 | 0.224 | 1.69691455168187e-09 | Cluster 6 | Slc31a2 |
| Ftl12 | 5.95270275234303e-14 | 1.47682209711806 | 1 | 1 | 1.84849278568508e-09 | Cluster 6 | Ftl1 |
| Cd68 | 2.9257738089929e-13 | 1.42569336222852 | 0.963 | 0.464 | 9.08540540906566e-09 | Cluster 6 | Cd68 |
| Psap | 3.20134416848059e-13 | 1.95521018868231 | 1 | 0.829 | 9.94113404638279e-09 | Cluster 6 | Psap |
| Gm13391 | 5.01508735666601e-13 | 1.03089755264537 | 0.37 | 0.009 | 1.5573350768655e-08 | Cluster 6 | Gm13391 |
| Rilpl2 | 1.25359019319184e-12 | 1.83288916511951 | 0.778 | 0.217 | 3.89277362691861e-08 | Cluster 6 | Rilpl2 |
| Lamp11 | 1.14490857204166e-11 | 1.24039844802889 | 1 | 0.675 | 3.55528458876096e-07 | Cluster 6 | Lamp1 |
| Tmem140 | 5.73544937387269e-11 | 1.04610825653043 | 0.407 | 0.036 | 1.78102909406869e-06 | Cluster 6 | Tmem140 |
| Bri31 | 9.45659200553440e-11 | 1.20376592372811 | 1 | 0.779 | 2.9365555154786e-06 | Cluster 6 | Bri3 |
| Tecpr1 | 9.9434895929473e-11 | 1.35390218634543 | 0.444 | 0.049 | 3.08775182329793e-06 | Cluster 6 | Tecpr1 |
| Renbp | 1.05454661909121e-10 | 0.893996396067519 | 0.37 | 0.041 | 3.27468361626392e-06 | Cluster 6 | Renbp |
| Ctsd3 | 1.39022759662201e-10 | 1.23258281474152 | 1 | 0.933 | 4.31707375579033e-06 | Cluster 6 | Ctsd |
| Ftl1-ps1 | 6.08413418705248e-10 | 0.970692188326351 | 0.852 | 0.326 | 1.88930618910541e-05 | Cluster 6 | Ftl1-ps1 |
| Ankrd12 | 8.4781748026964e-10 | 1.82354441548476 | 0.852 | 0.35 | 2.63272762148131e-05 | Cluster 6 | Ankrd12 |
| Amdhd21 | 2.15607686328868e-09 | 1.22376645957357 | 0.667 | 0.189 | 6.69526548357033e-05 | Cluster 6 | Amdhd2 |
| Osbpl8 | 2.69906580495632e-09 | 1.25447351821216 | 0.704 | 0.189 | 8.38140904413087e-05 | Cluster 6 | Osbpl8 |
| Ccdc126 | 3.6956239243183e-09 | 0.900177022608299 | 0.444 | 0.062 | 0.000114760209721856 | Cluster 6 | Ccdc126 |
| Tpp11 | 3.84975555292243e-09 | 0.862268983031542 | 0.704 | 0.231 | 0.0001195464591849 | Cluster 6 | Tpp1 |
| Cipc | 5.62989338289749e-09 | 0.561976123652664 | 0.296 | 0.033 | 0.000174825079219116 | Cluster 6 | Cipc |
| Tns3 | 7.63841209791898e-09 | 0.983277030308975 | 0.481 | 0.072 | 0.000237195610876678 | Cluster 6 | Tns3 |
| Gdf15 | 9.65688833444994e-09 | 2.38133346716128 | 0.407 | 0.06 | 0.000299875353449674 | Cluster 6 | Gdf15 |
| Dhrs3 | 1.02680621969309e-08 | 0.753630709261069 | 0.519 | 0.104 | 0.000318854135401294 | Cluster 6 | Dhrs3 |
| Creg11 | 1.16676540185191e-08 | 1.92101782505519 | 0.852 | 0.461 | 0.000362315660237075 | Cluster 6 | Creg1 |
| Mbnl2 | 1.21231518818506e-08 | 1.00003572406793 | 0.926 | 0.541 | 0.000376460235387105 | Cluster 6 | Mbnl2 |
| Hexa1 | 1.28019587847472e-08 | 0.956385542504248 | 0.889 | 0.398 | 0.000397539226142754 | Cluster 6 | Hexa |
| 1700017B05Rik | 1.39533297385768e-08 | 0.895341554914823 | 0.741 | 0.263 | 0.000433292748372025 | Cluster 6 | 1700017B05Rik |
| Atp6v1d1 | 1.4027208640375e-08 | 1.01365939914461 | 0.778 | 0.332 | 0.000435586909909564 | Cluster 6 | Atp6v1d |
| Gabarap | 1.69488704887837e-08 | 0.63927965008442 | 1 | 0.979 | 0.0005263132752882 | Cluster 6 | Gabarap |
| Npc1 | 2.50047696077668e-08 | 0.767048036622051 | 0.593 | 0.151 | 0.000776473110629981 | Cluster 6 | Npc1 |
| Cd93 | 2.70904313021129e-08 | 0.950040072503682 | 1 | 0.932 | 0.000841239163224512 | Cluster 6 | Cd9 |
| Gla | 2.98978596088122e-08 | 1.11130342912102 | 0.852 | 0.336 | 0.000928418234432446 | Cluster 6 | Gla |
| Gns1 | 3.11699850022806e-08 | 1.26568663027047 | 0.741 | 0.277 | 0.000967921544275819 | Cluster 6 | Gns |
| Fth12 | 5.81912430243358e-08 | 0.849495085637685 | 1 | 1 | 0.0018070126696347 | Cluster 6 | Fth1 |
| Vps39 | 8.6351613993817e-08 | 0.448052533742423 | 0.259 | 0.036 | 0.00268147666935 | Cluster 6 | Vps39 |
| Tst | 1.01740726914896e-07 | 0.591727914437337 | 0.185 | 0.005 | 0.00315935479288828 | Cluster 6 | Tst |
| Spdl1 | 1.1305814542356e-07 | 0.615370678828713 | 0.222 | 0.027 | 0.00351079458983781 | Cluster 6 | Spdl1 |
| Arfgef2 | 1.23245105570329e-07 | 0.917567510540033 | 0.407 | 0.066 | 0.00382713026327544 | Cluster 6 | Arfgef2 |
| Wdfy1 | 1.28440854438511e-07 | 0.456467139283021 | 0.407 | 0.088 | 0.00398847385287908 | Cluster 6 | Wdfy1 |
| Gclm | 1.70227695793566e-07 | 1.08134252602515 | 0.519 | 0.127 | 0.00528608063747759 | Cluster 6 | Gclm |
| Mpp1 | 1.91009376006252e-07 | 0.811224229167808 | 0.519 | 0.16 | 0.00593141415312213 | Cluster 6 | Mpp1 |
| Znrf1 | 1.91916253683838e-07 | 0.853819013749448 | 0.593 | 0.149 | 0.00595957542564421 | Cluster 6 | Znrf1 |
| Cd632 | 2.02401725972911e-07 | 1.8594743604983 | 0.852 | 0.678 | 0.00628518079663681 | Cluster 6 | Cd63 |
| Fuca2 | 2.42784110494522e-07 | 1.06906878705765 | 0.852 | 0.407 | 0.00753917498318641 | Cluster 6 | Fuca2 |
| Msrb13 | 2.4729044115341e-07 | 0.735709183951905 | 0.963 | 0.903 | 0.00767911006913685 | Cluster 6 | Msrb1 |
| Gnpda11 | 2.48005510899515e-07 | 0.753132392689263 | 0.63 | 0.178 | 0.00770131512996264 | Cluster 6 | Gnpda1 |
| Gde1 | 2.57838306248945e-07 | 0.874407383125383 | 0.556 | 0.163 | 0.00800665292394849 | Cluster 6 | Gde1 |
| Syne1 | 2.66857805258978e-07 | 0.726846551295767 | 0.37 | 0.084 | 0.00828673542670705 | Cluster 6 | Syne1 |
| Dtnbp1 | 2.84185866237217e-07 | 1.08339742860105 | 0.593 | 0.15 | 0.0088248237042643 | Cluster 6 | Dtnbp1 |
| 9930022D16Rik | 2.96377913429986e-07 | 0.664608127891142 | 0.296 | 0.039 | 0.00920342334574134 | Cluster 6 | 9930022D16Rik |

| | | | | | | | |
|---|---|---|---|---|---|---|---|
| Soat1 | 3.28843291898269e-07 | 0.944589375704528 | 0.63 | 0.234 | 0.010211570743317 | Cluster 6 | Soat1 |
| Lamtor11 | 3.34296536965799e-07 | 0.766333633211299 | 0.889 | 0.43 | 0.010380910362399 | Cluster 6 | Lamtor1 |
| Naa501 | 3.91516268849491e-07 | 0.928749229552959 | 0.63 | 0.211 | 0.0121577546965832 | Cluster 6 | Naa50 |
| Spsb2 | 4.2947376170935e-07 | 0.509402368035558 | 0.333 | 0.049 | 0.0133364487223604 | Cluster 6 | Spsb2 |
| Bax3 | 4.93297051117768e-07 | 0.890337506602325 | 0.778 | 0.378 | 0.0153183533283601 | Cluster 6 | Bax |
| Ap3s1 | 6.95790531879169e-07 | 0.703013248573593 | 0.704 | 0.244 | 0.0216063833864438 | Cluster 6 | Ap3s1 |
| Gm20604 | 7.68406037984916e-07 | 0.749128737695346 | 0.37 | 0.07 | 0.0238613126975456 | Cluster 6 | Gm20604 |
| Sc5d | 9.03403306540453e-07 | 0.768090376974463 | 0.444 | 0.096 | 0.0280533828780007 | Cluster 6 | Sc5d |
| Fnip1 | 9.33124163991179e-07 | 0.861745901595047 | 0.815 | 0.372 | 0.0289763046644181 | Cluster 6 | Fnip1 |
| Lrrfip2 | 9.76873657491577e-07 | 0.839077354552981 | 0.704 | 0.258 | 0.030334857686086 | Cluster 6 | Lrrfip2 |
| Hip1r | 9.94561892237404e-07 | 0.731309391486838 | 0.333 | 0.053 | 0.0308841304396481 | Cluster 6 | Hip1r |
| Nhlrc3 | 1.18762233973388e-06 | 0.57463444703037 | 0.444 | 0.084 | 0.0368792365157562 | Cluster 6 | Nhlrc3 |
| Gas2l3 | 1.22868981063028e-06 | 0.681996179180058 | 0.444 | 0.071 | 0.038154504689502 | Cluster 6 | Gas2l3 |
| Tpd52l2 | 1.42361743309183e-06 | 0.534611779265444 | 0.37 | 0.121 | 0.0442075921498006 | Cluster 6 | Tpd52l2 |
| Mafg1 | 1.57200957418934e-06 | 0.949319704226372 | 0.815 | 0.36 | 0.0488156133073016 | Cluster 6 | Mafg |
| Uap1l1 | 1.75462998929218e-06 | 0.838740485333607 | 0.407 | 0.083 | 0.0544865250574902 | Cluster 6 | Uap1l1 |
| 4833407H14Rik | 1.80328419028532e-06 | 0.807923859542242 | 0.889 | 0.483 | 0.0559973839609299 | Cluster 6 | 4833407H14Rik |
| Tlr6 | 1.99150253417737e-06 | 0.580120844927063 | 0.444 | 0.122 | 0.0618421281938099 | Cluster 6 | Tlr6 |
| Sqstm11 | 1.99830235910927e-06 | 0.854980187135652 | 0.926 | 0.663 | 0.0620532831574202 | Cluster 6 | Sqstm1 |
| Hmox11 | 2.00849887325077e-06 | 1.00313664815299 | 0.852 | 0.557 | 0.0623699155110563 | Cluster 6 | Hmox1 |
| Itgb3 | 2.07037627406964e-06 | 0.538503373046539 | 0.148 | 0.008 | 0.0642913944386847 | Cluster 6 | Itgb3 |
| Gucy2c | 2.07037627406964e-06 | 0.415658848575237 | 0.148 | 0.006 | 0.0642913944386847 | Cluster 6 | Gucy2c |
| D630024D03Rik | 2.07037627406964e-06 | 0.290314154551897 | 0.148 | 0.001 | 0.0642913944386847 | Cluster 6 | D630024D03Rik |
| Ncoa3 | 2.14636968570919e-06 | 0.598396124441174 | 0.556 | 0.208 | 0.0666512178503274 | Cluster 6 | Ncoa3 |
| Lrrc8a | 2.23981697713429e-06 | 0.482725265492972 | 0.185 | 0.028 | 0.0695530365909511 | Cluster 6 | Lrrc8a |
| Coro1c | 2.54897007823435e-06 | 0.619932211045076 | 0.556 | 0.192 | 0.0791531678394113 | Cluster 6 | Coro1c |
| Dedd2 | 3.0906523471002e-06 | 0.782087426217875 | 0.852 | 0.355 | 0.0959740273345026 | Cluster 6 | Dedd2 |
| Slc43a2 | 3.61630186053148e-06 | 0.78645722878171 | 0.63 | 0.197 | 0.112297021675084 | Cluster 6 | Slc43a2 |
| Ergic1 | 3.67792382028681e-06 | 0.636345411023127 | 0.519 | 0.135 | 0.114210568391366 | Cluster 6 | Ergic1 |
| Stxbp5 | 3.82667695026629e-06 | 0.737305131664234 | 0.37 | 0.07 | 0.118829799336619 | Cluster 6 | Stxbp5 |
| Impact1 | 3.90507240835853e-06 | 1.06163607258195 | 0.481 | 0.115 | 0.121264213496758 | Cluster 6 | Impact |
| Itpr1 | 3.96596679312046e-06 | 0.58509061706681 | 0.37 | 0.057 | 0.12315516682677 | Cluster 6 | Itpr1 |
| Laptm4a | 4.48203256811865e-06 | 0.421138506932594 | 0.481 | 0.232 | 0.139180557337789 | Cluster 6 | Laptm4a |
| Prok2 | 4.48663408724477e-06 | 1.20413579542123 | 0.259 | 0.017 | 0.139323448311212 | Cluster 6 | Prok2 |
| Tmem1891 | 4.52647697153886e-06 | 0.608558235426933 | 0.852 | 0.393 | 0.140560689397196 | Cluster 6 | Tmem189 |
| Por1 | 4.5460135564307e-06 | 0.951718307635437 | 0.741 | 0.334 | 0.141167358967843 | Cluster 6 | Por |
| Rbm381 | 4.61622109828024e-06 | 0.924784454356495 | 0.63 | 0.221 | 0.143347513764896 | Cluster 6 | Rbm38 |
| F101 | 4.73942820007292e-06 | 0.966438118462207 | 0.556 | 0.183 | 0.147173463896864 | Cluster 6 | F10 |
| Plpbp | 5.15639474734073e-06 | 0.928536661396779 | 0.519 | 0.1 | 0.160121526089172 | Cluster 6 | Plpbp |
| Atp6v0b | 5.81325627789714e-06 | 0.753622636212324 | 1 | 0.844 | 0.18051904719754 | Cluster 6 | Atp6v0b |
| Rragc | 6.75621324165697e-06 | 0.773211930349037 | 0.704 | 0.285 | 0.209800689793174 | Cluster 6 | Rragc |
| Rnh1 | 6.77534805833612e-06 | 1.52396978637789 | 0.778 | 0.461 | 0.210394883255512 | Cluster 6 | Rnh1 |
| Osgin11 | 8.1987769239637e-06 | 1.31857522905614 | 0.593 | 0.201 | 0.254596619819845 | Cluster 6 | Osgin1 |
| Sh2d3c2 | 8.78172247165147e-06 | 0.932723241924607 | 0.556 | 0.155 | 0.272698827912193 | Cluster 6 | Sh2d3c |
| Slc40a11 | 8.92882824204014e-06 | 0.894954119089755 | 0.444 | 0.124 | 0.277266903400072 | Cluster 6 | Slc40a1 |
| Dgkz | 9.51420111112839e-06 | 0.619579657582437 | 0.556 | 0.237 | 0.29544448710387 | Cluster 6 | Dgkz |
| Idua | 1.0504944933129e-05 | 0.561038870735434 | 0.333 | 0.055 | 0.326210055008454 | Cluster 6 | Idua |
| Atp6v0e1 | 1.28248968031561e-05 | 0.643778541248691 | 0.963 | 0.807 | 0.398251520428407 | Cluster 6 | Atp6v0e |
| Lasp1 | 1.29714451274278e-05 | 0.803099133940122 | 0.889 | 0.519 | 0.402802285542015 | Cluster 6 | Lasp1 |
| Ctdp1 | 1.49799736293128e-05 | 0.448467712071501 | 0.296 | 0.075 | 0.46517312111105 | Cluster 6 | Ctdp1 |
| Slc23a2 | 1.54089363227744e-05 | 0.5665385861274 | 0.407 | 0.095 | 0.478493699631112 | Cluster 6 | Slc23a2 |
| Snx8 | 1.77688530179692e-05 | 0.572132218223835 | 0.444 | 0.084 | 0.551776192766998 | Cluster 6 | Snx8 |
| Slc20a1 | 1.81664016646881e-05 | 1.0614070923435 | 0.519 | 0.222 | 0.564121270893559 | Cluster 6 | Slc20a1 |
| Atf31 | 1.90252206770524e-05 | 1.11180556446721 | 0.926 | 0.481 | 0.590790177684509 | Cluster 6 | Atf3 |
| Clcn7 | 1.98489984737816e-05 | 0.521610068051538 | 0.444 | 0.139 | 0.616370949606339 | Cluster 6 | Clcn7 |
| Zfp110 | 1.99346234064653e-05 | 0.517118402833416 | 0.296 | 0.067 | 0.619029860640968 | Cluster 6 | Zfp110 |
| Cerk | 2.07026288819616e-05 | 0.610365467587911 | 0.296 | 0.057 | 0.642878734671554 | Cluster 6 | Cerk |
| Ly96 | 2.2323968703384e-05 | 0.524458659967586 | 0.296 | 0.085 | 0.693226200146182 | Cluster 6 | Ly96 |
| Tmem202 | 2.41113430211483e-05 | 0.512643919614331 | 0.185 | 0.021 | 0.748729534835718 | Cluster 6 | Tmem202 |
| Vps13d | 2.48422175470729e-05 | 0.679917968112526 | 0.407 | 0.129 | 0.771425381489253 | Cluster 6 | Vps13d |
| B4galt5 | 2.51211378244748e-05 | 0.445133883379908 | 0.37 | 0.088 | 0.780086692863416 | Cluster 6 | B4galt5 |

| | | | | | | | |
|---|---|---|---|---|---|---|---|
| Akr1a1 | 2.9584856823167e-05 | 0.668006436361909 | 0.741 | 0.39 | 0.918698558929805 | Cluster 6 | Akr1a1 |
| Plin3 | 3.15722068586227e-05 | 0.71158041263324 | 0.37 | 0.119 | 0.980411739580811 | Cluster 6 | Plin3 |
| Hk3 | 3.2218262327124e-05 | 0.671641265919402 | 0.556 | 0.2 | 1 | Cluster 6 | Hk3 |
| F7 | 3.31802056642223e-05 | 0.298181914579607 | 0.185 | 0.028 | 1 | Cluster 6 | F7 |
| Atp6v1f1 | 3.37324133320555e-05 | 0.791465559205532 | 0.852 | 0.683 | 1 | Cluster 6 | Atp6v1f |
| Atp6v1g1 | 3.99844022114307e-05 | 0.584234477464773 | 0.889 | 0.799 | 1 | Cluster 6 | Atp6v1g1 |
| Haus8 | 4.09968126331303e-05 | 0.849473063921565 | 0.444 | 0.23 | 1 | Cluster 6 | Haus8 |
| Fam221a | 4.19304259103108e-05 | 0.453515946144409 | 0.111 | 0.008 | 1 | Cluster 6 | Fam221a |
| Adcy3 | 4.19304259103108e-05 | 0.321364333391908 | 0.111 | 0.006 | 1 | Cluster 6 | Adcy3 |
| Ubox5 | 4.19304259103108e-05 | 0.300317984829272 | 0.111 | 0.004 | 1 | Cluster 6 | Ubox5 |
| Proscos | 4.24042814844813e-05 | 0.443710786661708 | 0.148 | 0.017 | 1 | Cluster 6 | Proscos |
| Gm47428 | 4.24042814844813e-05 | 0.267368917463416 | 0.148 | 0.015 | 1 | Cluster 6 | Gm47428 |
| Glmp | 4.31412584421906e-05 | 0.747805660004329 | 0.481 | 0.176 | 1 | Cluster 6 | Glmp |
| Ostm1 | 4.67562202245788e-05 | 0.790197462744918 | 0.444 | 0.115 | 1 | Cluster 6 | Ostm1 |
| Ccl33 | 4.84016821526384e-05 | 0.860040060378486 | 1 | 0.836 | 1 | Cluster 6 | Ccl3 |
| Abr1 | 5.24403528222918e-05 | 0.791426359398147 | 0.741 | 0.413 | 1 | Cluster 6 | Abr |
| Mcfd2 | 5.30216722689466e-05 | 1.14932705038824 | 0.481 | 0.198 | 1 | Cluster 6 | Mcfd2 |
| Iqsec11 | 5.44311475739013e-05 | 0.632964209553612 | 0.704 | 0.34 | 1 | Cluster 6 | Iqsec1 |
| Commd8 | 5.44321686879879e-05 | 0.479724050978559 | 0.481 | 0.134 | 1 | Cluster 6 | Commd8 |
| Tbc1d2 | 6.01073501580934e-05 | 0.363881541745211 | 0.222 | 0.025 | 1 | Cluster 6 | Tbc1d2 |
| Dock10 | 6.37038604935776e-05 | 0.444140299324246 | 0.296 | 0.078 | 1 | Cluster 6 | Dock10 |
| Rragd | 6.55960192439677e-05 | 0.434126717432262 | 0.259 | 0.017 | 1 | Cluster 6 | Rragd |
| Ostf1 | 6.80803327061806e-05 | 0.671540682542812 | 0.889 | 0.903 | 1 | Cluster 6 | Ostf1 |
| Pnpla7 | 6.92189230544723e-05 | 0.955973630708176 | 0.593 | 0.233 | 1 | Cluster 6 | Pnpla7 |
| BC005537 | 7.29437828904907e-05 | 0.437141016140439 | 0.815 | 0.632 | 1 | Cluster 6 | BC005537 |
| C330007P06Rik | 7.29976957583548e-05 | 0.461616829656919 | 0.37 | 0.096 | 1 | Cluster 6 | C330007P06Rik |
| Basp12 | 7.6891507270699e-05 | 0.752559719412992 | 0.926 | 0.784 | 1 | Cluster 6 | Basp1 |
| Cd531 | 8.19627216723122e-05 | 0.565301076829112 | 0.963 | 0.922 | 1 | Cluster 6 | Cd53 |
| Myo5a | 8.48480666244544e-05 | 0.613615768628676 | 0.333 | 0.081 | 1 | Cluster 6 | Myo5a |
| Gaa | 9.21348976567843e-05 | 0.289177410070696 | 0.222 | 0.05 | 1 | Cluster 6 | Gaa |
| Ctsz1 | 9.27249399981569e-05 | 0.592417238313558 | 0.963 | 0.784 | 1 | Cluster 6 | Ctsz |
| Cln3 | 9.35975070451216e-05 | 0.582800601592654 | 0.63 | 0.213 | 1 | Cluster 6 | Cln3 |
| Rell1 | 0.000105056745967754 | 0.652962009863706 | 0.481 | 0.163 | 1 | Cluster 6 | Rell1 |
| Pomk | 0.000108446815024639 | 0.535824899818671 | 0.185 | 0.009 | 1 | Cluster 6 | Pomk |
| Ncapg2 | 0.000112468644339174 | 0.699064441373948 | 0.333 | 0.064 | 1 | Cluster 6 | Ncapg2 |
| A930007I19Rik | 0.000122914658849363 | 0.647202756891152 | 0.259 | 0.037 | 1 | Cluster 6 | A930007I19Rik |
| Mcoln1 | 0.000123685563960616 | 0.546465538944553 | 0.296 | 0.049 | 1 | Cluster 6 | Mcoln1 |
| Nip7 | 0.000136314404696593 | 0.445882009698383 | 0.296 | 0.066 | 1 | Cluster 6 | Nip7 |
| 4933432I03Rik | 0.000136688346108137 | 0.457725533425441 | 0.259 | 0.052 | 1 | Cluster 6 | 4933432I03Rik |
| Vps37b3 | 0.000137692132358598 | 0.628029770755482 | 0.852 | 0.467 | 1 | Cluster 6 | Vps37b |
| Zc3h12c | 0.000143033842074914 | 0.560956399199892 | 0.185 | 0.036 | 1 | Cluster 6 | Zc3h12c |
| Klhdc41 | 0.000143802464501986 | 0.893179102558011 | 0.444 | 0.163 | 1 | Cluster 6 | Klhdc4 |
| Usb1 | 0.000148944123493407 | 0.349079897706626 | 0.222 | 0.072 | 1 | Cluster 6 | Usb1 |
| Trappc3 | 0.000151468478243637 | 0.47247246233526 | 0.407 | 0.12 | 1 | Cluster 6 | Trappc3 |
| Tmem2511 | 0.000152688234268455 | 0.645024431219524 | 0.593 | 0.253 | 1 | Cluster 6 | Tmem251 |
| Mfsd5 | 0.000155049698561508 | 0.410994380432143 | 0.296 | 0.082 | 1 | Cluster 6 | Mfsd5 |
| Lhfpl2 | 0.000157324492717942 | 0.895032717593238 | 0.259 | 0.066 | 1 | Cluster 6 | Lhfpl2 |
| Keap1 | 0.000168709979115841 | 0.362452798060083 | 0.259 | 0.033 | 1 | Cluster 6 | Keap1 |
| Fkbp15 | 0.000169196760230895 | 0.552914454043582 | 0.407 | 0.184 | 1 | Cluster 6 | Fkbp15 |
| Dnmt1 | 0.000188221403196873 | 0.462920674891873 | 0.222 | 0.032 | 1 | Cluster 6 | Dnmt1 |
| Wdr81 | 0.000188221403196873 | 0.338704715741778 | 0.222 | 0.055 | 1 | Cluster 6 | Wdr81 |
| 1810058I24Rik | 0.000198773626636629 | 0.564963267918721 | 0.852 | 0.609 | 1 | Cluster 6 | 1810058I24Rik |
| Mrpl46 | 0.000219608765376874 | 0.397743101229684 | 0.222 | 0.028 | 1 | Cluster 6 | Mrpl46 |
| Hdac4 | 0.000226787232913883 | 0.814067126919791 | 0.444 | 0.131 | 1 | Cluster 6 | Hdac4 |
| Elp5 | 0.000228190219521379 | 0.28807665153168 | 0.222 | 0.052 | 1 | Cluster 6 | Elp5 |
| Sdcbp1 | 0.000241589373964887 | 0.581517301771723 | 1 | 0.904 | 1 | Cluster 6 | Sdcbp |
| Derl11 | 0.000246484681160138 | 0.611328901318435 | 0.778 | 0.422 | 1 | Cluster 6 | Derl1 |
| Gadd45g | 0.000248371081163063 | 1.5267744228359 | 0.444 | 0.176 | 1 | Cluster 6 | Gadd45g |
| Igf2r1 | 0.0002548080394644 | 0.491653999321512 | 0.444 | 0.194 | 1 | Cluster 6 | Igf2r |
| Hmox21 | 0.000271872093189925 | 0.545021063568664 | 0.444 | 0.164 | 1 | Cluster 6 | Hmox2 |
| Cuta2 | 0.000275280300079363 | 0.460766893856623 | 0.519 | 0.23 | 1 | Cluster 6 | Cuta |

| | | | | | | | | |
|---|---|---|---|---|---|---|---|---|
| Heatr1 | 0.000297669267929443 | 0.287234490534853 | 0.222 | 0.049 | 1 | | Cluster 6 | Heatr1 |
| Lipa | 0.00029882578143225 | 0.811251501230818 | 0.593 | 0.263 | 1 | | Cluster 6 | Lipa |
| Stx3 | 0.000301097708388892 | 0.923282265002473 | 0.296 | 0.073 | 1 | | Cluster 6 | Stx3 |
| Kpna41 | 0.000317809719454539 | 0.505494171315161 | 0.778 | 0.595 | 1 | | Cluster 6 | Kpna4 |
| Ctns | 0.000320311989718041 | 0.39007396272757 | 0.148 | 0.018 | 1 | | Cluster 6 | Ctns |
| Eif3a1 | 0.000349058563423631 | 0.596463322361585 | 0.778 | 0.487 | 1 | | Cluster 6 | Eif3a |
| Vps26a | 0.000353608349207906 | 0.60903616971211 | 0.556 | 0.248 | 1 | | Cluster 6 | Vps26a |
| Gstt2 | 0.000353905542915825 | 0.489353650843812 | 0.148 | 0.01 | 1 | | Cluster 6 | Gstt2 |
| Psenen1 | 0.000366206137442474 | 0.53511894377034 | 0.852 | 0.61 | 1 | | Cluster 6 | Psenen |
| Grn | 0.000368483419343298 | 0.499083771578103 | 0.741 | 0.493 | 1 | | Cluster 6 | Grn |
| Txn12 | 0.000376313307873274 | 1.30370412915557 | 0.852 | 0.869 | 1 | | Cluster 6 | Txn1 |
| Cd2741 | 0.000383680846943696 | 0.939844952560134 | 0.667 | 0.358 | 1 | | Cluster 6 | Cd274 |
| Echdc3 | 0.000390772138476534 | 0.499756385149455 | 0.148 | 0.012 | 1 | | Cluster 6 | Echdc3 |
| Lrrc27 | 0.000390772138476534 | 0.282445609946695 | 0.148 | 0.017 | 1 | | Cluster 6 | Lrrc27 |
| AC146911.1 | 0.000390772138476534 | 0.260296952111518 | 0.148 | 0.019 | 1 | | Cluster 6 | AC146911.1 |
| Stab1 | 0.00039825032102975 | 0.387949559384037 | 0.37 | 0.138 | 1 | | Cluster 6 | Stab1 |
| Cep162 | 0.000410523034820894 | 0.34477461056307 | 0.148 | 0.009 | 1 | | Cluster 6 | Cep162 |
| 2810004N23Rik | 0.00041340702154095 | 0.276469810088218 | 0.185 | 0.057 | 1 | | Cluster 6 | 2810004N23Rik |
| Arid5b | 0.000427743736039048 | 0.530923275030145 | 0.667 | 0.337 | 1 | | Cluster 6 | Arid5b |
| Sgpl1 | 0.000430294952356112 | 0.76730965779581 | 0.667 | 0.366 | 1 | | Cluster 6 | Sgpl1 |
| Slc38a10 | 0.000431089734738975 | 0.450579803432453 | 0.333 | 0.102 | 1 | | Cluster 6 | Slc38a10 |
| Nek6 | 0.000431581958906438 | 0.510425885352564 | 0.519 | 0.203 | 1 | | Cluster 6 | Nek6 |
| Eif4a1 | 0.00043579485353987 | 0.494241039704928 | 0.963 | 0.764 | 1 | | Cluster 6 | Eif4a1 |
| Selenok | 0.000452426811474331 | 0.361645353394697 | 0.963 | 0.751 | 1 | | Cluster 6 | Selenok |
| Sntb2 | 0.000457059447106094 | 0.367789487267264 | 0.556 | 0.258 | 1 | | Cluster 6 | Sntb2 |
| Tollip | 0.000459243812923349 | 0.340102346350341 | 0.37 | 0.163 | 1 | | Cluster 6 | Tollip |
| Dnase1l1 | 0.000463187967039952 | 0.360234155343536 | 0.259 | 0.081 | 1 | | Cluster 6 | Dnase1l1 |
| Slc38a11 | 0.000466510593541392 | 0.872204428207268 | 0.667 | 0.363 | 1 | | Cluster 6 | Slc38a1 |
| Topors | 0.0005051497410946 | 0.385367335865552 | 0.222 | 0.06 | 1 | | Cluster 6 | Topors |
| Sh3bp2 | 0.000526264734214632 | 0.602149041289083 | 0.333 | 0.103 | 1 | | Cluster 6 | Sh3bp2 |
| Rab18 | 0.000534225944985041 | 0.388417921033804 | 0.37 | 0.157 | 1 | | Cluster 6 | Rab18 |
| Tob1 | 0.000557583833335827 | 0.321417533138541 | 0.481 | 0.202 | 1 | | Cluster 6 | Tob1 |
| Cog1 | 0.000600916637887715 | 0.329024204132658 | 0.222 | 0.038 | 1 | | Cluster 6 | Cog1 |
| Carnmt11 | 0.000614524609443843 | 0.485472340967553 | 0.333 | 0.115 | 1 | | Cluster 6 | Carnmt1 |
| Ccr2 | 0.00062199301418782 | 0.277418666220699 | 0.222 | 0.05 | 1 | | Cluster 6 | Ccr2 |
| Thbs12 | 0.000637326067655861 | 0.619080886558736 | 0.889 | 0.623 | 1 | | Cluster 6 | Thbs1 |
| Dpp7 | 0.000641382179835608 | 0.262987867874603 | 0.185 | 0.039 | 1 | | Cluster 6 | Dpp7 |
| Tmem65 | 0.000651207664952183 | 0.396365585669753 | 0.296 | 0.083 | 1 | | Cluster 6 | Tmem65 |
| Plekhm2 | 0.00065164421716056 | 0.44362166890165 | 0.407 | 0.153 | 1 | | Cluster 6 | Plekhm2 |
| Gm20406 | 0.000652011432789594 | 0.55437022238629 | 0.556 | 0.261 | 1 | | Cluster 6 | Gm20406 |
| Arl11 | 0.00065404400723359 | 0.913310434450368 | 0.407 | 0.149 | 1 | | Cluster 6 | Arl11 |
| Sbno1 | 0.0006941557818426 | 0.418131733089068 | 0.926 | 0.566 | 1 | | Cluster 6 | Sbno1 |
| Tra2a | 0.000722840396507682 | 0.667488530465714 | 0.889 | 0.655 | 1 | | Cluster 6 | Tra2a |
| Prrg1 | 0.000758068654811029 | 0.558643815254885 | 0.111 | 0.008 | 1 | | Cluster 6 | Prrg1 |
| Vwf | 0.000758068654811029 | 0.464357577986049 | 0.111 | 0.004 | 1 | | Cluster 6 | Vwf |
| Mkrn1 | 0.00077640739567293 | 0.399668586407471 | 0.704 | 0.434 | 1 | | Cluster 6 | Mkrn1 |
| Tbc1d10a | 0.000801971274613394 | 0.453650119079979 | 0.37 | 0.088 | 1 | | Cluster 6 | Tbc1d10a |
| Rab15 | 0.000802821978974923 | 0.585448264465611 | 0.111 | 0.011 | 1 | | Cluster 6 | Rab15 |
| Vcam1 | 0.000802821978974923 | 0.506198554289054 | 0.111 | 0.007 | 1 | | Cluster 6 | Vcam1 |
| 9130221H12Rik | 0.000850016780507448 | 0.288832167617637 | 0.111 | 0.012 | 1 | | Cluster 6 | 9130221H12Rik |
| Cpeb4 | 0.000850327698039888 | 0.824979007884798 | 0.519 | 0.332 | 1 | | Cluster 6 | Cpeb4 |
| Ubc1 | 0.000851928240512699 | 0.411735315017712 | 1 | 0.986 | 1 | | Cluster 6 | Ubc |
| Mpeg11 | 0.000860193790336122 | 0.544106256414135 | 0.741 | 0.562 | 1 | | Cluster 6 | Mpeg1 |
| Rab5a | 0.000866257725909438 | 0.483481025768341 | 0.519 | 0.262 | 1 | | Cluster 6 | Rab5a |
| Ptpn23 | 0.000890787549679695 | 0.432914646102195 | 0.407 | 0.136 | 1 | | Cluster 6 | Ptpn23 |
| Ccl41 | 0.000908290329344272 | 0.731937329912605 | 0.852 | 0.618 | 1 | | Cluster 6 | Ccl4 |
| Ccrl23 | 0.000919301781136797 | 0.67995822488306 | 0.963 | 0.898 | 1 | | Cluster 6 | Ccrl2 |
| Ube2q11 | 0.000926093017966769 | 0.251590908905952 | 0.481 | 0.25 | 1 | | Cluster 6 | Ube2q1 |
| Gon7 | 0.000969075112841683 | 0.318487616708083 | 0.296 | 0.101 | 1 | | Cluster 6 | Gon7 |
| Zfr | 0.00101631934008994 | 0.467554050981116 | 0.407 | 0.138 | 1 | | Cluster 6 | Zfr |
| Cbx4 | 0.00104884246308376 | 0.423792057105526 | 0.296 | 0.093 | 1 | | Cluster 6 | Cbx4 |

| | | | | | | | | |
|---|---|---|---|---|---|---|---|---|
| Plekhm1 | 0.00104973875529537 | 0.411153354017035 | 0.259 | 0.051 | 1 | | Cluster 6 | Plekhm1 |
| Jtb | 0.0011666788990035 | 0.543186188834032 | 0.37 | 0.152 | 1 | | Cluster 6 | Jtb |
| Fam50a1 | 0.00122195431776426 | 0.519904047197808 | 0.519 | 0.254 | 1 | | Cluster 6 | Fam50a |
| Amz21 | 0.00129220419184787 | 0.296456530569949 | 0.296 | 0.119 | 1 | | Cluster 6 | Amz2 |
| Atp6ap22 | 0.00135802353578574 | 0.550311348312478 | 0.778 | 0.608 | 1 | | Cluster 6 | Atp6ap2 |
| Snx4 | 0.00138974794923151 | 0.366160625701876 | 0.37 | 0.184 | 1 | | Cluster 6 | Snx4 |
| Plbd11 | 0.00144419182819717 | 1.08949929932449 | 0.667 | 0.488 | 1 | | Cluster 6 | Plbd1 |
| Dvl2 | 0.00149186056740354 | 0.341677629927178 | 0.185 | 0.053 | 1 | | Cluster 6 | Dvl2 |
| Crkl | 0.00149186056740354 | 0.286131945399925 | 0.185 | 0.054 | 1 | | Cluster 6 | Crkl |
| Wdfy3 | 0.00157616811213329 | 0.44482364844299 | 0.296 | 0.137 | 1 | | Cluster 6 | Wdfy3 |
| Aspscr1 | 0.00159753645377124 | 0.401488628943246 | 0.259 | 0.08 | 1 | | Cluster 6 | Aspscr1 |
| Mospd1 | 0.00161254707428993 | 0.497233323453294 | 0.148 | 0.022 | 1 | | Cluster 6 | Mospd1 |
| Il10rb | 0.00162628730077474 | 0.470252868491047 | 0.926 | 0.676 | 1 | | Cluster 6 | Il10rb |
| Slc6a6 | 0.00166449988071514 | 0.525359156344964 | 0.815 | 0.545 | 1 | | Cluster 6 | Slc6a6 |
| Fbxo42 | 0.00175126790218753 | 0.278047117462282 | 0.148 | 0.032 | 1 | | Cluster 6 | Fbxo42 |
| Kdsr | 0.00177636590910841 | 0.295227503316593 | 0.185 | 0.049 | 1 | | Cluster 6 | Kdsr |
| Gm43796 | 0.001824668186479 | 0.284785262610182 | 0.148 | 0.015 | 1 | | Cluster 6 | Gm43796 |
| Ubb1 | 0.00183924467213701 | 0.499330086230927 | 1 | 0.982 | 1 | | Cluster 6 | Ubb |
| Cycs2 | 0.00184359917246608 | 0.586737625137023 | 0.519 | 0.337 | 1 | | Cluster 6 | Cycs |
| Eya3 | 0.00188399808811466 | 0.660585379944545 | 0.222 | 0.037 | 1 | | Cluster 6 | Eya3 |
| Washc2 | 0.00190064167124702 | 0.426613936485703 | 0.63 | 0.311 | 1 | | Cluster 6 | Washc2 |
| Fam46c | 0.00197380211386001 | 0.478203322904991 | 0.222 | 0.041 | 1 | | Cluster 6 | Fam46c |
| Sirt2 | 0.00200178359271215 | 0.415523055808371 | 0.519 | 0.234 | 1 | | Cluster 6 | Sirt2 |
| Wdr261 | 0.00204966053611225 | 0.442627321142767 | 0.704 | 0.39 | 1 | | Cluster 6 | Wdr26 |
| Retreg2 | 0.0020946223728596 | 0.327683871055416 | 0.444 | 0.205 | 1 | | Cluster 6 | Retreg2 |
| Zc3h4 | 0.00209724323253713 | 0.252720884207449 | 0.222 | 0.05 | 1 | | Cluster 6 | Zc3h4 |
| Ttc32 | 0.00219742452764554 | 0.391571606213901 | 0.333 | 0.098 | 1 | | Cluster 6 | Ttc32 |
| Tm9sf4 | 0.00223151038655342 | 0.478376162843879 | 0.296 | 0.107 | 1 | | Cluster 6 | Tm9sf4 |
| B2m1 | 0.00228670649639399 | 0.361876055324566 | 1 | 0.952 | 1 | | Cluster 6 | B2m |
| Tbc1d23 | 0.00230614471855707 | 0.365046640052124 | 0.407 | 0.198 | 1 | | Cluster 6 | Tbc1d23 |
| Eif51 | 0.00242411274842781 | 0.685613114076008 | 0.926 | 0.816 | 1 | | Cluster 6 | Eif5 |
| Slu7 | 0.00242789879379569 | 0.420629730866134 | 0.37 | 0.19 | 1 | | Cluster 6 | Slu7 |
| Vav1 | 0.00243117421386545 | 0.288241264124001 | 0.704 | 0.411 | 1 | | Cluster 6 | Vav1 |
| Prdx12 | 0.00245591630479007 | 0.941980788682162 | 0.667 | 0.433 | 1 | | Cluster 6 | Prdx1 |
| Dnajb91 | 0.00248809418931862 | 1.05944164043831 | 0.519 | 0.286 | 1 | | Cluster 6 | Dnajb9 |
| 5031425E22Rik | 0.00249291954392429 | 0.431927412022267 | 0.593 | 0.287 | 1 | | Cluster 6 | 5031425E22Rik |
| Ykt6 | 0.00250906425771311 | 0.306442670034822 | 0.296 | 0.111 | 1 | | Cluster 6 | Ykt6 |
| Fbxo38 | 0.00251160129666723 | 0.360797308715267 | 0.259 | 0.084 | 1 | | Cluster 6 | Fbxo38 |
| Gadd45b1 | 0.00258485744072348 | 0.385434189657058 | 1 | 0.923 | 1 | | Cluster 6 | Gadd45b |
| Slc25a331 | 0.00264004499859727 | 0.452174647587115 | 0.481 | 0.244 | 1 | | Cluster 6 | Slc25a33 |
| Plk2 | 0.00273986697177304 | 1.17486758976054 | 0.259 | 0.097 | 1 | | Cluster 6 | Plk2 |
| Myo9b | 0.00277982430844577 | 0.417617423270391 | 0.593 | 0.281 | 1 | | Cluster 6 | Myo9b |
| Npc21 | 0.00278811643379562 | 0.486729927580466 | 0.704 | 0.475 | 1 | | Cluster 6 | Npc2 |
| Washc5 | 0.00280412173617718 | 0.299040032497327 | 0.185 | 0.058 | 1 | | Cluster 6 | Washc5 |
| Wwp2 | 0.00283480862846424 | 0.411719421117598 | 0.407 | 0.21 | 1 | | Cluster 6 | Wwp2 |
| Atp6v1e1 | 0.00284230627634412 | 0.6362241522964 | 0.852 | 0.686 | 1 | | Cluster 6 | Atp6v1e1 |
| Dnttip1 | 0.00288957782932585 | 0.372013094813531 | 0.296 | 0.119 | 1 | | Cluster 6 | Dnttip1 |
| Zfand5 | 0.00314776159623572 | 0.651075875643254 | 0.741 | 0.46 | 1 | | Cluster 6 | Zfand5 |
| Clec4d4 | 0.00319492906631817 | 0.388238934820825 | 0.963 | 0.939 | 1 | | Cluster 6 | Clec4d |
| Nsmce2 | 0.0032801861800386 | 0.319694583293599 | 0.37 | 0.165 | 1 | | Cluster 6 | Nsmce2 |
| Nampt1 | 0.00333888776568181 | 0.336993857322879 | 0.481 | 0.187 | 1 | | Cluster 6 | Nampt |
| U2surp1 | 0.00334757346927642 | 0.582068498690716 | 0.481 | 0.229 | 1 | | Cluster 6 | U2surp |
| Fcgr2b1 | 0.00335825487079086 | 0.515994298850832 | 0.407 | 0.175 | 1 | | Cluster 6 | Fcgr2b |
| Purb1 | 0.00337379014555182 | 0.323319385907066 | 0.778 | 0.493 | 1 | | Cluster 6 | Purb |
| Rgs1 | 0.00342075072430875 | 1.11821306844269 | 0.481 | 0.194 | 1 | | Cluster 6 | Rgs1 |
| Rhov2 | 0.00344930842999267 | 0.254654521088867 | 0.37 | 0.171 | 1 | | Cluster 6 | Rhov |
| Hmg20b | 0.00349111498242401 | 0.319129809476239 | 0.259 | 0.107 | 1 | | Cluster 6 | Hmg20b |
| 2900093K20Rik | 0.00349827621556117 | 0.307851263630454 | 0.185 | 0.052 | 1 | | Cluster 6 | 2900093K20Rik |
| Slc31a1 | 0.00350600988789288 | 0.318661808699615 | 0.37 | 0.163 | 1 | | Cluster 6 | Slc31a1 |
| Tfeb | 0.00354072597009343 | 0.44984173489749 | 0.37 | 0.098 | 1 | | Cluster 6 | Tfeb |
| Kmt5a | 0.0035830442668255 | 0.421734167636142 | 0.481 | 0.246 | 1 | | Cluster 6 | Kmt5a |

| | | | | | | | | |
|---|---|---|---|---|---|---|---|---|
| Csf2rb2 | 0.00366837539167818 | 0.679168184349578 | 0.407 | 0.177 | 1 | | Cluster 6 | Csf2rb2 |
| Pkig | 0.00374707620857008 | 0.315102279342358 | 0.222 | 0.048 | 1 | | Cluster 6 | Pkig |
| Frrs11 | 0.00379381332893723 | 0.486017409601102 | 0.519 | 0.242 | 1 | | Cluster 6 | Frrs1 |
| Trim8 | 0.00383658268255038 | 0.51341597415929 | 0.444 | 0.185 | 1 | | Cluster 6 | Trim8 |
| Pigyl | 0.0038410256839532 | 0.28097604999254 | 0.185 | 0.048 | 1 | | Cluster 6 | Pigyl |
| Zmiz11 | 0.00384566936121019 | 0.472341032270965 | 0.593 | 0.368 | 1 | | Cluster 6 | Zmiz1 |
| Clcn41 | 0.00391659133716683 | 0.560861703984535 | 0.444 | 0.169 | 1 | | Cluster 6 | Clcn4 |
| Eea1 | 0.00393367170719031 | 0.719061742962017 | 0.593 | 0.314 | 1 | | Cluster 6 | Eea1 |
| Tcirg12 | 0.00396803814316432 | 0.727848408163172 | 0.556 | 0.332 | 1 | | Cluster 6 | Tcirg1 |
| Pip4k2c | 0.00405399372893031 | 0.386061559307306 | 0.37 | 0.116 | 1 | | Cluster 6 | Pip4k2c |
| Acot13 | 0.00410127297051318 | 0.275350417990845 | 0.259 | 0.093 | 1 | | Cluster 6 | Acot13 |
| Rybp | 0.0041261117486967 | 0.727593216695095 | 0.593 | 0.306 | 1 | | Cluster 6 | Rybp |
| Strn | 0.00415207867113568 | 0.340498695629831 | 0.259 | 0.086 | 1 | | Cluster 6 | Strn |
| Etf1 | 0.00424123721211677 | 0.317422689795364 | 0.926 | 0.722 | 1 | | Cluster 6 | Etf1 |
| Zfp655 | 0.00429612324039083 | 0.585202282933213 | 0.296 | 0.089 | 1 | | Cluster 6 | Zfp655 |
| Tmem120a | 0.00432800833541538 | 0.284959461858012 | 0.333 | 0.144 | 1 | | Cluster 6 | Tmem120a |
| Gna131 | 0.00445979824437368 | 0.572045751983226 | 0.704 | 0.537 | 1 | | Cluster 6 | Gna13 |
| Tnni2 | 0.00448241396301299 | 0.786937050823594 | 0.296 | 0.099 | 1 | | Cluster 6 | Tnni2 |
| S1pr2 | 0.00451991700966174 | 0.310418304059515 | 0.148 | 0.034 | 1 | | Cluster 6 | S1pr2 |
| Atp6v0d2 | 0.00456925234732687 | 0.643280302513534 | 0.111 | 0.012 | 1 | | Cluster 6 | Atp6v0d2 |
| Comtd1 | 0.00465842877003152 | 0.274052071948398 | 0.296 | 0.137 | 1 | | Cluster 6 | Comtd1 |
| Napa1 | 0.00473214395634142 | 0.384741305203157 | 0.63 | 0.313 | 1 | | Cluster 6 | Napa |
| Sh2b1 | 0.00476241425681014 | 0.271250459534958 | 0.185 | 0.03 | 1 | | Cluster 6 | Sh2b1 |
| Cdc5l1 | 0.00478383524652619 | 0.564268573185928 | 0.444 | 0.248 | 1 | | Cluster 6 | Cdc5l |
| 4833418N02Rik | 0.00490924301304368 | 0.250778242370499 | 0.185 | 0.027 | 1 | | Cluster 6 | 4833418N02Rik |
| Hnrnpm2 | 0.00505748164163164 | 0.454105580639133 | 0.667 | 0.381 | 1 | | Cluster 6 | Hnrnpm |
| Exosc4 | 0.00517581668143081 | 0.262268132672455 | 0.222 | 0.082 | 1 | | Cluster 6 | Exosc4 |
| S100a1 | 0.00520515698032571 | 0.588206800413151 | 0.148 | 0.056 | 1 | | Cluster 6 | S100a1 |
| Snrnp35 | 0.00520515698032571 | 0.268937193772177 | 0.148 | 0.035 | 1 | | Cluster 6 | Snrnp35 |
| Dhfr | 0.00521758622012796 | 0.304842905293771 | 0.111 | 0.014 | 1 | | Cluster 6 | Dhfr |
| Lamtor31 | 0.00522084791419361 | 0.341455112887341 | 0.556 | 0.311 | 1 | | Cluster 6 | Lamtor3 |
| Nbr1 | 0.00526277223544877 | 0.577675739893185 | 0.259 | 0.093 | 1 | | Cluster 6 | Nbr1 |
| Arl8a | 0.0052810117532234 | 0.339568917426081 | 0.593 | 0.283 | 1 | | Cluster 6 | Arl8a |
| Tgoln11 | 0.00530336673514483 | 0.523156157935647 | 0.852 | 0.764 | 1 | | Cluster 6 | Tgoln1 |
| Map3k8 | 0.00540224696336241 | 0.401666210594351 | 0.556 | 0.327 | 1 | | Cluster 6 | Map3k8 |
| Ttc1 | 0.00541124196625451 | 0.364989090909256 | 0.333 | 0.139 | 1 | | Cluster 6 | Ttc1 |
| Snx291 | 0.00546923483719018 | 0.539545954884206 | 0.333 | 0.105 | 1 | | Cluster 6 | Snx29 |
| Tmem86a | 0.0055818295711603 | 0.254791094132399 | 0.148 | 0.039 | 1 | | Cluster 6 | Tmem86a |
| Rsrp1 | 0.00559171869880347 | 0.702791507309588 | 0.741 | 0.546 | 1 | | Cluster 6 | Rsrp1 |
| Wbp21 | 0.00582245492842416 | 0.337603432746423 | 0.519 | 0.308 | 1 | | Cluster 6 | Wbp2 |
| Cd300lb | 0.0059802281352362 | 0.336297669700934 | 0.333 | 0.152 | 1 | | Cluster 6 | Cd300lb |
| Foxn3 | 0.00606425082770107 | 0.36211348807097 | 0.296 | 0.123 | 1 | | Cluster 6 | Foxn3 |
| Asna1 | 0.00606425082770107 | 0.320288368020085 | 0.296 | 0.115 | 1 | | Cluster 6 | Asna1 |
| H2-D14 | 0.00607350029364129 | 0.551060600179481 | 1 | 0.959 | 1 | | Cluster 6 | H2-D1 |
| Esd1 | 0.00609164860513995 | 0.919329780925971 | 0.593 | 0.397 | 1 | | Cluster 6 | Esd |
| Lmbrd1 | 0.00612007616481797 | 0.511526138073082 | 0.407 | 0.179 | 1 | | Cluster 6 | Lmbrd1 |
| Rnf341 | 0.00615374752447767 | 0.321406053649138 | 0.333 | 0.136 | 1 | | Cluster 6 | Rnf34 |
| Scamp31 | 0.00666774386654758 | 0.291929800333354 | 0.333 | 0.167 | 1 | | Cluster 6 | Scamp3 |
| Washc4 | 0.00667952473754297 | 0.333073505545649 | 0.407 | 0.192 | 1 | | Cluster 6 | Washc4 |
| Smim14 | 0.00673201758546516 | 0.580044935167846 | 0.519 | 0.317 | 1 | | Cluster 6 | Smim14 |
| Tbc1d15 | 0.0068024805827578 | 0.530575150596692 | 0.481 | 0.228 | 1 | | Cluster 6 | Tbc1d15 |
| Map1lc3b2 | 0.00726823056358954 | 0.427904336992755 | 0.889 | 0.841 | 1 | | Cluster 6 | Map1lc3b |
| Eif4g11 | 0.0072951802246408 | 0.294421758932461 | 0.444 | 0.214 | 1 | | Cluster 6 | Eif4g1 |
| Ppp4r21 | 0.00738419243403812 | 0.540671434309873 | 0.63 | 0.397 | 1 | | Cluster 6 | Ppp4r2 |
| Ifit1bl21 | 0.00752048737517424 | 0.416448093798183 | 0.185 | 0.069 | 1 | | Cluster 6 | Ifit1bl2 |
| Taldo12 | 0.00753930349538132 | 0.815464099261 | 0.778 | 0.689 | 1 | | Cluster 6 | Taldo1 |
| Elmo11 | 0.0076495196034324 | 0.27100003665741 | 0.481 | 0.3 | 1 | | Cluster 6 | Elmo1 |
| Spty2d1 | 0.00773452242896875 | 0.363809468181382 | 0.593 | 0.317 | 1 | | Cluster 6 | Spty2d1 |
| Zbtb43 | 0.00779704390410921 | 0.323216725761669 | 0.222 | 0.061 | 1 | | Cluster 6 | Zbtb43 |
| Ubn11 | 0.00784551990224165 | 0.520526820216257 | 0.63 | 0.343 | 1 | | Cluster 6 | Ubn1 |
| Gpatch8 | 0.00795871012115318 | 0.29886569368634 | 0.259 | 0.092 | 1 | | Cluster 6 | Gpatch8 |

| | | | | | | | |
|---|---|---|---|---|---|---|---|
| Ubap1 | 0.00796543381800024 | 0.59084375907921 | 0.444 | 0.189 | 1 | Cluster 6 | Ubap1 |
| Lgmn | 0.00803385052816583 | 0.43987709811799 | 0.519 | 0.345 | 1 | Cluster 6 | Lgmn |
| St132 | 0.00808563392157539 | 0.486145449003744 | 0.704 | 0.356 | 1 | Cluster 6 | St13 |
| Acin1 | 0.00816321103415473 | 0.357052416152741 | 0.444 | 0.253 | 1 | Cluster 6 | Acin1 |
| Azin1 | 0.00821708678691963 | 0.419382388875516 | 0.704 | 0.377 | 1 | Cluster 6 | Azin1 |
| Cd84 | 0.00833031617092558 | 0.435863313547638 | 0.481 | 0.225 | 1 | Cluster 6 | Cd84 |
| Ccdc591 | 0.00834996300204101 | 0.456778714334612 | 0.481 | 0.168 | 1 | Cluster 6 | Ccdc59 |
| Plekhf2 | 0.00842829060172437 | 0.602051107156203 | 0.481 | 0.174 | 1 | Cluster 6 | Plekhf2 |
| Mtm1 | 0.00856720376805795 | 0.422530992346877 | 0.222 | 0.084 | 1 | Cluster 6 | Mtm1 |
| Mgea5 | 0.00863121019301816 | 0.29228448457493 | 0.556 | 0.377 | 1 | Cluster 6 | Mgea5 |
| Vapa | 0.00875139888917107 | 0.286854998347918 | 0.815 | 0.552 | 1 | Cluster 6 | Vapa |
| Ilf3 | 0.00882539237702054 | 0.57782799760145 | 0.556 | 0.192 | 1 | Cluster 6 | Ilf3 |
| Csnk1e | 0.00895416490243223 | 0.3805973251793 | 0.519 | 0.3 | 1 | Cluster 6 | Csnk1e |
| Rcbtb2 | 0.00899055272944 | 0.426681709616122 | 0.37 | 0.179 | 1 | Cluster 6 | Rcbtb2 |
| Trappc2l2 | 0.00900261406530126 | 0.254532600778458 | 0.444 | 0.233 | 1 | Cluster 6 | Trappc2l |
| AW549877 | 0.0091235957959556 | 0.351748465818488 | 0.296 | 0.102 | 1 | Cluster 6 | AW549877 |
| Gpbp1 | 0.00919914658345299 | 0.424984596601242 | 0.778 | 0.493 | 1 | Cluster 6 | Gpbp1 |
| Acsl11 | 0.00953808082884691 | 0.413780117702262 | 0.481 | 0.274 | 1 | Cluster 6 | Acsl1 |
| Vps11 | 0.00954327048903813 | 0.285658448977201 | 0.222 | 0.065 | 1 | Cluster 6 | Vps11 |
| Rbbp6 | 0.00962347548776574 | 0.32264391727632 | 0.556 | 0.258 | 1 | Cluster 6 | Rbbp6 |
| 9030624J02Rik | 0.00984247224475412 | 0.496596813215174 | 0.296 | 0.091 | 1 | Cluster 6 | 9030624J02Rik |
| Chst11 | 0.00990582071464564 | 0.47341785096056 | 0.407 | 0.189 | 1 | Cluster 6 | Chst11 |

#### Marker genes Cluster 1 versus Cluster 3

| Column1 | Column2 | Column3 | Column4 | Column5 | Column6 |
|---|---|---|---|---|---|
| | p_val | avg_logFC | pct.1 | pct.2 | p_val_adj |
| Cxcl2 | 4.62777675365037e-22 | 0.530196528348989 | 0.998 | 0.993 | 1.43706351531105e-17 |
| Ndufa11 | 9.64348877789325e-16 | -0.441653560138767 | 0.112 | 0.352 | 2.99459257019919e-11 |
| Nfkbia | 1.28979821848568e-13 | 0.31629278106156 | 0.994 | 0.962 | 4.00521040786359e-09 |
| Rpl28 | 4.65755611118426e-13 | -0.311964850319171 | 0.84 | 0.955 | 1.44631089920605e-08 |
| Rps23 | 1.08452834952899e-12 | -0.308609982519372 | 0.832 | 0.979 | 3.36778588379238e-08 |
| Cox7a2 | 1.4610904262849e-12 | -0.360721191358581 | 0.759 | 0.934 | 4.5371241007425e-08 |
| Lsm12 | 2.70756344331345e-11 | -0.330771818936794 | 0.201 | 0.449 | 8.40779676052126e-07 |
| Snrpe | 3.56143581351908e-11 | -0.403399426884656 | 0.448 | 0.707 | 1.10593266317208e-06 |
| Rgs10 | 7.08500431949533e-11 | -0.366873626291343 | 0.301 | 0.557 | 2.20010639133288e-06 |
| Rbx1 | 1.5714199401108e-10 | -0.330176335678346 | 0.651 | 0.854 | 4.87973034002608e-06 |
| Txnip | 5.73539948570844e-10 | -0.576968073551431 | 0.566 | 0.746 | 1.78101360229704e-05 |
| Ndufb8 | 9.83185063410171e-10 | -0.34932672538248 | 0.328 | 0.582 | 3.0530845774076e-05 |
| Rpl32 | 1.87665156392319e-09 | -0.261510044887632 | 0.88 | 0.958 | 5.8275661014507e-05 |
| Ndufs6 | 2.87775295602711e-09 | -0.332004203638751 | 0.326 | 0.551 | 8.93628625435099e-05 |
| Ptgs2 | 3.734552979254e-09 | 0.517908765716685 | 0.905 | 0.861 | 0.000115969073664774 |
| Eif3m | 5.100485859041e-09 | -0.312843239222281 | 0.183 | 0.383 | 0.0001583853873808 |
| Sdha | 5.21500250867107e-09 | -0.254003037833731 | 0.183 | 0.394 | 0.000161941472901763 |
| Slc7a11 | 5.30252317773227e-09 | 0.460077432165651 | 0.838 | 0.718 | 0.00016465925223812 |
| Ilf2 | 5.41339123575349e-09 | -0.328597432843082 | 0.12 | 0.293 | 0.000168102038043853 |
| Acadl | 5.64020809018692e-09 | -0.313095650363024 | 0.16 | 0.348 | 0.000175145381824574 |
| Hist1h1c | 5.93228025726665e-09 | -0.385794602384935 | 0.282 | 0.505 | 0.000184215098828901 |
| Snrpd2 | 6.80067576767185e-09 | -0.323094679834689 | 0.299 | 0.526 | 0.000211181384613514 |
| Tnfaip3 | 8.10122262320701e-09 | 0.381638533874465 | 0.923 | 0.875 | 0.000251567266118447 |
| Il1rn | 8.88673022390444e-09 | 0.415756740103177 | 0.963 | 0.92 | 0.000275959633642905 |
| Tnf | 9.66818300848187e-09 | 0.460009827713949 | 0.959 | 0.875 | 0.000300226086962387 |
| Nol7 | 9.89269207698203e-09 | -0.294641155438223 | 0.239 | 0.46 | 0.000307197767066523 |
| Ndufa5 | 1.49147795023942e-08 | -0.26825148397776 | 0.054 | 0.181 | 0.000463148647887846 |
| Rps18 | 1.69672636633984e-08 | -0.278814179858798 | 0.803 | 0.906 | 0.000526884438539509 |
| Rpl26 | 1.78145738758223e-08 | -0.264016770097453 | 0.896 | 0.948 | 0.000553195962565911 |
| mt-Nd2 | 1.78995216054647e-08 | -0.349901064559556 | 0.722 | 0.868 | 0.000555833844414494 |
| Ndufa13 | 1.90722293220365e-08 | -0.255588755622146 | 0.624 | 0.822 | 0.0005922499371372 |
| Mrps14 | 1.93361075575508e-08 | -0.323869907411517 | 0.28 | 0.505 | 0.000600444147984625 |
| Eif3f | 2.31582925260694e-08 | -0.275762587705524 | 0.232 | 0.449 | 0.000719134457812032 |
| Snw1 | 2.35341258521862e-08 | -0.294796244272857 | 0.205 | 0.404 | 0.000730805210087937 |
| Rpl18a | 2.39492487342605e-08 | -0.263212950239446 | 0.921 | 0.969 | 0.000743696020944991 |
| Dek | 2.63349303706596e-08 | -0.305301690913911 | 0.205 | 0.408 | 0.000817778592800092 |
| Acod1 | 2.64649259053288e-08 | 0.512663578513624 | 0.793 | 0.659 | 0.000821815344138174 |
| Nlrp3 | 2.70346794730053e-08 | 0.365290997362772 | 0.902 | 0.861 | 0.000839507901675235 |
| Trf | 3.12109617325718e-08 | -0.318958752378695 | 0.158 | 0.331 | 0.000969193994681553 |
| Ndufb5 | 3.42910738161199e-08 | -0.316592931449286 | 0.274 | 0.484 | 0.00106484071521197 |
| Prkar1a | 3.451002536049e-08 | -0.273127741746819 | 0.479 | 0.732 | 0.0010716398175193 |
| Rpl34 | 3.70149745531494e-08 | -0.253303265283204 | 0.896 | 0.976 | 0.00114942600479895 |
| Srsf3 | 4.08890970395978e-08 | -0.264983874914839 | 0.554 | 0.756 | 0.00126972913037063 |
| Psmc3 | 5.2480457841165e-08 | -0.296234283186724 | 0.191 | 0.38 | 0.0016296756573417 |

| | | | | | |
|---|---|---|---|---|---|
| Ppp1r15a | 5.41156172786163e-08 | 0.336764735100561 | 0.886 | 0.843 | 0.00168045226335287 |
| 1110004F10Rik | 5.66579973065346e-08 | -0.309145372831981 | 0.154 | 0.328 | 0.00175940079035982 |
| Rps24 | 5.88462861012774e-08 | -0.268849791500711 | 0.871 | 0.941 | 0.00182735372230297 |
| Atp5h | 1.02288843939108e-07 | -0.279099845736453 | 0.571 | 0.774 | 0.00317637547084111 |
| Lyl1 | 1.05130582051595e-07 | -0.299393429502934 | 0.108 | 0.254 | 0.00326461996444817 |
| Dnajc3 | 1.10187217193091e-07 | -0.256487208089655 | 0.127 | 0.289 | 0.00342164365549705 |
| Ybx1 | 1.24738932377398e-07 | -0.256899252260115 | 0.695 | 0.875 | 0.00387351806711535 |
| Mrpl30 | 1.31471598688327e-07 | -0.28604478864272 | 0.197 | 0.38 | 0.00408258755406861 |
| Limd2 | 1.31496775449906e-07 | -0.315025716655047 | 0.222 | 0.422 | 0.00408336936804592 |
| Lamtor2 | 1.33162904067912e-07 | -0.301364155482372 | 0.459 | 0.679 | 0.00413510766002088 |
| Slfn2 | 1.42906280755647e-07 | 0.31028381931962 | 0.925 | 0.892 | 0.00443766873630512 |
| Sec11c | 1.51800111276612e-07 | -0.255975452367086 | 0.322 | 0.547 | 0.00471384885547262 |
| Ebna1bp2 | 1.75155425559484e-07 | -0.264303208959647 | 0.197 | 0.39 | 0.00543910142989864 |
| Crlf2 | 1.96282962875001e-07 | -0.290212894230697 | 0.311 | 0.519 | 0.0060951748461574 |
| Cwc15 | 2.59294050956761e-07 | -0.292209419813186 | 0.27 | 0.46 | 0.00805185816436029 |
| Psmb3 | 2.90877270252339e-07 | -0.283554273083556 | 0.521 | 0.756 | 0.00903261187314587 |
| Plek | 2.9723667888306e-07 | 0.271553911444919 | 0.977 | 0.965 | 0.00923009058935568 |
| Etfb | 3.15224545998112e-07 | -0.260607386540141 | 0.195 | 0.376 | 0.00978866782687937 |
| Ube3a | 4.40980398568296e-07 | -0.312589824942789 | 0.16 | 0.314 | 0.0136937643167413 |
| Dnttip2 | 4.47253722340172e-07 | -0.296174256439314 | 0.172 | 0.334 | 0.0138885698398293 |
| Tma7 | 4.7580640652893e-07 | -0.267035036538935 | 0.589 | 0.787 | 0.0147752163419429 |
| Ndufb4 | 5.12535645491033e-07 | -0.254050896612517 | 0.268 | 0.47 | 0.0159157693994331 |
| Rpl15 | 5.83029088712787e-07 | -0.272478286244779 | 0.697 | 0.875 | 0.0181048022917982 |
| Rnf149 | 6.2307222572029e-07 | 0.339451016030104 | 0.859 | 0.791 | 0.0193482618252922 |
| Fuca1 | 6.5369690651919e-07 | -0.274844264554 | 0.257 | 0.449 | 0.0202992500381404 |
| Lsm6 | 6.72236990180926e-07 | -0.253189638029915 | 0.268 | 0.46 | 0.0208749752560883 |
| Hcar2 | 6.99772639224193e-07 | 0.469712878571507 | 0.728 | 0.627 | 0.0217300397658289 |
| Zfp36 | 7.41524584163721e-07 | 0.316202523141746 | 0.934 | 0.899 | 0.023026562912036 |
| Ghitm | 7.44724675730619e-07 | -0.250616017065646 | 0.174 | 0.345 | 0.0231259353554629 |
| Pfdn6 | 1.12458383322973e-06 | -0.273401390868211 | 0.365 | 0.568 | 0.0349217017732829 |
| Mrps18c | 1.19039770200489e-06 | -0.255902449441209 | 0.18 | 0.348 | 0.0369654198403579 |
| Cox6c | 1.29727389100755e-06 | -0.289615764202188 | 0.556 | 0.725 | 0.0402842461374574 |
| Higd2a | 1.38464814242516e-06 | -0.262826424391507 | 0.276 | 0.46 | 0.0429974787667284 |
| Rbm5 | 1.57840773569288e-06 | -0.284243413700537 | 0.166 | 0.317 | 0.0490142954164711 |
| H2afy | 1.88082662186057e-06 | -0.268464358203085 | 0.276 | 0.46 | 0.0584053090886363 |
| Siglecf | 1.94024435264069e-06 | -0.285312743537338 | 0.5 | 0.672 | 0.0602504078825515 |
| Sdhb | 1.96914393160291e-06 | -0.268160892738531 | 0.222 | 0.401 | 0.0611478265080651 |
| Ppp1cc | 2.06941277576165e-06 | -0.254256626094098 | 0.317 | 0.505 | 0.0642614749257265 |
| Atp5j | 2.32124260187041e-06 | -0.250497556004211 | 0.651 | 0.791 | 0.072081546515882 |
| Ccl3 | 2.36514373386009e-06 | 0.618132284996139 | 0.857 | 0.822 | 0.0734448083675573 |
| Cox7c | 2.53446729501875e-06 | -0.343519032059558 | 0.376 | 0.54 | 0.0787028129122172 |
| Dnajc19 | 2.91036197695311e-06 | -0.262565439411828 | 0.143 | 0.279 | 0.090375470470325 |
| Ccng1 | 3.11833277500391e-06 | -0.287874509461244 | 0.253 | 0.425 | 0.0968335876621966 |
| Cnbp | 3.26501478415679e-06 | -0.257653519603008 | 0.649 | 0.826 | 0.101388504092421 |
| Cdk11b | 3.30920381359556e-06 | -0.256049805613298 | 0.351 | 0.537 | 0.102760706023583 |
| Tex261 | 3.49043269365522e-06 | -0.287313248070695 | 0.189 | 0.338 | 0.108388406436075 |
| Rbm42 | 4.09778790012012e-06 | -0.261292172563005 | 0.28 | 0.467 | 0.12724860766243 |
| Npm1 | 4.10464694526457e-06 | -0.261501970472438 | 0.309 | 0.509 | 0.127461601591301 |

| | | | | | |
|---|---|---|---|---|---|
| Uqcrb | 4.31568505518056e-06 | -0.262616110129415 | 0.338 | 0.53 | 0.134014968018522 |
| Spcs1 | 4.44688181532958e-06 | -0.254765026749128 | 0.28 | 0.463 | 0.138089021011429 |
| Sri | 4.82601374331356e-06 | -0.266909956742117 | 0.23 | 0.394 | 0.149862204771116 |
| Atp5g1 | 4.9150820259713e-06 | -0.266110877306042 | 0.376 | 0.561 | 0.152628042152487 |
| Lilr4b | 5.00354802971813e-06 | 0.319965822511782 | 0.811 | 0.732 | 0.155375176966837 |
| Gpd1l | 6.01382591570124e-06 | -0.251764466317664 | 0.089 | 0.206 | 0.186747336160271 |
| Clec4d | 6.29066365893112e-06 | 0.285599150622511 | 0.929 | 0.871 | 0.195343978600788 |
| Uqcr10 | 8.40685351361737e-06 | -0.27953582165788 | 0.402 | 0.582 | 0.26105802215836 |
| Cox7b | 1.41050594227361e-05 | -0.266096843732854 | 0.523 | 0.721 | 0.438004410254226 |
| Tmco1 | 1.43380025433053e-05 | -0.264869922734146 | 0.27 | 0.429 | 0.445237992977261 |
| H1f0 | 1.52720801085601e-05 | -0.431177125154129 | 0.041 | 0.125 | 0.474243903611118 |
| Rpl24 | 1.56331783255912e-05 | -0.258357674932365 | 0.705 | 0.854 | 0.485457086544584 |
| Pdcd6 | 1.58732251852729e-05 | -0.259136078209953 | 0.361 | 0.537 | 0.492911261678279 |
| Dnajc8 | 1.65427116925708e-05 | -0.265111329814675 | 0.344 | 0.519 | 0.5137008261894 |
| Ndufb9 | 1.71978756248733e-05 | -0.258295499272792 | 0.541 | 0.728 | 0.534045631779192 |
| Zc3h15 | 1.81932730965924e-05 | -0.256235229057798 | 0.263 | 0.422 | 0.564955709468482 |
| Lsm7 | 1.84882074931611e-05 | -0.256179930243267 | 0.147 | 0.279 | 0.574114307285131 |
| Ltc4s | 1.98067238830614e-05 | -0.29115985233158 | 0.195 | 0.345 | 0.615058196740705 |
| Nsrp1 | 2.15033565941042e-05 | -0.313970642396172 | 0.201 | 0.341 | 0.667743732316716 |
| Ptpn18 | 2.68781540370274e-05 | -0.25471637066658 | 0.407 | 0.582 | 0.83464731731181 |
| Cytip | 2.72021403458391e-05 | -0.276157866223366 | 0.521 | 0.693 | 0.84470806415934 |
| Basp1 | 2.80946064137801e-05 | 0.423431967534841 | 0.776 | 0.728 | 0.872421812967113 |
| Rgs2 | 2.82403536938376e-05 | -0.334392467813993 | 0.311 | 0.467 | 0.876947703254738 |
| Ccrl2 | 3.02692785647788e-05 | 0.283803178852424 | 0.954 | 0.927 | 0.939951907272076 |
| Odc1 | 3.16223440945563e-05 | -0.353172426124017 | 0.317 | 0.481 | 0.981968651168257 |
| B3gnt5 | 3.3459825833291e-05 | -0.270685105521905 | 0.124 | 0.244 | 1 |
| Il23a | 4.24836038754166e-05 | 0.662596196329502 | 0.415 | 0.303 | 1 |
| Eif3k | 5.02620959946942e-05 | -0.257074577743837 | 0.38 | 0.568 | 1 |
| Il1a | 5.29245891199274e-05 | 0.65053695692484 | 0.6 | 0.498 | 1 |
| Cflar | 5.49026613993243e-05 | 0.29262099006742 | 0.753 | 0.693 | 1 |
| Isy1 | 6.98183179342981e-05 | -0.255131485326043 | 0.276 | 0.425 | 1 |
| Rasgef1b | 0.000113369681550018 | -0.339785941718351 | 0.12 | 0.226 | 1 |
| Psmd11 | 0.000122124337532743 | -0.252156273172277 | 0.234 | 0.366 | 1 |
| Maff | 0.000159651792670214 | 0.258291297569406 | 0.759 | 0.686 | 1 |
| Ctsz | 0.000163223754367209 | 0.255078950349217 | 0.813 | 0.756 | 1 |
| Mt1 | 0.000163829246979454 | -0.417210276805163 | 0.232 | 0.359 | 1 |
| Emsy | 0.000169415412289658 | -0.272835552535281 | 0.095 | 0.188 | 1 |
| Tagln2 | 0.000186129174881076 | -0.265124290504693 | 0.427 | 0.568 | 1 |
| Ier3 | 0.00019667234266992 | 0.368886013174468 | 0.886 | 0.868 | 1 |
| Slc16a3 | 0.000217303514194879 | -0.278998191582845 | 0.577 | 0.718 | 1 |
| Hdc | 0.00046939303396848 | 0.261800047244352 | 0.766 | 0.728 | 1 |
| Mapk6 | 0.000479005645922879 | 0.282078189889182 | 0.608 | 0.54 | 1 |
| Stap1 | 0.0010321261772854 | -0.292760507668239 | 0.141 | 0.233 | 1 |
| Upp1 | 0.00143902761163716 | 0.386628037248905 | 0.234 | 0.15 | 1 |
| Ifitm1 | 0.00297524494160608 | 0.359364028478562 | 0.751 | 0.7 | 1 |
| Vegfa | 0.00569236054976958 | -0.308669841490757 | 0.278 | 0.38 | 1 |
| Ltb4r1 | 0.00709319643688475 | 0.25917774944788 | 0.305 | 0.233 | 1 |
| Rhov | 0.00778762483873459 | 0.252051237593856 | 0.147 | 0.087 | 1 |

| | | | | | |
|---|---|---|---|---|---|
| Creb1 | 0.010171725224693 | 0.259314045427999 | 0.38 | 0.324 | 1 |
| Slc11a1 | 0.0103344510759415 | 0.253742901297471 | 0.386 | 0.338 | 1 |
| Cxcl3 | 0.0307865290894197 | 0.64707128648919 | 0.438 | 0.397 | 1 |
| C1qa | 0.0549855225106923 | 0.287451795546472 | 0.32 | 0.293 | 1 |
| Gm43936 | 0.101320523167155 | 0.260065012767514 | 0.158 | 0.122 | 1 |
| Bcl2l11 | 0.104383000335771 | 0.280599106346698 | 0.674 | 0.686 | 1 |
| Lcn2 | 0.12151186531841 | -0.298509992531839 | 0.189 | 0.237 | 1 |
| 1110002J07Rik | 0.17582798954825 | 0.305681532107355 | 0.251 | 0.22 | 1 |
| Ccl4 | 0.200667693729462 | 0.588298772387979 | 0.583 | 0.613 | 1 |
| Il1f9 | 0.208418745981514 | 0.28753653677014 | 0.286 | 0.258 | 1 |
| Chka | 0.216571567847054 | 0.323719381876702 | 0.295 | 0.282 | 1 |
| Gm5483 | 0.230175754894127 | 0.461438353785447 | 0.187 | 0.164 | 1 |
| Ccl6 | 0.325792863829744 | 0.255432830993254 | 0.402 | 0.408 | 1 |
| Cd83 | 0.360452781072358 | -0.362976335361099 | 0.12 | 0.143 | 1 |
| Ccl9 | 0.375560847343684 | -0.336925480353332 | 0.197 | 0.23 | 1 |
| Malt1 | 0.397228344722694 | 0.299512737065734 | 0.494 | 0.537 | 1 |
| Cxcl1 | 0.612986922793236 | 0.523367843703141 | 0.28 | 0.275 | 1 |
| Tnfsf9 | 0.649580793581171 | 0.29284570269915 | 0.237 | 0.23 | 1 |
